# Benchmarking within-sample minority variant detection with short-read sequencing in *M. tuberculosis*

**DOI:** 10.64898/2026.02.13.704885

**Authors:** Shandukani Mulaudzi, Sanjana Kulkarni, Maximillian G. Marin, Maha Farhat

## Abstract

**Motivation:** Low-frequency (minority) variants—variants detectable within-sample at low allele frequencies—are relevant in several areas of research and health, from cancer to pathogen heteroresistance. There is uncertainty around the optimal bioinformatic approach to accurately and reproducibly distinguish low-frequency variants from sequencing or mapping errors. To address this, we benchmarked seven variant callers on precision, recall, and false positive characteristics for detecting low-frequency variants using simulated short-read whole-genome sequencing data for 700 *Mycobacterium tuberculosis* strains. We developed a new low-frequency error model for filtering the output of the best-performing tool using read mapping and quality metrics.

**Results:** We simulated 378 unique variants across five genomic backgrounds spanning four lineages. Variants were simulated to represent three genomic region categories, 10 allele frequencies and five sequencing depths. FreeBayes, a haplotype-based variant caller, achieved the highest pooled F1 score of the seven tools in drug resistance regions (average F1=0.86) and its higher performance held across genomic context and background. Across tools, we identified lower performance in repetitive (low mappability) regions, and strong reference bias in low-frequency variant calling. We validated variant caller performance on *in-vitro* strain mixtures substantiating our ranking. The error model excludes <1% of true variants identified by FreeBayes on average per strain, and excludes 97% of FPs on average per strain when combined with low mappability region and rRNA gene masking. Our analysis informs best practices for low-frequency variant calling, including tool choice, masking, and filtering. We also provide a new error model that excludes false low-frequency variant calls from FreeBayes output.

**Availability and implementation:** All relevant code is available at https://github.com/shandu-m/benchmark-minority-variants-Mtb.

**Supplementary Information:** Supplementary Files 1-3 respectively are attached with this submission.

## Introduction

Bulk whole-genome sequencing (WGS) is usually applied to study genetic variation that differentiates distinct cell populations. However, bulk WGS can also capture recently-evolved diversity emerging within a single population. The latter requires the accurate detection of low-frequency (minority) or sub-consensus variants present at mapped read frequencies typically <75-95%. Low-frequency variants are relevant for understanding within-host pathogen evolution, somatic oncogenesis in cancers, and microbiome diversity, among other applications. Specifically for *Mycobacterium tuberculosis* (*Mtb*), the causative agent of tuberculosis (TB), patient-derived *Mtb* isolates from bacterial culture are not pure clones (Lieberman *et al*. 2011, 2014, Marvig *et al*. 2015, Copin *et al*. 2016, Didelot *et al*. 2016). Bulk *Mtb* whole-genome sequencing at medium-to-high depth suggests that up to 85% of variant calls per sample occur at mapped read frequencies of less than 50% (Sun *et al*. 2012, Black *et al*. 2015, O’Neill, Mortimer, and Pepperell 2015, Casali *et al*. 2016, Copin *et al*. 2016, Trauner *et al*. 2017). This variation may have been present at infection (e.g. multi-strain primary infections) or may have arisen during reinfection with additional *Mtb* strains. Low-frequency variation can also arise during the course of chronic clonal infection when subpopulations of bacteria acquire new variants in-host (Ford *et al*. 2012, Guerra-Assunção *et al*. 2015, Lieberman *et al*. 2016, Vos *et al*. 2019). Some of these low-frequency variants have been shown to rise to fixation in-host, and within-population frequencies of 19% or more have been shown to predict subsequent fixation in-host (Eldholm *et al*. 2014, Engelthaler *et al*. 2019, Polsfuss *et al*. 2019, Vos *et al*. 2019, Andersson, Nicoloff, and Hjort 2019, Vargas, Freschi, Marin *et al*. 2021). Low-frequency variants therefore present an opportunity for early diagnosis of drug resistance (DR) in *Mtb* or early detection of poor TB treatment response, as has been shown previously for *Mtb* and other pathogens (Izopet *et al*. 2002, Cohen *et al*. 2016, Houldcroft *et al*. 2016).

Although whole-genome sequencing enables the detection of low-frequency variants, such variants are hard to distinguish from base level sequencing error especially at lower depths, or from mapping error due to sequence homology or reference bias. Previous *Mtb* WGS studies have employed several variant calling approaches to distinguish between true and false low-frequency variants including: (1) hard filtering variant calls on quality metrics including depth, allele frequency, base quality, strand bias (Sun *et al*. 2012, Chen *et al*. 2021, Nimmo *et al*. 2019, Lee *et al*. 2020, Martin *et al*. 2018, Vargas, Freschi, Marin *et al*. 2021, Vargas, Freschi, Spitaleri *et al*. 2021, Vargas *et al*. 2023), (2) variant calling against a more closely matched reference genome (Lee *et al*. 2020), (3) excluding variants in repetitive regions (Casali *et al*. 2016, Copin *et al*. 2016, Trauner *et al*. 2017, Nimmo *et al*. 2019, 2020), (4) empiric comparison to data from deep targeted sequencing (Goossens *et al*. 2022), and (5) taking a consensus of variant calls across two or more tools (Liu *et al*. 2015, Trauner *et al*. 2017). Despite the empiric application of these different bioinformatic approaches, data guiding the choice of variant calling tool is currently lacking and this significantly limits the interpretation of the results of previous analyses. In this *Mtb*-focused benchmarking analysis we explicitly consider the effects of repetitive sequence and homopolymeric context as well as reference bias on low-frequency variant calling, surpassing existing benchmarking efforts on minority variants in gut microbiome, viral and human tumor data (Andreu-Sánchez *et al*. 2021, Said Mohammed *et al*. 2018, Fang *et al*. 2015, Bohnert, Vivas, and Jansen 2017, Spencer *et al*. 2014, Bian *et al*. 2018, Maruzani *et al*. 2024, Sergi *et al*. 2024).

We assess the performance of seven tools on simulated data, providing one of the most comprehensive benchmarking studies to date for low-frequency within-sample variant calling from bulk sequence data. Tools were selected either because they were specifically developed for *Mtb* low-frequency variant detection (binoSNP; Dreyer *et al*. 2020) or general microbial variant detection (Pilon; Walker *et al*. 2014), are commonly applied for *Mtb* low-frequency variant calling (LoFreq, VarScan2; Koboldt *et al*. 2012; Wilm *et al*. 2012), or because they are highly cited methods for somatic mutation calling in human data (FreeBayes, Mutect2, VarDict; Garrison and Marth 2012; Lai *et al*. 2016; Benjamin *et al*. 2019). Specifically we benchmark: (1) FreeBayes: a haplotype-based bayesian caller used for SNV and small indel genotyping in the second edition of the WHO catalog of MTBC DR mutations (WHO 2023), (2) LoFreq: an ultra-sensitive variant caller for uncovering cell-population heterogeneity, (3) Mutect2: GATK’s recommended variant caller for somatic short variant discovery, (4) Pilon: an integrated tool for microbial variant calling and genome assembly, (5) VarDict: developed for next-generation sequencing in cancer research, and previously benchmarked against FreeBayes and other variant callers for bacterial WGS data (Seah *et al*. 2023), (6) VarScan2: a tool for somatic mutation and copy number alteration discovery in cancer, and (7) BinoSNP: developed for low-frequency detection of single nucleotide variants (SNVs) in *Mtb* complex (MTBC) strains. We simulate WGS data using two different approaches (InSilicoSeq and ART; Huang *et al*. 2012; Gourlé *et al*. 2019). Each simulated *Mtb* strain has a distinct combination of 1) mutant background 2) simulated depth, 3) mutant allele frequency and 4) reference genome. The simulations are replicated five times for a total of 7000 WGS simulations.

## Results

### Simulations

We benchmarked seven variant callers for low-frequency variant detection in *Mtb* using a total of 500 *Mtb* strains simulated from an H37Rv background and 200 *Mtb* strains simulated from a non-H37Rv lineage 1-4 background. Each H37Rv strain had 50 simulated mutations across drug resistance (DR), homopolymer tract (HT), and low mappability (LM) regions, and each L1-4 strain had 20 simulated DR mutations. Strains were simulated in five replicates by both InSilicoSeq (ISS) and ART for a total of 7000 simulated strains (Methods).

Across genome backgrounds (H37Rv, L1-4), the simulators successfully produced the target depths and all simulated bases had a quality score above 35 (Figure S15). The simulated allele frequencies (AFs) show high fidelity to the expected AFs for both simulators and more details are provided in the supplement (Figure S16; Supplementary Results A). The ART-simulated variants demonstrated slightly higher AF variance than the ISS-simulated variants (standard error 0.049% vs 0.031%). All results presented are based on the ISS simulation data. The ART data resulted in similar conclusions and is provided in Supplementary Results E.

When comparing tools, we assessed accuracy as the weighted F1 score pooled over simulated variant AF, sequencing depth and type of region (DR, HT or LM; see Methods) for the H37Rv simulations. Accuracy by genomic region was pooled over variant AF and sequencing depth, and accuracy as a function of minimum variant AF was pooled over sequencing depth. BinoSNP is not included in the final tool performance analysis due to high time complexity (see Supplementary Results B).

### FreeBayes achieves the highest overall accuracy

FreeBayes demonstrates the highest average weighted H37Rv F1 score over all mutation types (mean = 0.87, Mann-Whitney P-value = 5.12E-06 for comparison to VarDict; Fig. 1a). VarDict has the next highest weighted F1 score of 0.80, and Pilon achieves the lowest weighted F1 score of 0.58. Across the five genomic backgrounds, all tools except Pilon achieve an average DR-F1 ≥ 0.71 (Pilon average DR-F1 = 0.57; Table S3).

**Fig. 1.**
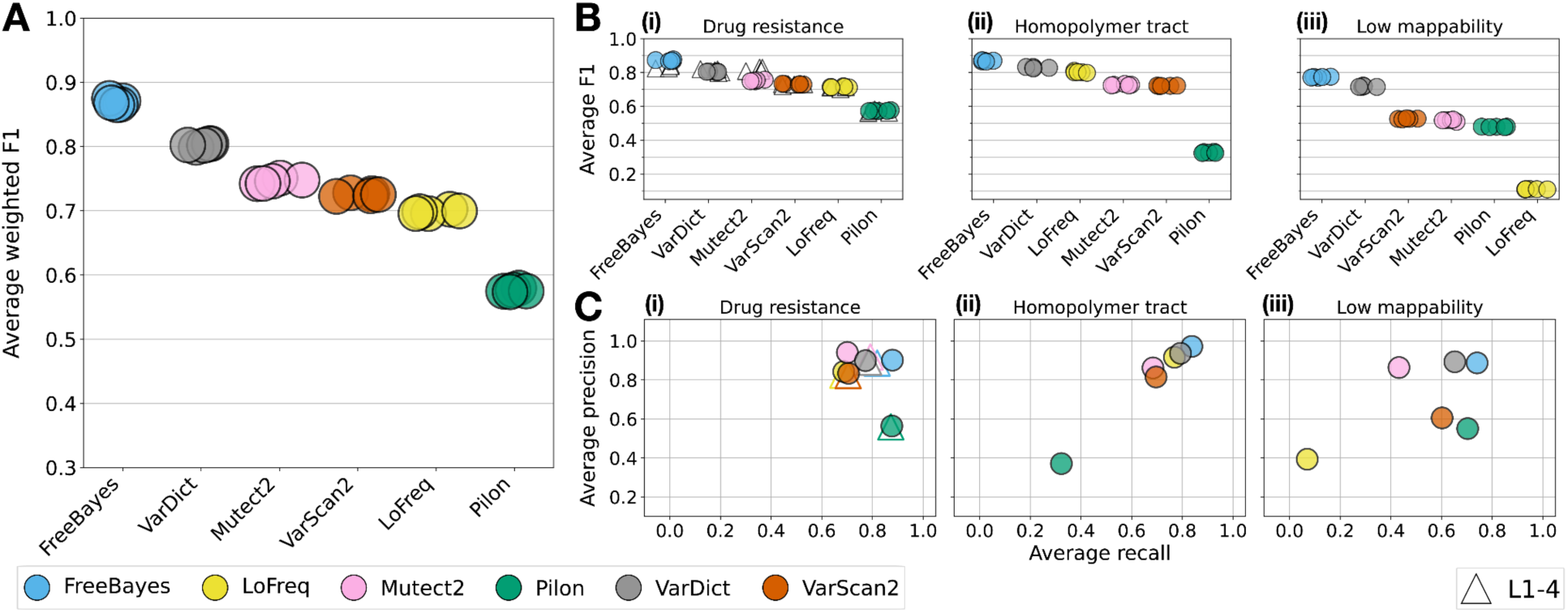
Overall variant caller accuracy. **a** Weighted F1 score achieved by each variant caller in the H37Rv strains, averaged over simulated variant AFs, depths and mutation regions. Each point represents the average weighted F1 score for one of the replicate simulations. **b** Average F1 score achieved by each variant caller in (i) drug resistance, (ii) homopolymer tract, and (iii) low mappability regions. Subplot (i) includes both the H37Rv simulations (colorful circles) and L1-4 simulations (white triangles). **c** Average precision and recall achieved by each variant caller in (i) drug resistance, (ii) homopolymer tract, and (iii) low mappability regions. Subplot (i) includes both the H37Rv simulations (colorful circles) and L1-4 simulations (white triangles with edge colors corresponding to each tool). Note: Pilon does not report INDELs at an AF < 25% by default.

### Low-frequency variant calling accuracy varies by genomic region and is highly uncertain in low mappability regions

Across the three genomic regions, accuracy is lowest and most variable for LM compared with DR or HT regions for all tools except Pilon (LM-F1 is on average 0.22 lower than DR-F1 in the H37Rv samples) (Fig. 1b). FreeBayes achieves either the highest or second-highest precision and recall in each region (Fig. 1c, Tables 1a-c). Tools with the highest overall accuracy demonstrate the lowest variance in performance across the mutation types (DR, HT, LM).

In DR regions, all variant callers achieve comparable H37Rv and L1-4 F1 scores (Fig. 1b(i), Table S3), and the DR-F1 score tool ranking is consistent with the weighted H37Rv F1 score pooled across regions. For each of LoFreq, Mutect2, Pilon and VarScan2, the average DR F1 scores do not differ significantly across the four L1-4 backgrounds (all P-values > 0.05; Figure S17). VarDict and FreeBayes exhibit statistically significant differences in the average F1 score for L3 and every other lineage (P-values ≤ 0.05). The F1 in L3 samples is 0.018 and 0.043 lower than the average F1 across lineages 1, 2 and 4 for FreeBayes and VarDict respectively (see Table S4 and Supplementary Results C).

**Table 1a.** Average drug resistance precision and recall in the strains simulated from an H37Rv and L1-4 background.

| Drug resistance regions |  |  |  |  |  |  |  |  |
| --- | --- | --- | --- | --- | --- | --- | --- | --- |
| Rank | Precision |  |  |  | Recall |  |  |  |
|  | H37Rv |  | L1-4 |  | H37Rv |  | L1-4 |  |
| 1 | Mutect2 | 0.941** | Mutect2 | 0.918** | FreeBayes | 0.879 | Pilon | 0.873** |
| 2 | FreeBayes | 0.902** | VarDict | 0.891** | Pilon | 0.876** | FreeBayes | 0.819** |
| 3 | VarDict | 0.899** | FreeBayes | 0.886** | VarDict | 0.772 | Mutect2 | 0.791** |
| 4 | LoFreq | 0.843 | LoFreq | 0.824 | VarScan2 | 0.706** | VarDict | 0.785 |
| 5 | VarScan2 | 0.834** | VarScan2 | 0.822** | Mutect2 | 0.701** | VarScan2 | 0.704 |
| 6 | Pilon | 0.564 | Pilon | 0.559 | LoFreq | 0.687 | LoFreq | 0.683 |

Asterisks indicate the precision and recall rankings that are statistically significant after Benjamini-Hochberg correction (Mann-Whitney U test with FDR = 0.05).

Tools are ranked according to precision and recall separately for each background genome group. Mann-Whitney U tests were performed for each sequential pair of tools, e.g. in the first row of the precision section of the table we tested for a significant difference in the distribution of precision values between the first and second ranking tools.

**Table 1b.** Average homopolymer tract precision and recall in the strains simulated from an H37Rv background.

| Homopolymer tract regions |  |  |  |  |
| --- | --- | --- | --- | --- |
| Rank |  | Precision |  | Recall |
| 1 | FreeBayes | 0.972** | FreeBayes | 0.837** |
| 2 | VarDict | 0.937** | VarDict | 0.793 |
| 3 | LoFreq | 0.916** | LoFreq | 0.769** |
| 4 | Mutect2 | 0.862** | VarScan2 | 0.696** |
| 5 | VarScan2 | 0.815** | Mutect2 | 0.684** |
| 6 | Pilon | 0.371 | Pilon | 0.322 |

Asterisks indicate the precision and recall rankings that are statistically significant after Benjamini-Hochberg correction (Mann-Whitney U test with FDR = 0.05).

Tools are ranked according to precision and recall separately.

**Table 1c.** Average low mappability precision and recall in the strains simulated from an H37Rv background.

| Low mappability regions |  |  |  |  |
| --- | --- | --- | --- | --- |
| Rank |  | Precision |  | Recall |
| 1 | VarDict | 0.893** | FreeBayes | 0.740** |
| 2 | FreeBayes | 0.887** | Pilon | 0.703** |
| 3 | Mutect2 | 0.864** | VarDict | 0.652** |
| 4 | VarScan2 | 0.605 | VarScan2 | 0.602** |
| 5 | Pilon | 0.551** | Mutect2 | 0.432** |
| 6 | LoFreq | 0.394 | LoFreq | 0.070 |

Asterisks indicate the precision and recall rankings that are statistically significant after Benjamini-Hochberg correction (Mann-Whitney U test with FDR = 0.05).

Tools are ranked according to precision and recall separately.

### Tools demonstrate more than 95% accuracy at AF ≥ 10% outside of low mappability regions

We assessed how the six tools vary in their accuracy as a function of variant AF and sequencing depth. We focused on depths 50-200x as these depths are common in clinical *Mtb* sequencing, and because detection power of low AFs expectedly increases at depths > 200x where tool accuracy also converges (Figure S18). Tool rankings are consistent based on F1 score pooled across all depths and genomic backgrounds, where the average DR F1 > 0.92 for all tools at a variant AF ≥ 5% (Figure S19).

At depths 50-200x and variant AF ≥ 10%, all tools have equivalent cumulative accuracy across DR and HT regions (F1 ≥ 0.95, Fig. 2, Table S5), with the exception of Pilon in HT regions (F1 = 0.69; Pilon does not report INDELs at AF < 25% by default). In LM regions, while FreeBayes, Pilon and VarDict achieve an F1 ≥ 0.93 at variant AF ≥ 10%, VarScan2 and Mutect2 only achieve comparable F1 at variant AFs of 20% and 50% respectively. LoFreq performs poorly in LM regions, achieving an F1 < 0.50 even for variants AF ≥ 50%. Tool rankings are similar for variant calling at AF ≥ 10% and AF < 10% but across the AF range examined, tool accuracy differed the most at lower AFs (AF 1-5%, Fig. 1, Figure S20).

**Fig. 2.**
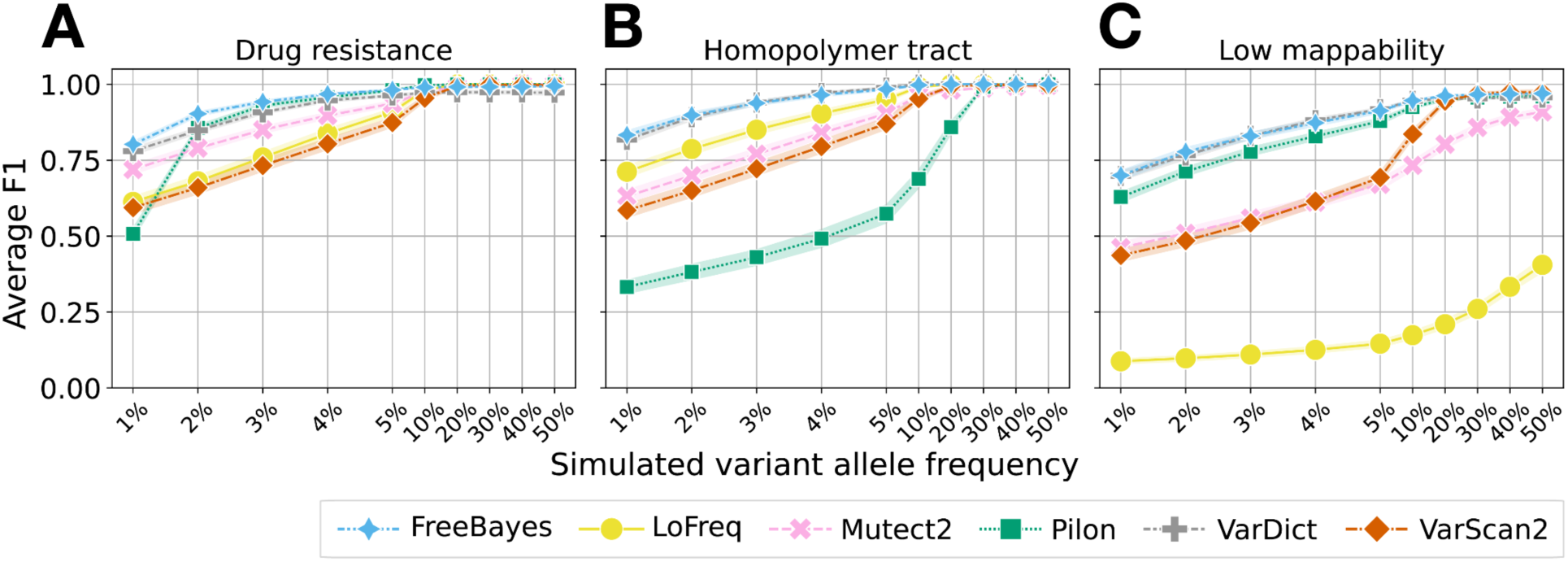
Average cumulative F1 across variant AF pooled over depths of 50x, 100x and 200x. **a** Average cumulative F1 in drug resistance regions (H37Rv and L1-4 strains). **b** Average cumulative F1 in homopolymer tract regions (H37Rv strains only). **c** Average cumulative F1 in low mappability regions (H37Rv strains only). To compute the average cumulative F1, we computed cumulative precision and recall as a function of increasing minimum variant AF for each of the six tools, averaged over haplotype, depths 50-200x and replicate. The band around each line represents the 95% confidence interval. Note that the x-axis tick gaps are not proportional to the actual simulated variant AF, and are larger for AF < 10% as this is where the greatest tool-wise differences occur.

### Low mappability regions are prone to false positives with high allele frequency

We next determined the number of false positives (FP) detected by each tool by region type and sequencing depth. We expanded the FP assessment to consider LM regions more comprehensively than regions in which we simulated LM low-frequency variants (Supplementary Methods K; “LM” from this point onwards refers to these more comprehensive LM regions). The benchmarking showed strong reference bias, with a false positive rate (FPR) at least fivefold higher in L1-4 than in H37Rv strains for all tools except Pilon (P-values ≤ 0.05; median genome-wide FPR for all tools >7E-05 in L1-4 strains and <1E-04 in H37Rv strains; Figure S21). The two variant callers developed specifically for low-frequency variants, LoFreq and Mutect2, detect the fewest FP per strain (the lowest FPR of 2.90E-05 in an L1-4 strain is achieved by Mutect2). A median of 96% of FPs are SNPs across all tools and genome backgrounds (Figure S22).

In the L1-4 and H37Rv strains, the FPR is highest in LM regions (median FPR in LM regions is 4.10E-06, versus 0 in DR regions and 1.54E-06 elsewhere in the genome–non-DR and non-LM; Figure S23, Table S6). The average AF for FP LM variants is also significantly higher than for DR or other regions (average AF for LM FPs is 7%, versus 1% for DR and other regions respectively; Fig. 3, Figure S24). Consequently, most of the FP detected in these strains are in LM regions, with the exception of Pilon which detects an excess of FPs at AF = 1% (Table S7). We note that the relationship between FPR and sequencing depth is tool-specific (Figure S6), and the differences in FPR by lineage background (L1-4) are small and detailed in Figure S25.

**Fig. 3.**
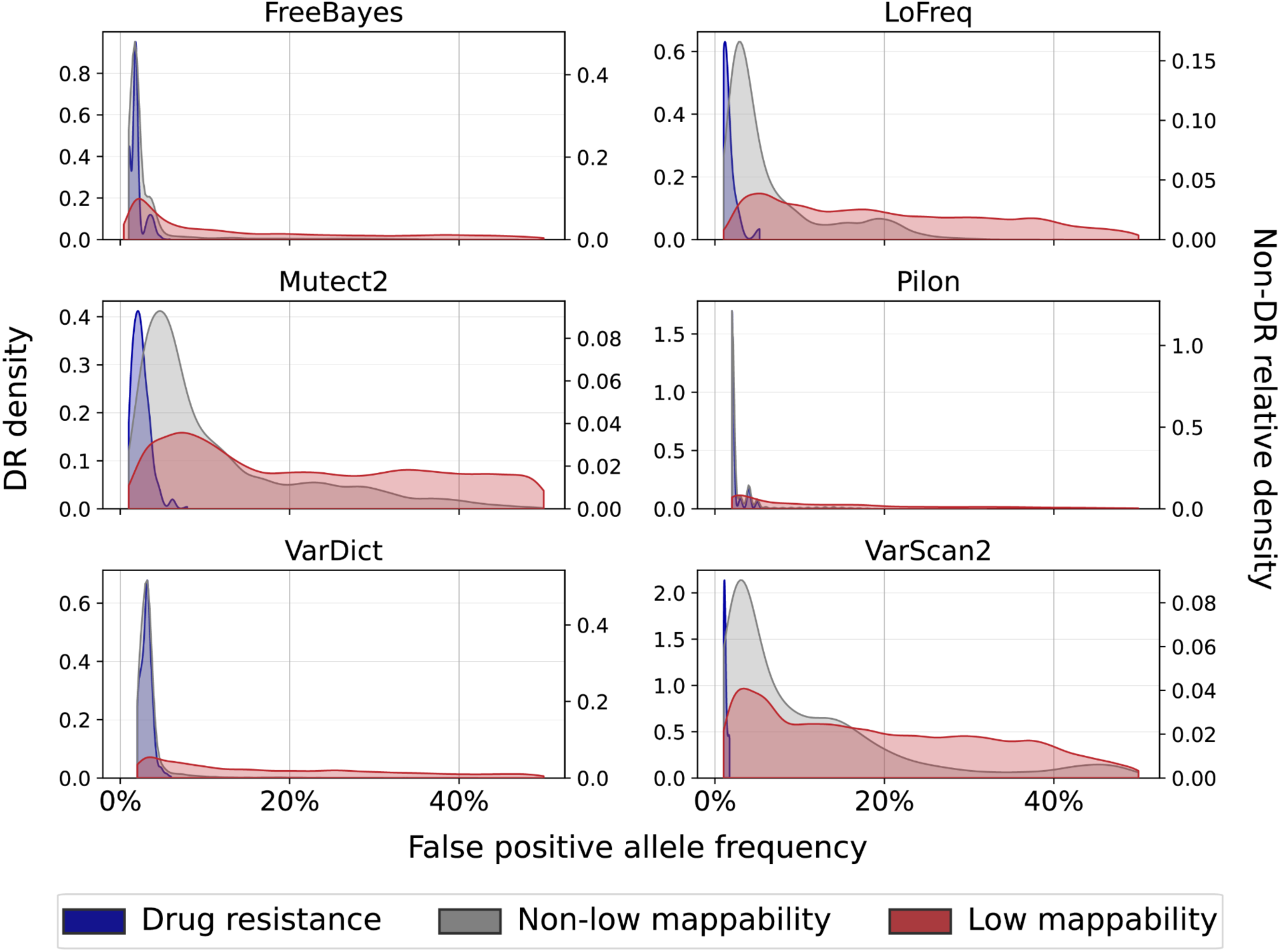
False positive allele frequency distribution in the L1-4 strains by region. Each region is defined to be mutually exclusive for this comparison i.e. the non-low mappability regions do not include the drug resistance regions. The left y-axis displays the densities of the DR AF distributions, and the right y-axis displays the relative densities of the low mappability (non-DR) and non-low mappability (non-DR) AF distributions (normalized independently). For Pilon we include only the FP with AF > 1% (a median of 42% of Pilon FPs across all strains have AF = 1%). All DR FP occur at AF < 8% and FP in the other two regions occur at AFs 1-50%. Between 2-38% of the FP in non-low mappability regions occur at AF > 10%, while more than 60% of the LM FPs occur at AF > 10% for all tools except FreeBayes and Pilon (47% and 50% of the LM FPs have AF > 10% for these tools respectively).

To better understand sources of FP variant calls we studied read mapping and quality characteristics at variant site positions in the L1-4 strains: including base and mapping quality, coverage relative to the average regional coverage (coverage ratio), the number of discordantly-aligned reads normalized to site coverage and the number of soft-clipped bases normalized to site coverage (Figures S26-30). Mapping quality, coverage ratio, discordantly-aligned reads ratio and soft-clipped bases ratios at a variant site differed significantly between FP and TP variants. There are smaller differences in base quality between FPs and TPs in the simulated data (Figure S15b).

### An error model to improve the specificity of low-frequency variant detection using FreeBayes

FreeBayes had the highest overall performance as measured by average weighted F1, and high performance at AF < 10%. However FreeBayes performance favors higher recall over slightly lower precision for some regions compared with other highly performing tools such as VarDict and Mutect2 (Tables 1a-c). We hypothesized that pairing a high-recall tool like FreeBayes with post-filtering can further improve the balance of recall and precision for low-frequency variant calling.

Closer examination of read mapping and quality characteristics at FreeBayes SNV call positions identified varied reasons for false positive low-frequency variant calls (Fig. 4). We hence used the read quality metrics to build a multivariate logistic error model that estimates the probability of a false call for any unfixed SNV call with AF 5-95%, at least 2 forward and reverse strand mapped reads, total depth ≥ 5, and MQ ≥ 40 (Methods).

**Fig. 4.**
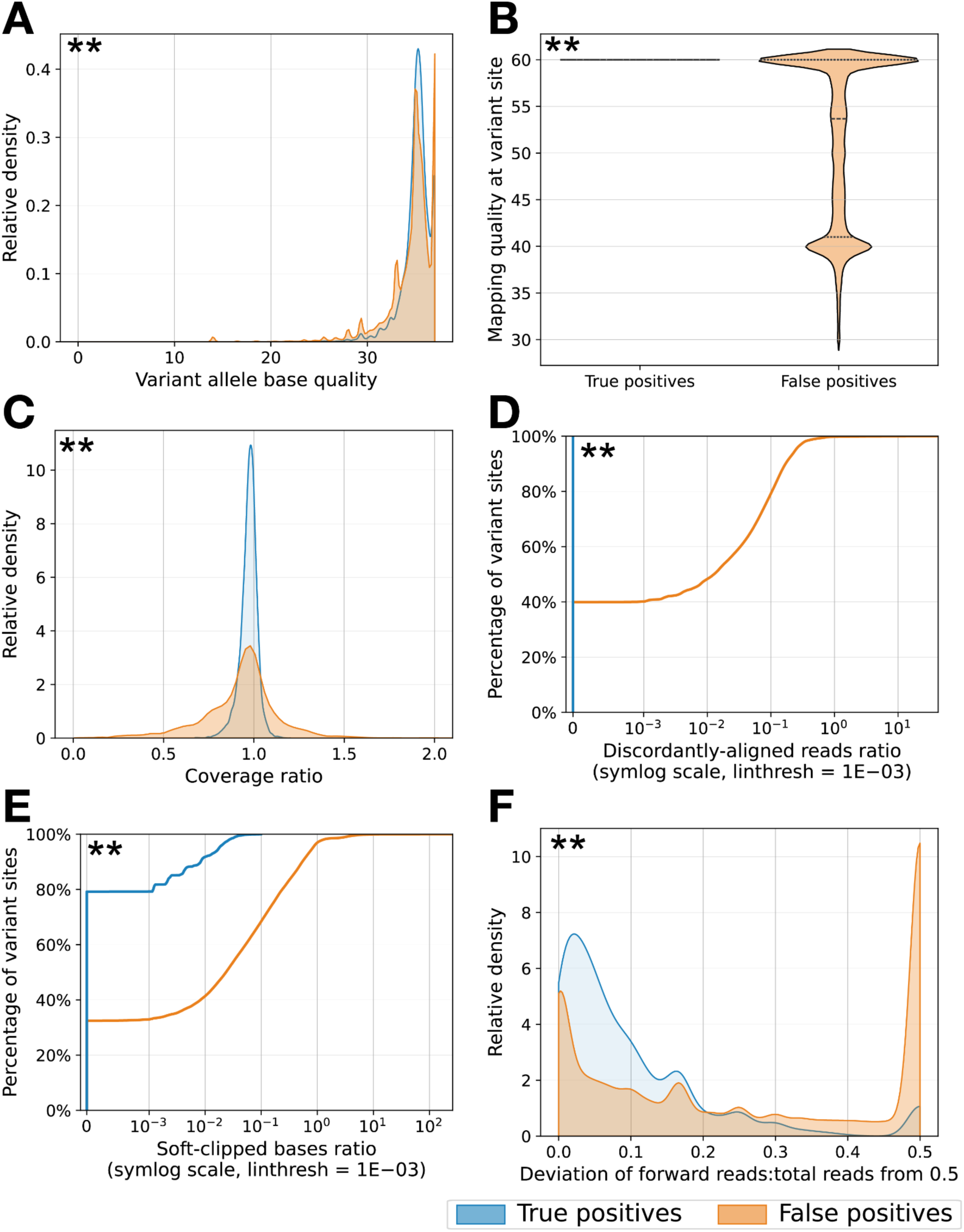
Read mapping and quality characteristics at FreeBayes SNV sites (AF ≤ 50%) split by true and false positives. **a** Base quality. **b** Mapping quality. **c** Coverage ratio: coverage at a variant site relative to the average regional coverage; x-axis is cut-off at 2; 0% of TP sites and 0.09% of FP sites have a coverage ratio > 2. **d** Discordantly-aligned reads ratio: number of discordantly-aligned reads at a variant site relative to site coverage; the empirical cumulative distribution function is shown on a symmetric logarithmic x-axis with a linear threshold of 1E-03; 100% of TP sites and 40% of FP sites have a discordantly-aligned reads ratio of 0. **e** Soft-clipped bases ratio: number of soft-clipped bases at a variant site relative to site coverage; the empirical cumulative distribution function is shown on a symmetric logarithmic x-axis with a linear threshold of 1E-03; 79% of TP sites and 32% of FP sites have a soft-clipped bases ratio of 0. **f** Strand bias: the magnitude of the deviation from 0.5 of the number of forward reads supporting a variant as a fraction of the total reads at the variant site. Mapping quality, coverage ratio, discordantly-aligned reads ratio and soft-clipped bases ratio at a variant site have differing characteristics for FP variant calls compared to TP variant calls. Strand bias and base quality have smaller differences between FPs and TPs. The differences between the TP and FP distributions are statistically significant for each metric after Benjamini-Hochberg correction (indicated by asterisks; Mann-Whitney U test with FDR = 0.05).

We tested this error model on the simulated L1-4 strains to filter FreeBayes SNV calls, focusing on the FP detected with comparable AF to the introduced variants (AF ≤ 50%). The error model excludes, on average, 49% of FreeBayes SNV FPs (AF 5-50%) detected in an L1-4 strain, and <1% of TPs on average per strain (F1 in Table 2). Error model filtering combined with masking low mappability regions and rRNA genes (F3) excludes 97.3% of FPs on average, and excludes 15.5% of TPs on average which correspond to the simulated variants in *rpoB* (n = 2) and *rrs* (n = 1). Error model filtering with region masking provides the most substantial gain in precision, while limiting the reduction in recall to a set of known regions, which can be more fairly adjusted for in downstream analyses. An average SNV FPR < 3E-6 is achieved per strain after error model filtering and region masking (F3), more than 30 times lower than the initial FPR. For variant AFs 5-50%, error model filtering with region masking is more effective at reducing FPs than for variant AFs < 5%, especially at lower sequencing coverages (Table S9).

To filter FP INDEL calls made by FreeBayes we built a pipeline to adjust the allele fraction for unfixed FreeBayes INDEL calls (AF 5-95%) and retain only INDEL calls with AF ≥ 5% post-adjustment (Methods). After allele fraction adjustments for the FreeBayes INDEL calls (AF 5-50%), 43% of the INDEL FPs are filtered out on average per strain for the L1-4 strains (Table S10). Hard filtering on read count thresholds and mapping quality and region masking (see Supplementary Methods N) filters out an average of 99% of the INDEL FPs. Only 6.5% of the total INDEL TPs detected at AF ≥ 5% in the strains simulated from an H37Rv background are lost on average with this filtering scheme, and this loss in sensitivity is more pronounced at lower sequencing coverages (Table S10).

**Table 2.**
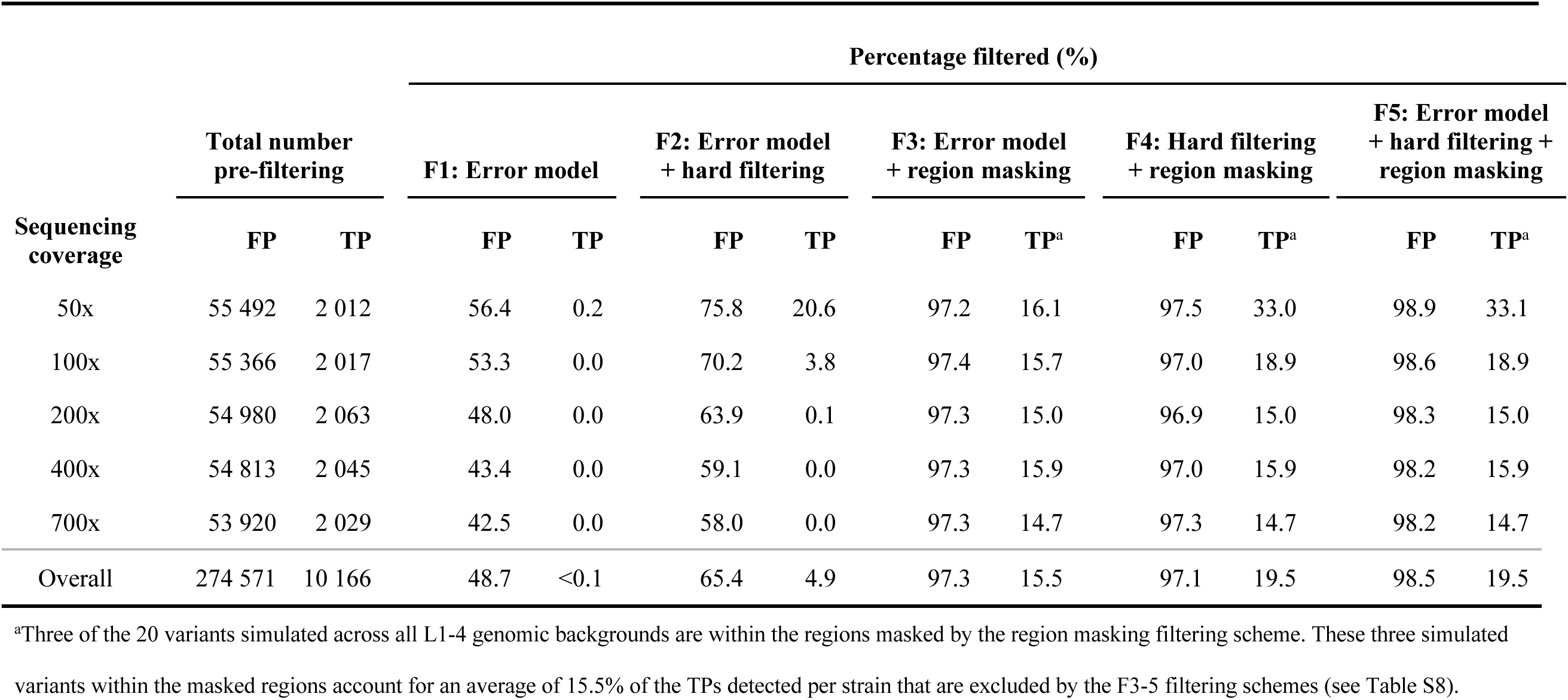
SNV filtering of false and true variants called by FreeBayes (AF 5-50%) in L1-4 strains.

The pre-filtering group of columns displays the total number of FP and TP SNVs called by FreeBayes across all strains simulated at a specific sequencing depth. A total of 120 strains are considered in each sequencing depth group (six variant AFs 5-50%, four background genomes, five replicates). The F1-F5 groups of columns correspond to each of the three sequential SNV filtering schemes and display the average percentages of FPs and TPs filtered out per strain. Error model filtering excludes variants with a model predicted probability ≤ 0.46; hard filtering excludes variants with forward and reverse strand allele counts < 2, depth < 5, mapping quality < 40); region masking excludes variants in low mappability regions and rRNA genes. FP and TP statistics are broken down by simulated sequencing coverage group, and summarized overall.

### Tools overestimate allele frequency on average

The variant AFs quantified by each tool differ only slightly from the simulated AF as a function of sequencing depth (AF median difference = 0.42% IQR 0-1.3% pooled across tools and depths, Figure S31). Variant callers are more likely to overestimate than to underestimate AF, likely due to the exclusion of low quality reads at variant sites which lowers the total depth captured by the variant caller at that site. The AF differences are smallest for LoFreq, VarDict and VarScan2 (median difference for all three tools is between 0-0.34%, Fig. 5, Figure S32). For all three tools, the AF differences of at least 92% of reported variants are within ±2%. FreeBayes, Mutect2 and Pilon have slightly larger differences (median differences are 0.52%, 1.2% and 1.0% respectively), still, 73%, 63% and 60% of the all variant AFs reported by each of these three tools are within ±2% of the simulated value. The larger AF differences for these three tools can be attributed to low-frequency variants in LM regions (Figure S33). The bias in AF measurement does not differ substantially by genomic background in DR regions (median AF difference is 0.24% and 0.26% respectively for H37Rv and L1-4).

**Fig. 5.**
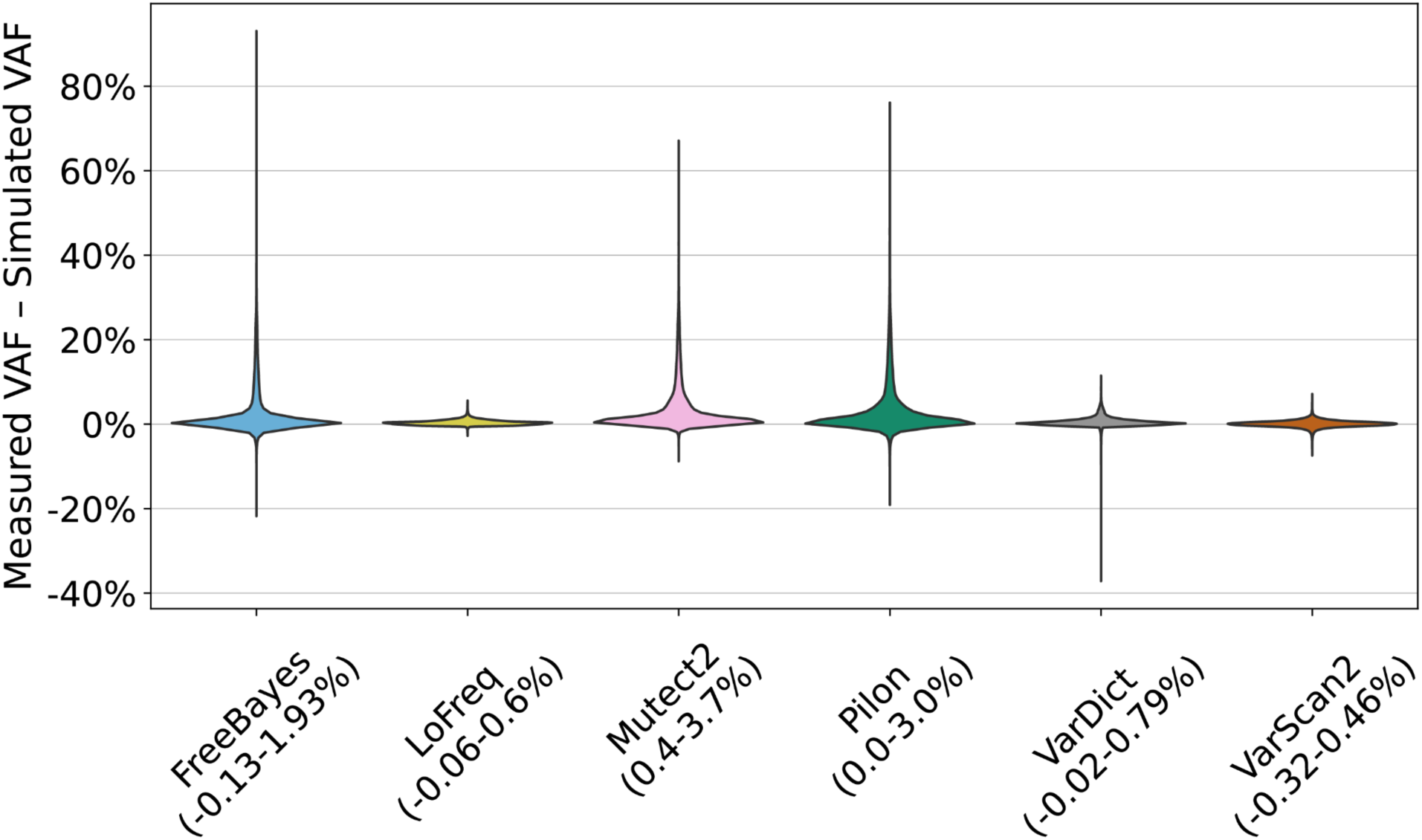
Distribution of the difference between the measured allele frequency and the simulated allele frequency. Each distribution includes these allele frequency differences for variants across all simulated variant allele frequencies, depths, genomic regions, genomic background and replicates. The IQR for each distribution is displayed next to the tool name. VarDict reported 18 variants with an AF at least 35% lower than the simulated AF, six of which were all within ±9 base pairs of each other in the same strain (H37Rv background, sequencing coverage 100x, simulated AF = 40%). All of these variants in the H37Rv strains are in rpoB, which overlaps with our definition of comprehensive LM regions, and 10 of these variants occurred at the same position in gyrA which was found to be problematic for FreeBayes due to its proximity to a baseline (fixed) lineage variant (see Supplementary Results C).

### Detection of low-frequency variants in experimental strain mixtures

We compared tool accuracy using sequencing data from six *in-vitro* strain mixtures. The wildtype H37Rv strain that did not harbor a mutation in *rpoB* (SR1a or SR4k) was mixed in proportion with its daughter colony harboring a single *rpoB* mutation (Ser531Leu and His526Pro respectively) generating a final *rpoB* mutation frequency of 1%, 5% or 10% (Dreyer *et al*. 2020).

Five of the six tools successfully detected variants present at AFs of 5% and 10% (Fig. 6a). The exception was VarDict which failed to produce results for one isolate after running for 3 days. Focusing on FPs (variants that were not known to occur in the daughter or wild type strains) at AF ≥ 5%, no FP are detected in *rpoB* by any tool. Outside of *rpoB* in the remaining DR regions, no FPs at AF ≥ 5% were detected except by LoFreq (3 FP across all isolates) and Pilon (42 FP across all isolates) (Fig. 6b). All of the FP detected by LoFreq had AF ≤ 7.5%, while Pilon’s detected FP ranged from AF 5-15%.

**Fig. 6.**
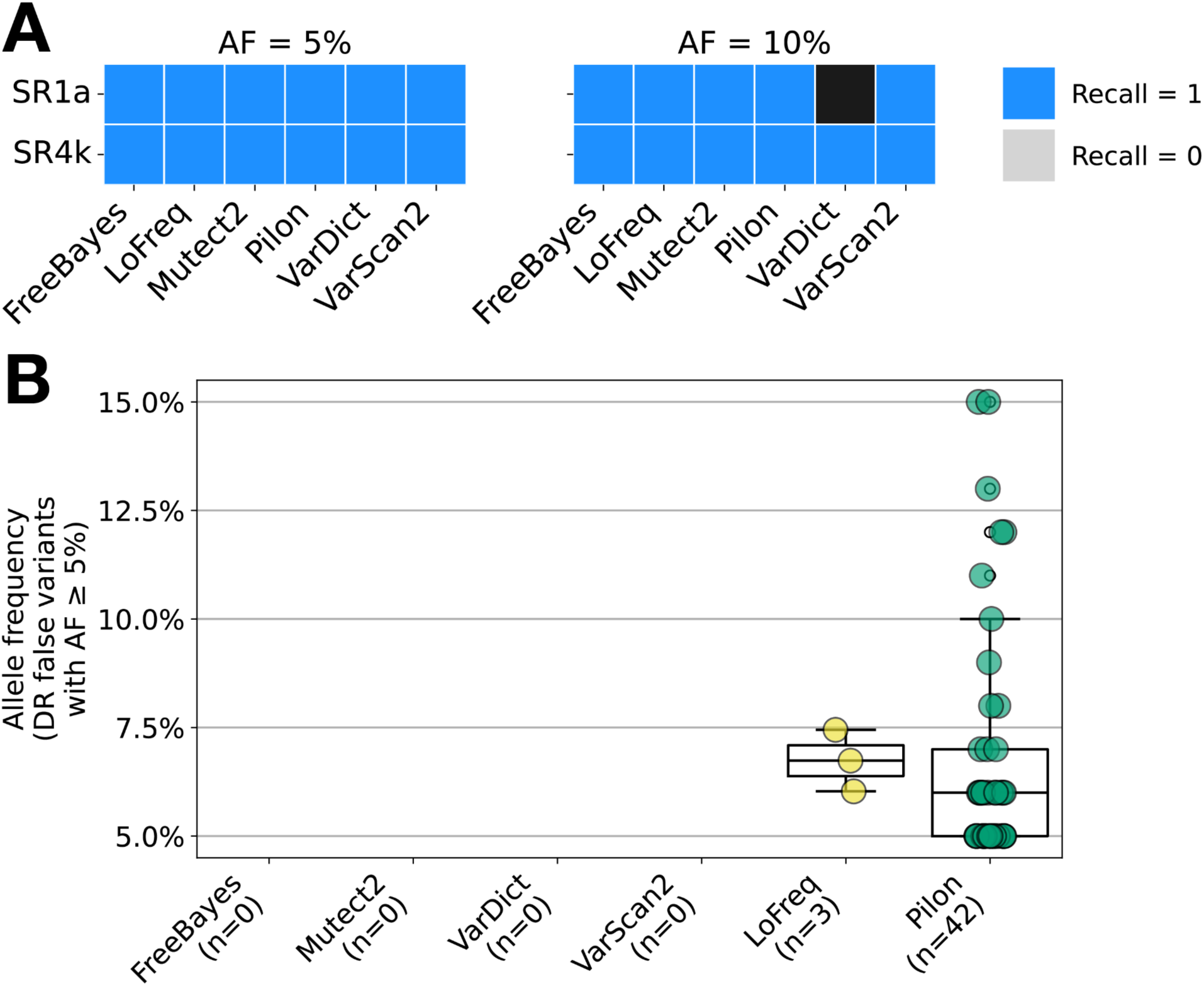
Variant caller performance in in-vitro samples with introduced *rpoB* mutations for AF ≥ 5%. **a** Heatmap of recall for each tool on each in-vitro isolate with rpoB mutations introduced at AF ≥ 5%. VarDict timed out after 3 days for the 10% SR1a isolate. **b** Distribution of the false variant AFs detected by each tool across all DR regions at AF ≥ 5%. The number of total false variants detected at AF ≥ 5% by a tool is shown in parentheses below each tool name.

Of the 6 tools, only Pilon detected both *rpoB* variants at an AF of 1% (Figure S34a), but its precision was low and it falsely identified an average of 87 variants per isolate (AFs 1-4%), 95% of its variants are at AF = 1% (Figure S34b). FreeBayes and VarScan2 were able to identify the AF = 1% *rpoB* variant in the SR1a sample, and both tools also falsely identified an average of 1 variant per isolate.

## Discussion

In this *Mtb*-focused benchmarking analysis, we evaluate a range of factors that influence low-frequency variant calling. Firstly, we consider the genomic context of a variant by comparing the detection of variants in and out of repetitive regions and homopolymeric tracts. Secondly, we study the signal-to-noise ratio by examining the accuracy of variant detection as a function of sequencing depth and within-sample allele frequency. Lastly, we explore reference bias by comparing variant calling in samples simulated from differing background genomes across different *Mtb* lineages and thus, with different levels of baseline genetic divergence from the alignment reference. Across six variant callers, we find the average precision and recall in DR regions to be consistently over 0.92 and 0.82 respectively at a minimum AF of 3% across sequencing depths 50-700x, regardless of background genome. Yet, genome-wide in non-H37Rv strains, a median of more than 300 FP are detected at AF ≥ 1%, with more than a half of these occurring in low mappability regions. This poor precision at low variant AFs, which is reflected in our *in-vitro* analysis, necessitates both a higher minimum variant AF cutoff for genome-wide studies of low-frequency variants, and masking regions of the genome in which variants are inherently harder to detect accurately with short-read sequencing data. Overall, FreeBayes performs consistently well across genomic regions and background genomes, making it a good choice as a broadly applicable tool for both studies of drug resistance and genome-wide within-host diversity.

The high performance of FreeBayes compared to the other tools is possibly due to its haplotype-based variant detection method, as opposed to an alignment-based method. FreeBayes’s local phasing of genotypes into haplotype clusters can prevent variant calls based on poor alignments in low complexity regions, or regions enriched for structural variation, as has been noted in other studies and seen for other haplotype-based tools (Kim *et al*. 2018, Cooke, Wedge, and Lunter 2021, Seah *et al*. 2023). FreeBayes had not previously been benchmarked on low-frequency variant detection in *Mtb* WGS data, though other studies reported that FreeBayes achieved the highest precision for fixed SNVs and a high recall comparable to 6 out of the 7 other benchmarked tools for fixed SNVs and INDELs in *Mtb* WGS data, and the highest sensitivity at the cost of the lowest precision for minor variants in simulated viral data (Said Mohammed *et al*. 2018, Seah *et al*. 2023). Mutect2 is also haplotype-aware, though its precision and recall do not match those achieved by FreeBayes.

VarDict, like Mutect2 and VarScan2, were initially developed for somatic mutation detection. And despite VarDict’s shortcomings with close-proximity variants, all three of these somatic mutation detection tools perform comparably to, if not better than, the other more general-purpose tools we benchmarked. Our study therefore substantiates the use of somatic mutation detection tools for low-frequency variant detection in bacterial data. GATK already encourages this use case with its recent release of a call filter function mode for Mutect2 which fine-tunes it for microbial data. Though LoFreq was developed specifically for low-frequency variant detection in a range of WGS data types, it ranks only above Pilon based on average weighted F1 score. Comparatively, LoFreq is a more conservative variant caller, as observed by others, and exhibits poor performance in repetitive regions of the genome (Said Mohammed *et al*. 2018, Goossens *et al*. 2022). LoFreq was run with its default mapping quality settings min=0 and max=255, consistent with other studies, but it still achieves poor recall in low mappability regions (Said Mohammed *et al*. 2018, Bush *et al*. 2020, Goossens *et al*. 2022). Pilon is a widely used tool for calling variants in microbial data but we note it was not designed for low-frequency variant reporting (Walker *et al*. 2014). For example, (1) INDELs occurring at AF < 25% are not reported, and (2) while low-frequency SNVs can be determined from Pilon’s output using the quality weighted percentage (QP) values, we did observe an abundance of false positive variants with AF = 1%, causing uncertainty around the validity of these low (<10) QP values produced by Pilon.

Overall, our analysis necessitates an approach to low-frequency variant detection that (1) imposes sequencing depth-dependent minimum AF thresholds and (2) mitigates false variants in regions of the genome with poor mappability. For all tools, false variants are prevalent at low AFs in DR and other generally easy-to-call regions. Additionally, our analysis of *in-vitro* data showed that all tested variant callers struggled to distinguish between introduced mutations at AF = 1% and false variants, likely caused by factors such as sequencing error and poor mapping. Consequently, we do not recommend calling low-frequency variants at an AF of 1% with short-read sequencing data at standard depths. Further, the effect of reference bias on precision outside of DR regions is substantial and most pronounced in known low mappability regions, where tool performance is also more variable. False variants in these regions are likely due to non-unique mapping between highly similar, repetitive segments of the genome and are observed at AFs 1-50%. Masking these regions in order to mitigate this issue is standard practice in the field, and our analysis shows it to have noticeable benefits on the precision of FreeBayes. Still, the low mappability regions defined in this analysis are not exhaustive, are not all automatically generalizable across organisms, and mask all positions in three *Mtb* DR-associated genes (*rpoB*, *rpoC* and *rrs*) which will impact the clinical relevance of variant discovery.

Our analysis also shows generalizable characteristics of false variants that allowed us to develop an error model for variant filtering that can be used with or without region masking. The model substantially improves the precision of FreeBayes at a low cost to recall. Alternative approaches for accurately detecting variants in these regions include variant calling against a personal or closely related reference genome and/or ultra deep short-read sequencing (500-1000x) or combined short and long-read sequencing (Rhoads and Au 2015, Schmid *et al*. 2018, Cechova 2020, Marin *et al*. 2022, Vollger *et al*. 2023). Looking forward, graph-based alignment methods for short-reads can offer further improvements for minority variant calling, especially in structurally divergent regions of the genome (Hickey *et al*. 2020, Letcher, Hunt, and Iqbal 2021, Marin *et al*. 2022, Yang *et al*. 2023).

Our analysis is not without limitations. We focus strictly on five *Mtb* genomic backgrounds and examine a subset of variants observed in the *Mtb* population, specifically SNVs and homopolymeric indels. We simulated sequencing data using pre-specified error models that only partially simulate types of error expected in real-world sequencing data. We address this by performing parallel analyses in two simulated sequencing datasets produced by two different simulators (InSilicoSeq and ART), and subsequently benchmarking on experimentally mixed *Mtb* populations. We note that contamination is an important source of low-frequency variant calls especially in conserved regions of the genome, and we do not examine the effect of contamination on error rates of low-frequency variant detection in this study.

Further, we examine precision but not recall outside of DR regions in strains simulated from a non-H37Rv background genome. With these caveats our explicit assessment of mappability and different sources of false positives supports generalizability of our benchmarking exercise to other genomic backgrounds. None of the final six tools discussed in this analysis were developed for *Mtb* specifically, but all achieve practically high and clinically relevant accuracy across high complexity areas of the genome.

## Conclusions

In conclusion, FreeBayes is the top-performing tool for all regions, but low-frequency variant calling is significantly less accurate in low mappability regions. Low-frequency variant detection in DR regions is not affected by reference bias where all tools achieve an average F1 ≥ 0.92 for variant AF ≥ 5% across all background genomes and simulated depths. Our study finds that low mappability region exclusion and minimum variant allele frequency thresholds significantly mitigate high false variant calls genome-wide. Lastly, we provide a new error model that can be used to filter FreeBayes variant calling output and reduce low-frequency false positives while maintaining high recall.

## Methods

### Definition

Throughout this study, we use the term low-frequency (minority or unfixed) variants to refer to alleles present within a single *Mtb* sample at allele frequencies below the consensus threshold of 95%, unless the allele frequency is otherwise specified.

### WGS data simulation and variant calling

We simulated 500 different mutant H37Rv strains in duplicate using ISS and ART. Each strain had a distinct combination of 10 different haplotypes of 50 mutations spanning drug resistance (DR), low mappability (LM), or homopolymeric tract (HT) regions (Supplementary Methods A), simulated depths (5 depths ranging 50 to 700x), and mutant allele frequencies (AFs; 10 AFs ranging 1% to 50%). Strains were simulated in five random replicates by both tools for a total of 5000 WGS simulations. In a second set of simulations, 50 mutant strains were simulated from four non-H37Rv reference genomes belonging to L1, L2, L3 and non-H37Rv L4 respectively (Supplementary Methods B). Sequencing depth and variant allele frequency were varied as above for 20 DR SNVs per strain. Both ISS and ART were used to simulate five replicates for a total of 2000 WGS simulations. Data simulation is further described in Supplementary Methods C.

Each pair of simulated sequencing reads underwent read trimming, alignment to H37Rv and duplicate read removal. Pileup output was also generated for each alignment. The resulting alignment files were used as input into each of the seven benchmarked variant callers. Further details on sequence processing and variant caller parameters are described in Supplementary Methods D, E and F.

### Performance analysis

Overall tool performance in the H37Rv samples was assessed using a weighted F1 score metric, averaged across all strains. For each strain this weighted F1 score was computed as the harmonic mean of the weighted precision and weighted recall, which were determined as the weighted average of the precision and recall, respectively, across each of the regions of interest (DR, HT, LM). Weights were determined based on the expected number of low-frequency variants in each region, determined from a set of clinical isolates (Supplementary Methods I).

For all variant caller performance analyses, reported P-values for pair-wise comparisons were obtained using the Mann-Whitney U test (two-sided) and adjusted for multiple testing using the Benjamini-Hochberg procedure to control the false discovery rate. A significance level of 0.05 was used for all tests.

### Error model for SNV false positive filtering and INDEL allele fraction adjustment

We built a logistic error model to estimate the probability of a false call for an unfixed SNV called by FreeBayes and tested its effect on the FPR and recall of FreeBayes. The following read mapping and quality metrics were used as predictors in this model: base quality, mapping quality, coverage ratio, discordantly-aligned reads ratio, soft-clipped bases ratio and strand bias (full description of these metrics provided in Supplementary Methods L). We trained the model on a ground truth set of unfixed SNVs (n=946, 394 true and 552 false) identified by FreeBayes in short-read mapping of 172 *Mtb* isolates to a personal genome built and polished using hybrid long-reads and short-read sequencing (Supplementary Methods M). The model achieves an AUC of 0.929, precision of 0.812 and recall of 0.967 for SNVs on the training set.

The allele fractions of FreeBayes INDEL calls were adjusted to account for reads that do not sufficiently cover each INDEL, as well as reads with soft-clipping. This process and additional hard filters applied to INDEL calls are described in Supplementary Methods N.

## Acknowledgements

We thank Hu Jin for prompting our inclusion of Mutect2 and advising us on its application for bacterial sequencing data.

## Authors’ contributions

SM and MF designed the benchmarking analysis, interpreted all data and wrote the manuscript. SM generated the simulated data, ran all variant callers on the simulated data and conducted all benchmarking-related analyses. SK designed and trained the error model for filtering FreeBayes output, developed the method we implemented for allele fraction adjustment of INDELs identified by FreeBayes, and processed the raw sequence data for the in-vitro analysis. MM assembled the reference genomes used for the non-H37Rv simulations and determined the set of baseline lineage variants. All authors read and approved the final version of the manuscript.

## Data availability and materials

All code and data required to re-create the simulated data is available at this GitHub repository: https://github.com/shandu-m/benchmark-minority-variants-Mtb. Tables 1-3 in Supplementary File 3 contain run accession numbers for non-simulated isolates used in this analysis. Table 1 details the run accession numbers associated with the sequencing reads used to generate the L1-4 reference genomes from previously published sources (Chiner-Oms *et al*. 2019, Marin *et al*. 2022). The L1-4 reference fasta files are available for download here: https://zenodo.org/records/13761165. Table 2 details the ENA run accession numbers for the Illumina data used in the experimental strain (*in-vitro*) analysis that were made publicly available with the BinoSNP publication (Dreyer *et al*. 2020). Table 3 details the run accession numbers for the clinical isolate sequencing data used to determine expected numbers of low-frequency variants in each of the studied regions.

