## Supplementary File 1 for "Benchmarking within-sample minority variant detection with short-read sequencing in *M. tuberculosis*"

### Supplementary Results and Methods

---

**Description of Supplementary Files**

**File S1 (.pdf)** Supplementary results and methods.

**File S2 (.pdf)** Supplementary Figures S1-34 and Supplementary Tables S1-10.

**File S3 (.xls)** Additional data.

Table 1 Run accessions for the sequencing reads used to generate the L1-4 reference genomes.

Table 2 Run accessions for the experimental strains.

Table 3 Run accessions for the clinical isolate cohort. Samples that have been made publicly available have an ID in the Biosample Accession column. The last 8 isolates (in grey) are slated to be made public as part of a different publication from our lab referenced in the table. These were not yet publicly available as of May 21, 2026.

#### Supplementary Results

##### A. Simulated allele frequencies show high fidelity to the expected allele frequencies

The standard error of the simulated AFs for ISS and ART was respectively 0.011% and 0.015% for AF < 10%, and respectively 0.051% and 0.084% for AF ≥ 10% (averaged across all depths and background genomes). Most of the variability in the simulated AFs for each simulator tool can be attributed to the higher standard error of the simulated AFs in LM regions, and the simulated LM variant AF is consistently lower than the expected AF across all depths (Figure S1, Table S1). For both ISS and ART, the simulated values are consistent across all replicates (all P-values > 0.001 for replicate comparisons of simulated coverage and AF in each expected sequencing coverage group, and simulated base quality).

##### B. BinoSNP runs inefficiently and detects many of the same false positives as the other variant callers

BinoSNP requires a user to supply it a BED file with regions in which to look for mutations and a table with an entry for each position in these regions specifying the reference and alternate allele. We tested BinoSNP on a subset of 500 strains (one of the replicates), supplying input regions of length $10^1$, $10^2$, $10^3$, $10^4$, $10^5$ and $10^6$. With a region size of $10^5$, the run times were already well above 10 hours per strain, while the strain runs with a region size of $10^6$ timed out after 5 days (Figure S2). The BinoSNP tool offers no parallelization, though it would be possible to split the input region files and call variants for a strain in multiple regions concurrently. We do find, however, that BinoSNP is not able to filter out FP variants detected by other tools successfully. For each of the strains simulated by InSilicoSeq (n=2500), we provided BinoSNP with all positions at which a variant had been detected by at least one of the other variant callers and determined the percentage of FPs detected by at least one of the other variant callers at AF > 1% that was also detected by BinoSNP. Across all strains, BinoSNP detects an average of 85% of the FPs detected by at least one other variant caller (minimum = 33%), and this average is similar in each subset of strains simulated with a different sequencing depth (averages across all sequencing depth groups range 84-88%). BinoSNP displays no bias towards genomic region, detecting an average of 85-90% of the FPs per strain detected by at least one other variant caller at AF > 1% in all regions of the genome outside of HT regions. In HT regions, BinoSNP detected only 7% of the FPs detected by at least one other variant caller, though in preliminary analysis, we found the INDEL reporting of BinoSNP to be unreliable.

**C. Variant caller failure to detect simulated variants in close proximity to fixed baseline lineage variants**

FreeBayes consistently missed one mutation in L1, L2 and L4 samples, and two mutations in L3 samples, and VarDict consistently missed two mutations in L3 samples (Figure S3). These shortcomings in the recall of FreeBayes and Vardict were linked to three consistently missed mutations across all simulated L1-4 strains, all three of which were outside of the LM regions. FreeBayes missed the *gyrA* mutation at position 7582 in at least 94% of strains in each lineage group, as well as the *pncA* mutation at position 2289050, which was missed in 98% of L3 strains. Both of these mutations were found to occur within 3bp of a lineage variant (Figure S4). This proximity to a lineage variant was able to differentiate both of these mutations that were consistently missed by FreeBayes from the other mutations that FreeBayes did not consistently miss (Figure S5a). While VarDict also missed the *pncA* mutation at position 2289050 in 98% of L3 strains, it additionally missed a *pncA* mutation at position 2289072 in 76% of L3 strains. This mutation at position 2289072 was not as close to any L3 lineage variants, or other simulated mutations, as the other two consistently missed mutations. This mutation could not be differentiated from other unproblematic mutations for any base pair window size (Figures S5b-d).

**D. FreeBayes and Pilon detect an excessive number of false positives at specific simulated sequencing depths**

Both FreeBayes and Pilon exhibited an unexpected excess number of FPs at a subset of the tested sequencing depths: most significantly at 100-200x for FreeBayes and 200-400x for Pilon (Figure S6). This pattern is present in the ART data as well (Figure S7). Using a set of 20 clinical isolates, with average sequencing depths distributed across our simulated depths, we investigated the total number of variants found by each tool in each depth group (Figure S8). We discovered a similar pattern as exists in the simulated data: FreeBayes found an excess of variants at depths of 100-200x, while Pilon found an excess of variants at depths of 200-400x.

**E. Alternative WGS data simulator comparison: variant caller accuracy rankings and major FP trends are consistent with the ISS-simulated data**

FreeBayes still achieves the highest accuracy in the ART H37Rv simulations according to weighted F1 score (Fig. 1a, Figure S9a). While Pilon still achieves the lowest accuracy too, the two pairs of tools achieving intermediate weighted F1 scores are swapped (Mutect2-VarDict and LoFreq-VarScan2). All variant callers perform consistently between the ISS and ART simulations in DR and HT regions (average F1 score difference ART-ISS < ±0.03 in both regions), performance in LM regions is lower overall (average F1 score difference ART-ISS = -0.34) and variant caller ranking is significantly shuffled (Fig. 1b, Figure S9b). Consistent with the ISS simulations, variant caller F1 scores are most variable for variant AF < 10% where FreeBayes achieves the highest F1, and LoFreq, VarDict and VarScan2 most successfully recapitulate simulated variant AF (Figures S10, S11).

While fewer FPs are detected by all tools in the ART data than in the ISS data (1.45E-05 and 2.21E-05 median FPR respectively across all strains and tools), a higher FPR is still observed in LM regions than in DR regions or elsewhere in the genome (Figure S12). Additionally, FPs in DR regions remain at low AFs <10.2% in the ART data (Fig. 3, Figure S13). Finally, though low mappability regions are still prone to high AF FPs in the ART data, there are more high AF FPs outside of LM regions in the ART data than in the ISS data.

#### <u>Supplementary Methods</u>

##### A. Variant position choices for simulations

<u>i. SNVs in drug resistance regions</u>

SNV positions for each *Mtb* mutant are randomly chosen from the list of category 1 or 2 confidence level SNVs reported by the WHO (variants with final confidence gradings for resistance associations of “Assoc w R - Interim” or “Assoc w R”; WHO 2021).

For our analysis of false positives, we considered a broader definition of drug resistance regions. These were based on gene set categorizations from *Vargas et al. 2021*.

<u>iii. Insertions in homopolymer tracts</u>

Homopolymer tract (HT) regions came from the list of regions with a single nucleotide repeated ≥7 times generated by *Vargas et al.* in Supplementary Data File 3 (Vargas *et al.* 2023). From these HT regions, we included only those with previously known antibiotic resistance/tolerance associations (Safi *et al.* 2019), or reported to have such an association by *Vargas et al. 2023*. We also included the 6-guanine homopolymer in *Rv0678* (at position 193) in which spontaneous G-insertions and G-deletions at position 193 were observed in a study on *Mtb* clinical isolates and associated with low-level bedaquiline resistance (Xu *et al.* 2023). Frameshifts in this HT have been observed in several other studies on *Mtb* clinical isolates (Villellas *et al.* 2017, Ismail *et al.* 2018, Kadura *et al.* 2020, Peretokina *et al.* 2020).

For analysis of false positives, mutations in all HT regions were considered (not only those with antibiotic resistance/tolerance associations).

<u>iii. SNVs in low mappability regions</u>

Low mappability regions were defined based on pileup mappability scores, a metric of how easy it is to align reads uniquely across the genome. To determine these scores, the pupmapper pipeline (https://github.com/maxgmarin/pupmapper/) was run on the H37Rv genome. This pipeline determines a pileup mappability score for each genomic position based on k-mer uniqueness up to a defined edit distance. We used pupmapper with a k-mer size of 50bp and a maximum mismatch threshold (edit distance) of 4bp. In addition, we included genomic positions with a low empirical base-level recall (EBR; Marin *et al.* 2022).

A low mappability region for our purposes is defined as a feature (gene or intergenic region) in which all positions have a pileup mappability score < 0.8 and EBR < 0.8. We focus on PE/PPE genes, which are often excluded in analyses due to their repetitive regions, in addition to intergenic regions. Positions in these regions were then randomly selected for each simulated mutant strain.

**B. Non-H37Rv (L1-4) reference genomes and establishing baseline lineage variants**

Reference genomes used for the non-H37Rv simulations were generated by *de novo* PacBio HiFi read assembly and iteratively polished with PacBio and Illumina reads to generate a complete circular genome as described previously (Marin *et al.* 2022). One sample from each of the lineages 1-4 was selected as the reference genome for the L1, L2, L3 and L4 simulated strains (Additional File 3). We characterized fixed/consensus mutational differences between H37Rv and these reference genomes by aligning each reference to the H37Rv genome with minimap2 (Li 2018), and using paftools.js as described by *Marin et al. 2022*. These consensus differences were excluded from analyses assessing false variant detection.

**C. WGS data simulation**

We used InSilicoSeq and ART to simulate paired-end Illumina reads spanning the whole *Mtb* genome. In the first set of simulations, sequencing reads for 500 mutant *Mtb* strains were simulated from the H37Rv reference genome (chromosome NC_000962.3). We considered (1) all 409 SNV DR mutations with known or likely resistance associations reported in the 2021 WHO *Catalogue of mutations in Mycobacterium tuberculosis complex and their association with drug resistance* (WHO 2021), (2) SNV mutations at 46800 LM positions, and (3) 1-base insertion mutations at 18 HTs associated with antibiotic resistance or tolerance, for a total of 47227 potential variant positions (see Supplementary Methods A). In each strain we simulated 50 different variants: 20 SNVs at the DR-associated positions, 20 SNVs at the LM positions, and 10 1-base insertion mutations in the HTs with previously known antibiotic resistance or tolerance associations. We generated 10 different haplotypes of these 50 mutations. In the second set of simulations, sequencing reads were simulated from a reference genome belonging to lineage 1 (L1), 2 (L2), 3 (L3), or a non-H37Rv lineage 4 (L4) genome. We simulated 100 strains from each background, for a total of 400 *Mtb* strains, each with 20 simulated DR SNVs. Across both sets of simulations, we varied sequencing depth (50x, 100x, 200x, 400x, 700x), and variant allele frequency (1%, 2%, 3%, 4%, 5%, 10%, 20%, 30%, 40%, 50%). Each combination of parameters was simulated in five replicates.

#### D. WGS processing

Each pair of *Mtb* sequencing reads was processed as follows: (1) read trimming from the 3' end to a quality of 25 with PRINSEQ-lite (v0.20.4; Schmieder and Edwards 2011), (2) BWA-MEM (v0.7.17; Li 2013) alignment to the *Mtb* reference genome H37Rv (NCBI RefSeq NC_000962.3), (3) duplicate read removal with Picard (v2.8.0; Broad Institute 2019), and (4) pileup output (mpileup) file creation with SAMtools (v1.15.1; Li *et al.* 2009).

#### E. Variant calling

Seven variant callers were run on the simulated sequence data alignments: BinoSNP, FreeBayes, LoFreq, Mutect2, Pilon, VarDict and VarScan2. For overall analysis we attempted to run each tool with similar or equivalent parameters. For example, for all tools for which we were able, we set the minimum number of alleles required for a variant to be called to 2, as is the default for FreeBayes, VarDict and VarScan2. We also set the minimum variant allele frequency to 0.01 (1%), our minimum simulated variant frequency. When we tested setting the minimum variant allele frequency to 0, some tools picked up significantly more false positives, specifically FreeBayes, which called on average 7000 more variants in a subset of strains (n=25; Figure S14). For tools without these user-defined parameters, we wrote a script to filter the output of these tools to include only calls with at least 2 variant-supporting reads at a fraction of 0.01 of the coverage at that position. We note that recovery of variants at a frequency of 1% (allele fraction of 0.01) will be limited by the ability of the simulation to accurately simulate enough reads, especially for lower coverages. Ploidy was also set to 1 where possible. Additional variant caller details, parameter tuning and normalization for haplotype-aware tools is described in the next section.

#### F. Variant caller parameters, normalization and filtering

<u>BinoSNP (v1.0.1)</u>

BinoSNP was run with its default parameters with a P-value of 0.05 used to filter variants.

<u>FreeBayes (v1.3.6)</u>

For main performance analysis, FreeBayes was run with its default parameters except for our settings of ploidy (1) and minimum alternate fraction (0.01), as well as the settings of minimum mapping quality (30) and minimum base quality (30) as is used by *Nimmo et al.* who used FreeBayes to call low-frequency variants in *Mtb* sputum samples (Nimmo *et al.* 2019). FreeBayes outputs a quality score for each putative variant and suggests filtering variants with this metric on its GitHub page (https://github.com/freebayes/freebayes). In preliminary analysis on simulated data to determine an appropriate quality score threshold for variant filtering, we found overlapping quality score distributions for true positive and false positive variants. As a result, variants called by FreeBayes were not filtered on quality scores for the simulated data analysis.

LoFreq (v2.1.5)

LoFreq was run in its default form to support INDEL calling: *lofreq indelqual* was used to insert INDEL qualities into the input alignment file (BAM) and the resulting alignment file was run through *lofreq call-parallel*. The default P-value of 0.01 was used after testing LoFreq on a subset of simulated data to determine the optimal value (results were consistent across the data simulated by both InSilicoSeq and ART).

Mutect2 (GATK v4.5.0.0)

Mutect2 was run following the GATK best practices documentation for somatic short variant discovery (https://gatk.broadinstitute.org/hc/en-us/articles/360035894731-Somatic-short-variant-discovery-SNVs-Indels), excluding the contamination determination step as we ran all tools on simulated data: candidate variants were called using Mutect2, orientation bias artifacts were determined using *LearnReadOrientationModel*, and putative variants were labeled with *FilterMutectCalls* in microbial mode. In the *FilterMutectCalls* step, calls are assigned labels such as “PASS”, “orientation”, “weak_evidence” and “strand_bias.” Typically only “PASS” calls are used for downstream analysis. We found, however, that the “orientation” label seemed to be inflated in the simulated data in a way that was not reflective of true error. Further, for the InSilicoSeq simulated data, we found that the average true positive rate increased by 0.447, while the average false positive rate only increased by less than 1E-09 when “orientation” labeled calls were included. An average true positive rate increase of only 0.026 was observed for the ART simulated data (average false positive rate only increased by less than 1E-10), indicating that Mutect2 was more sensitive to orientation bias in the InSilicoSeq data. The “orientation” labels were not prevalent in our cohort of clinical isolates used in this study (<1% of variants called by all tools were labeled with “orientation” by *FilterMutectCalls*), further substantiating that the “orientation” over-labeling of true positives by *FilterMutectCalls* is a simulation-specific issue.

Pilon (v1.24)

For main performance analysis, Pilon was run with the same parameter settings used by *Marin et al.* in a paper benchmarking the empirical accuracy of short-read sequencing in *Mtb*: minimum mapping quality and minimum depth were set to 40 and 5 respectively (Marin *et al.* 2022). The variant allele read count was calculated with QP and DP Pilon VCF output fields, *(QP/100)*DP*. The Pilon output VCF was filtered to include all positions at which there were at least 2 high quality reads supporting the variant allele and at which the quality-weighted support for that base was at least 1 (corresponding to an allele frequency of 1%).

VarDict (Java port v1.8.3)

VarDict was run in its default form which includes feeding its output through the *testsomatic.R* and *var2vcf_paired.pl* scripts for paired variant calling, as described by VarDict's GitHub page (https://github.com/AstraZeneca-NGS/VarDict). The VarDict output VCF was filtered to include all positions at which there were at least 2 reads supporting the variant allele and at which the allele frequency for that base was at least 1%.

VarScan2 (v2.3)

VarScan2 tools *mpileup2snp* and *mpileup2indel* were run with minimum variant frequency set to 0.01 (1%) and their outputs were combined into one TSV file. A P-value of 0.01 was used after testing VarScan2 on a subset of simulated data to determine the optimal value (InSilcoSeq and ART results were consistent) and is supported by literature (Mariner-Llicer *et al.* 2024). Otherwise, all VarScan2 parameters were set to their default.

Variant set normalization and filtering

To ensure accurate comparison of variants called between tools and to the introduced variants, we took steps to standardize the variant output across all tools. We used the *norm --multiallelics -any* function (with a reference genome fasta file) from bcftools (v1.21) to split multi-alleleic sites and left-align INDELs in the FreeBayes, VarDict and Mutect2 output VCF files (Danecek *et al.* 2021). The *vcfwave* tool from vcflib (v1.0.14) was additionally used to simplify complex alleles called by FreeBayes and VarDict (Garrison *et al.* 2022). Multi-nucleotide polymorphisms called by FreeBayes, VarDict and Mutect2 were split into SNPs in a custom Python script. Finally, for all tools we used a custom Python script to extract variant calls from the tool output VCF or TSV file and perform the necessary filtering.

**G. Real variant analysis**

To obtain a baseline expectation for the number of variants in each of the studied regions (DR, HT, LM), we called variants in 38 clinical isolates from the MIC ML Consortium. To estimate the relative number of low-frequency variants in each of the three regions of interest (DR, HT, LM), for each isolate we considered only 1-base insertions in HT regions, and SNVs in DR, LM and all other regions, that were called by *all* six tools (excluding BinoSNP, Table S2). Here, low-frequency variants were defined as variants with average AF (across all tools) between 5% and 95%.

We created two groups from these 38 isolates to investigate differences in the number of variants detected for (1) isolates with difference average sequencing coverages and (2) isolates from different lineages (L1-4). In the first analysis, 20 lineage 4 isolates were studied. Isolates were picked so that there were four isolates with a mean coverage approximately equal to one of the five studied depths (50x, 100x, 200x, 400x, 700x). For the second analysis, six isolates from each of lineages 1-4 were studied. Mean coverage was balanced across all lineage groups and restricted to 50-200x. We also had drug resistance data for 17 drugs; on average, 28% of isolates were resistant for a given drug. The average number of low-frequency variants detected per region was similar between all isolates in each sequencing coverage group and lineage (all P-values > 0.05 for the Mann-Whitney U tests to compare the number of variants in each region between each sequencing depth group and each lineage; Benjamini–Hochberg correction for multiple testing with FDR = 0.05).

**H. *In-vitro* isolate analysis**

We compared the accuracy of each tool for a set of 6 *in-vitro* isolates each with a single frequently occurring *rpoB* mutation at a frequency of 1%, 5% or 10%. These isolates were prepared by the authors of the binoSNP tool from the *Mtb* reference lab strain (*Mtb* H37Rv ATCC 27294; Dreyer *et al.* 2020). A Ser531Leu mutation was introduced in the SR1a isolates, while a His526Pro mutation was introduced in the SR4k isolates. Alignments were prepared and variant calling was performed as described in Supplementary Methods D, E and F.

**I. Performance analysis**

For each simulated strain we extracted all variants reported by each variant caller and an introduced variant was considered to be present if it occurred in the variant caller output file, regardless of the reported AF. The numbers of ground truth/true positive (TP), false negative (FN) and false positive (FP) variants were determined for each strain. False positive variants were considered at AF ≤ 50%. All tools detected additional variants than the determined baseline lineage variants at AF > 50% in the non-H37Rv strains (median number of these per strain is 77-164 across all tools). These variants are not described as many have very high allele frequency (90-95%) and because a proportion of these are expected due to lineage differences. The latter may have been missed in the baseline lineage variant determination or their call AF determination may have been underestimated by neighboring variants especially for haplotype-based variant callers.

Weights for the weighted F1 score were defined as the median number of low-frequency variants across 38 clinical isolates for each region as described in Supplementary Methods G. We assumed that variant calling accuracy outside of defined DR, HT and LM regions was comparable to accuracy in *Mtb* DR regions in this weighting schema. Precision and recall for the H37Rv samples was determined in all regions (DR, HT, LM). Precision and recall in the L1-4 samples was determined in DR regions only (considering the 20 introduced DR SNVs) and any FP detected in DR regions. Known fixed variants in the L1-4 references relative to H37Rv were excluded.

Accuracy as a function of minimum variant AF (AF 1-50%) was assessed considering all TP, FN and FP with an AF equal to or greater than that minimum variant AF. The per base false positive rate (FPR) per strain was calculated for each region (DR, HT, LM, elsewhere) relative to the region size.

**J. Assessing the accuracy of variant caller allele frequency estimation**

The simulated allele frequency for each introduced variant in a strain was determined from the raw read counts in the pileup out (mpileup) file generated from that strain's alignment file (see Supplementary Methods D). The allele frequency estimated by each variant caller for the introduced variants was compared to this simulated allele frequency.

**K. Comprehensive low mappability regions**

We expanded our definition of low mappability regions for analysis beyond the low mappability regions used as candidate variant simulation positions. These regions are based on a range of factors including pileup mappability scores, or determined through empirical studies of the *Mtb* genome in work within our lab and with collaborators.

These comprehensive low mappability regions include:

1. Regions of low pileup mappability determined by pupmapper (https://github.com/maxgmarin/pupmapper/) in H37Rv and four other representative genomes from lineages 1-4.
2. minimap2-based H37Rv homologous regions, which allow for INDELs between homologous sequences (Li 2018).
3. A set of refined low confidence (RLC) regions, describing regions that account for major sources of error in Illumina WGS analysis, determined by *Marin et al.* (Marin *et al.* 2022).
4. Genes with high homology to other bacterial genomes (*aspT*, *clpB*, *hsp*, *rpoB*, *rpoC*, *rpsC*, *rrl*, *rrs*, *tuf*).
5. The first 500bp of the genome which are prone to clusters of erroneous low-frequency variants due to poor alignment and generally lower coverage (in *dnaA*).

Cumulatively, these regions account for 10.6% of the *Mtb* genome.

**L. Sources of false positive variants**

To better understand sources of false positive variant calls we studied read mapping and quality characteristics at variant site positions in the L1-4 strains. These read mapping and quality characteristics were determined at each variant site as follows:

1. *Base quality*: average base quality across all variant calls at the site, determined from the alignment pileup.
2. *Mapping quality* (FreeBayes, Mutect2, Pilon and VarDict only): average mapping quality of the variant allele, determined from the mapping quality field in the VCF of each variant caller.
3. *Coverage ratio*: ratio of the site coverage to the average regional coverage. This was computed as the ratio of the total site coverage to the maximum of the rolling average coverage from the left or right (window size of 100bp).
4. *Discordantly-aligned reads ratio*: ratio of the number of discordantly-aligned reads at the site to the total site coverage. Discordantly-aligned reads were defined as all reads mapped to a variant position in an orientation other than “LR” (Left-Right), determined from the alignment file.
5. *Soft-clipped bases ratio*: ratio of the number of soft-clipped bases at the site to the total site coverage. Soft-clipped bases were determined from the CIGAR strings in the alignment file.

6. *Strand bias* (FreeBayes only): magnitude of the deviation from 0.5 of the proportion of forward reads to total reads. Forward read count was determined from the “SAF” FreeBayes VCF field (number of alternate observations on the forward strand).

**M. Linear error model for SNV FP filtering and validation on simulated data**

To build a ground truth set of unfixed SNVs and INDELs, we used hybrid assemblies constructed from short- and long-reads generated for 172 *Mtb* samples. PacBio HiFi reads were assembled using 3 iterations of flye (v2.9.2; Kolmogorov *et al.* 2019), circularized with Circlator (v1.5.5; Hunt *et al.* 2015), and then polished with Illumina reads using Pilon (v1.23; Walker *et al.* 2014) generating “personal reference genomes.” The Illumina reads were preprocessed using fastp (v1.0.1) for adapter trimming and removal of reads shorter than 50 base pairs (Chen *et al.* 2018), and only reads mapped to the MTBC (taxid 77643) and its lineage by kraken2 (v2.1.3) using the standard database (downloaded in June 2020) were retained (Wood, Lu, and Langmead 2019).

To call low-frequency variants from the personal reference genomes, we aligned Illumina reads from the previous step to the personal genome using BWA-MEM (v0.7.19) with a seed length of 80 (Li 2013). We then marked duplicates with Picard (v3.4.0) and performed variant calling with FreeBayes (v1.3.10) using minimum mapping quality = 30, minimum base quality = 30, minimum alternate allele count = 2, and minimum allele fraction = 0.01 (Garrison and Marth 2012, Broad Institute 2019). Unfixed variants were transferred from the personal genome coordinates to H37Rv coordinates using *paftools liftover*, which is part of the minimap2 suite (v2.30; Li 2018). The resulting variants made up the set of real unfixed variants.

The same Illumina reads were aligned to the H37Rv reference genome (NCBI RefSeq NC_000962.3), followed by variant calling using the same tools, versions, and parameters as above for the personal reference genomes. SNVs detected from H37Rv with a within-sample allele fraction ≥0.05 and ≤0.95 were passed into a mixed effects logistic model implemented in R (v4.4.2) using the lme4 package. The five fixed effects in the model were the number of discordantly paired reads normalized to coverage, number of clipped bases normalized to coverage, average base quality of bases supporting the variant, ratio of coverage at the site to the rolling average, and the absolute value of the difference between 0.5 and the proportion of reads supporting the variant that are in the forward orientation. The rolling average of coverage was computed with a window size of 100 base pairs, and we took the maximum of the rolling averages computed from the left and right directions.

The error model was trained on 946 low-frequency SNV calls (394 real, 552 not real) from the 172 WGS samples with matched personal reference genomes. The 552 false variants were detected when aligning Illumina reads to H37Rv, but not when the same reads were aligned to the personal reference genomes. The single random effect was the sample of origin because all variants from all samples were pooled into a single model, and therefore, the predictions for the 3357 candidate variants in samples not in the training set could only be made based on the estimated fixed effects. After fitting the model, the classification threshold for dichotomizing predicted probabilities was selected to be 0.46 to maximize the sum of precision and recall on the training set.

For each simulated strain, we filtered the set of FreeBayes variants to include only SNVs, extracted the metrics used as fixed effects in the model for each variant, fit the model and retained only variants with a predicted probability > 0.46. We explored the reduction of false variants and loss of true variants after (1) error model filtering, (2) error model and hard filtering (forward and reverse strand allele counts ≥ 2, depth ≥ 5, mapping quality ≥ 40), and (3) error model filtering, hard filtering and low mappability region and rRNA gene masking.

**N. Adjusting allele fractions to correct unfixed INDELs**

Calling low-frequency INDELs is generally less affected by reference bias because variant callers require greater evidence to call INDELs than SNVs. The primary issue in accurately calling low-frequency INDELs is reads not sufficiently covering an INDEL. These reads artificially bring down the allele fraction of the INDEL, making it appear as though a consensus indel is low-frequency and inflating the number of low-frequency INDELs when simply thresholding on allele fraction.

After variant calling using FreeBayes using the same parameters as for SNVs, INDELs were left-aligned and normalized using bcftools (v1.21; Danecek *et al.* 2021). For each INDEL with an allele fraction 0.05-0.95 (inclusive), we then extracted all the reads from the pileup at the position where the INDEL begins. We excluded all reads with soft clipping and all reads that start or end within 10 base pairs of the putative INDEL, including reads that terminate in the middle of the INDEL. We then determined if the start and end positions for reads that support the INDEL are significantly different from those of reads that do not support any INDEL. We compared the medians of the distributions using two Mann-Whitney U tests, one for the start positions and one for the end positions. If both tests returned P-values < 0.01, then this was considered evidence that the reads that do not support the INDEL do not sufficiently cover it. These reads would be unable to support an INDEL, even if it truly exists, and so they too were excluded. We then recomputed the allele fraction using all remaining reads that do and do not support the INDEL (if both P-values from the Mann-Whitney U tests were < 0.01, then these two values are equivalent, and the new allele fraction is 1).

For each simulated strain, we filtered the set of FreeBayes variants to include only INDELs, adjusted the allele fractions for these INDEL variants as described above, and retained only variants with an adjusted allele fraction above 0.05. We explored the reduction of false variants and loss of true variants after (1) allele fraction adjustment, (2) allele fraction adjustment and hard filtering (forward and reverse strand allele counts ≥ 2 for any INDEL except deletions >10bp, depth ≥ 5, mapping quality ≥ 40, adjusted INDEL-supporting reads ≥ 5, variant site coverage ≥ one-third of the genome-wide median), and (3) allele fraction adjustment, hard filtering and masking of low mappability regions, rRNA genes and regions within 100bp of insertion sequences or phages.
