## Supplementary File 2 for "Benchmarking within-sample minority variant detection with short-read sequencing in *M. tuberculosis*"

### Supplementary Figures and Tables

---

#### Supplementary Figures

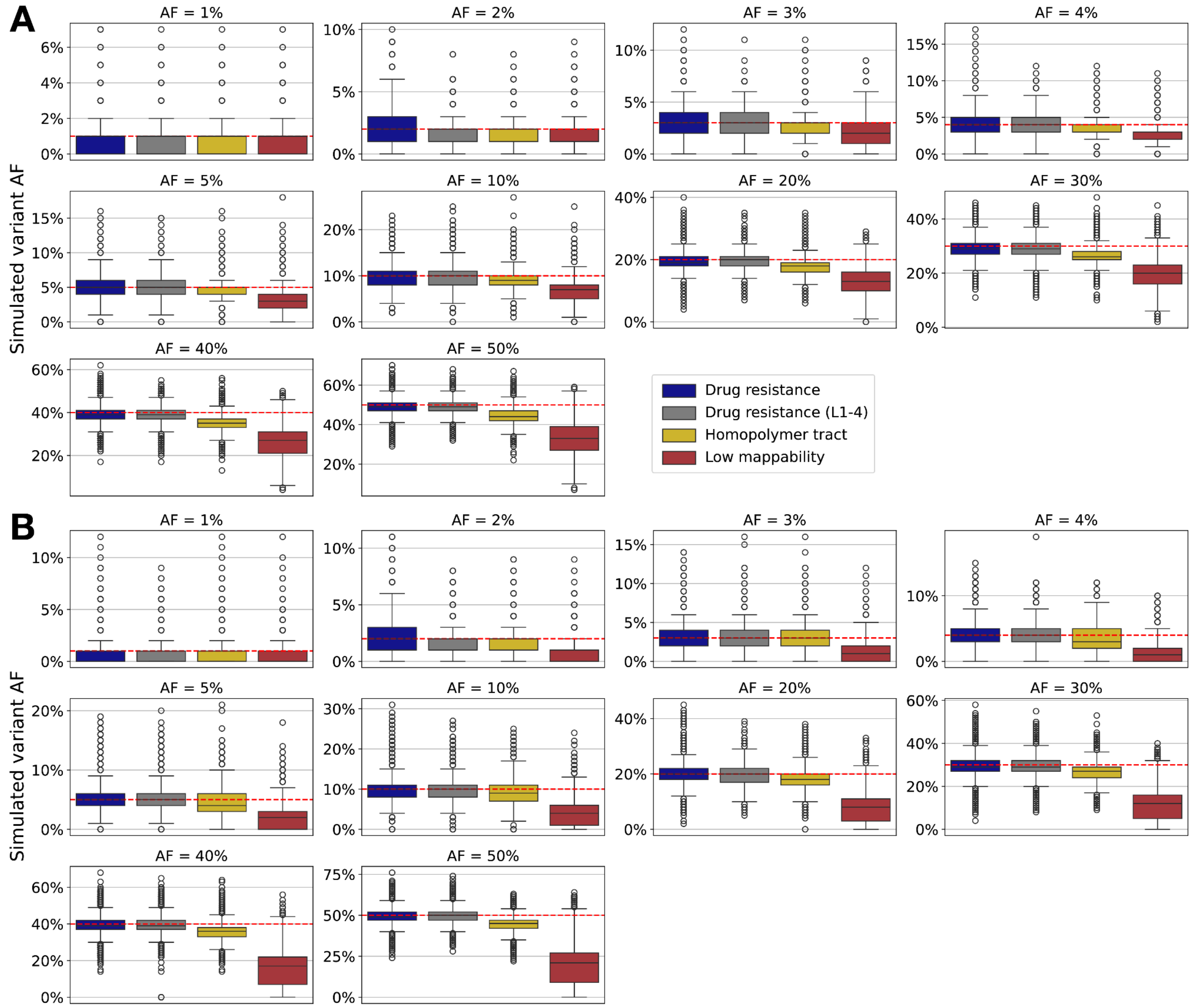


**Figure S1** Simulated variant allele frequency achieved in each genomic region considered. **a** Simulated AFs achieved by InSilicoSeq (ISS). **b** Simulated AFs achieved by ART. The red dotted line indicates the expected allele frequency. The simulated variant AFs tend to be lower than expected in each region, and this is most pronounced in LM regions.

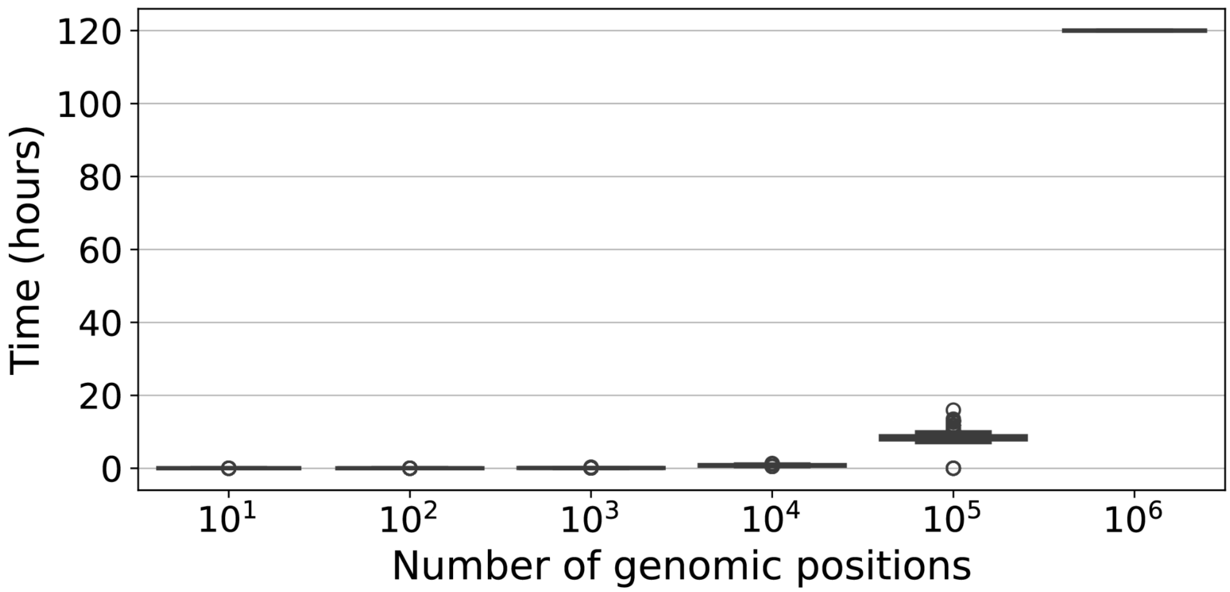


**Figure S2** BinoSNP CPU runtime for a range of input positions from $10^1$-$10^6$. The distribution of the runtimes for each input position set is shown for 500 strains. The BinoSNP runs with $10^6$ positions timed out after 5 days.

**Figures S3-5** Simulated mutations in the L1-4 strains consistently missed by FreeBayes and VarDict.

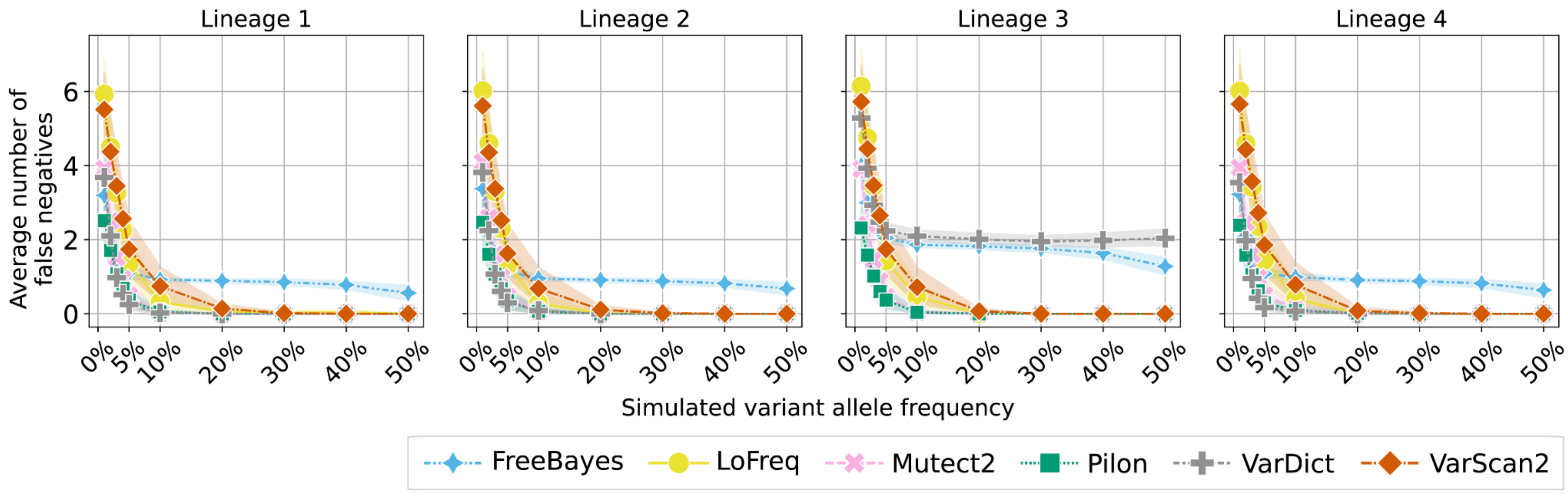


**Figure S3** Average number of false negatives for each tool across variant AFs for mutations introduced in DR regions only (L1-4 strains).

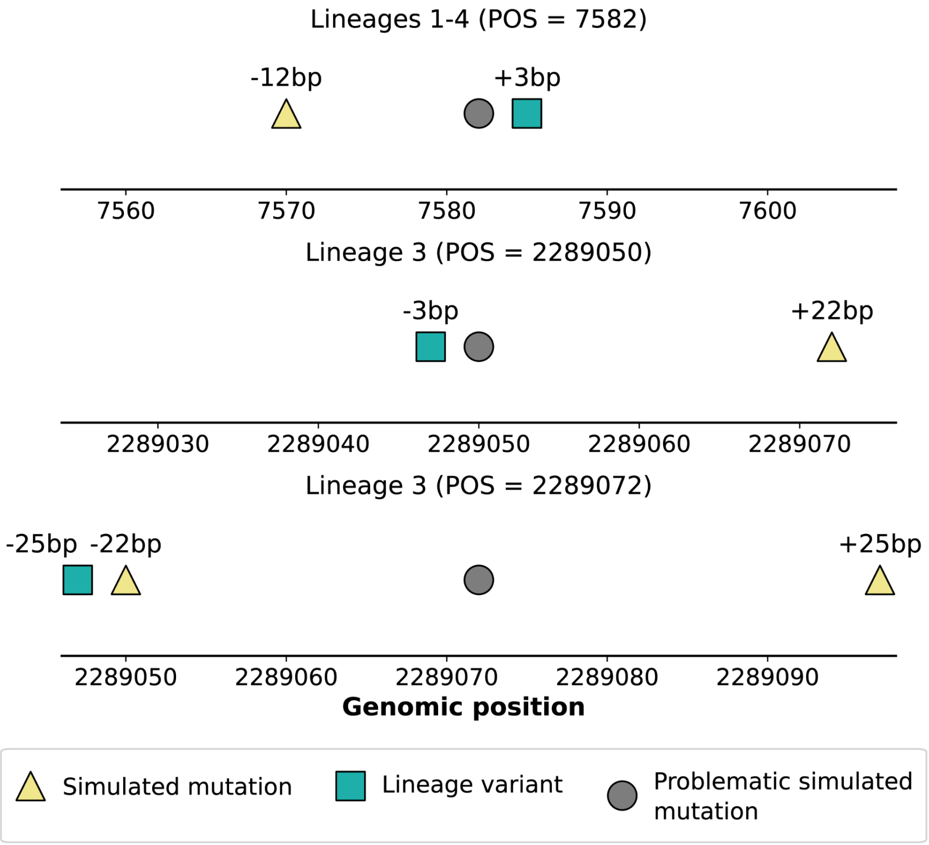


**Figure S4** Distance between problematic simulated variants at DR positions and nearby (within 25bp) (1) lineage variants, and (2) additional simulated variants.

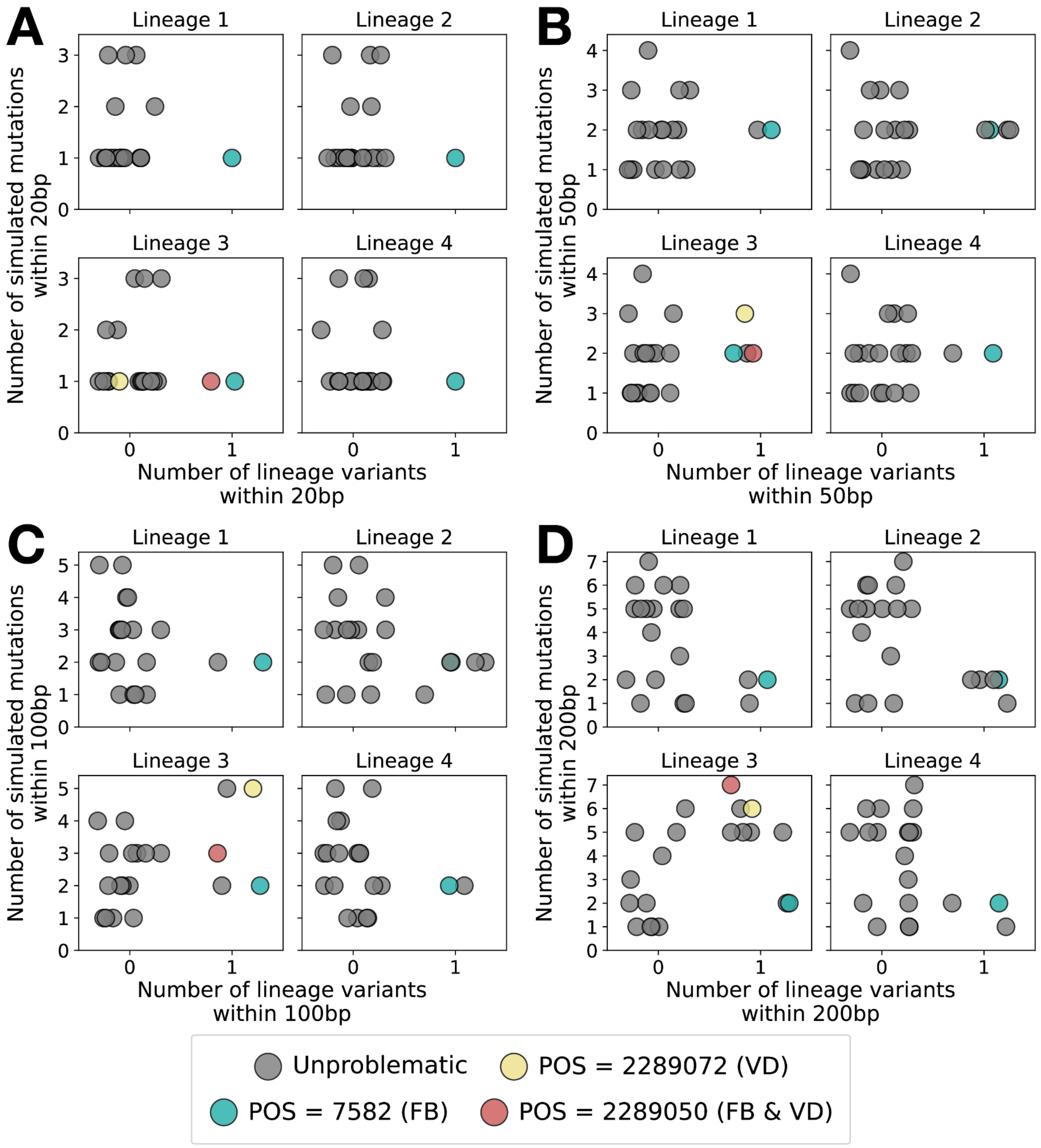


**Figure S5** Number of nearby simulated mutations plotted against number of nearby lineage variants for each simulated DR mutation in the L1-4 strains for increasing window sizes. **a** Window size ±20bp. **b** Window size ±50bp. **c** Window size ±100bp. **d** Window size ±200bp. The three problematic mutations are colored as per the legend: POS = 7582 was consistently missed only by FreeBayes in L1-4, POS = 2289050 was consistently missed by FreeBayes and VarDict in L3, and POS = 2289072 was consistently missed only by VarDict in L3.

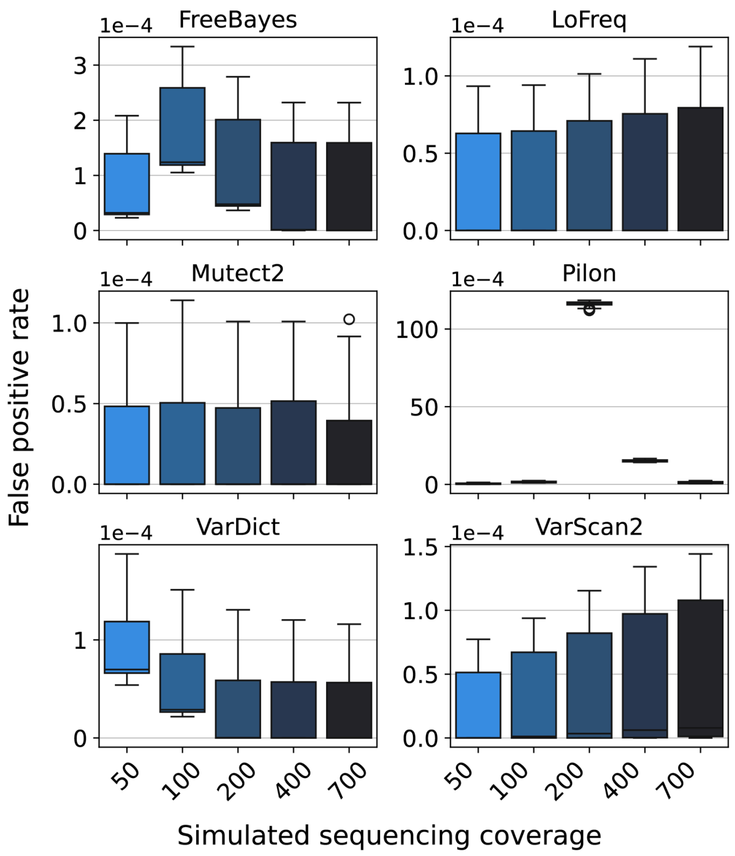


**Figure S6** Genome-wide false positive rate across all strains in each simulated sequencing coverage group. False positive rate is calculated per-base across the entire genome. All tools were run on the same simulated strains, and further, the excessively high number of FP detected by Pilon in strains simulated at 200x is consistent with the excess number of variants detected by Pilon in clinical isolates with depths ~ 200x (see Supplementary Results D), motivating a lack of simulation bias in this result.

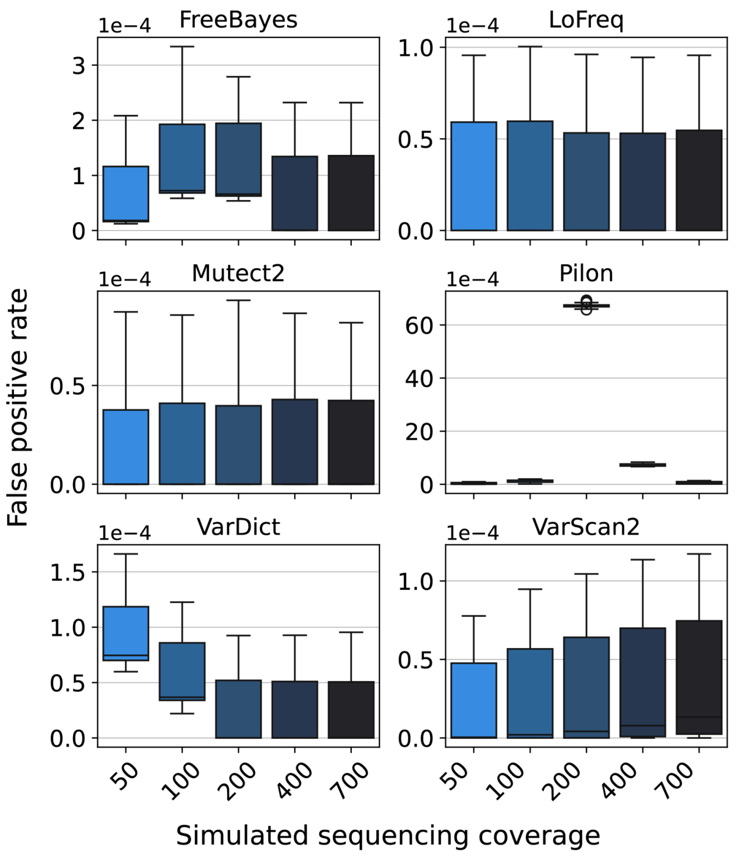


**Figure S7** Genome-wide false positive rate across all strains in each ART-simulated sequencing coverage group. False positive rate is calculated per-base across the entire genome. The FPRs for FreeBayes in strains simulated at 100x-200x and for Pilon in strains simulated at 200x-400x is noticeably higher than the FPRs for those tools at other depths, which is consistent with the ISS-simulated data.

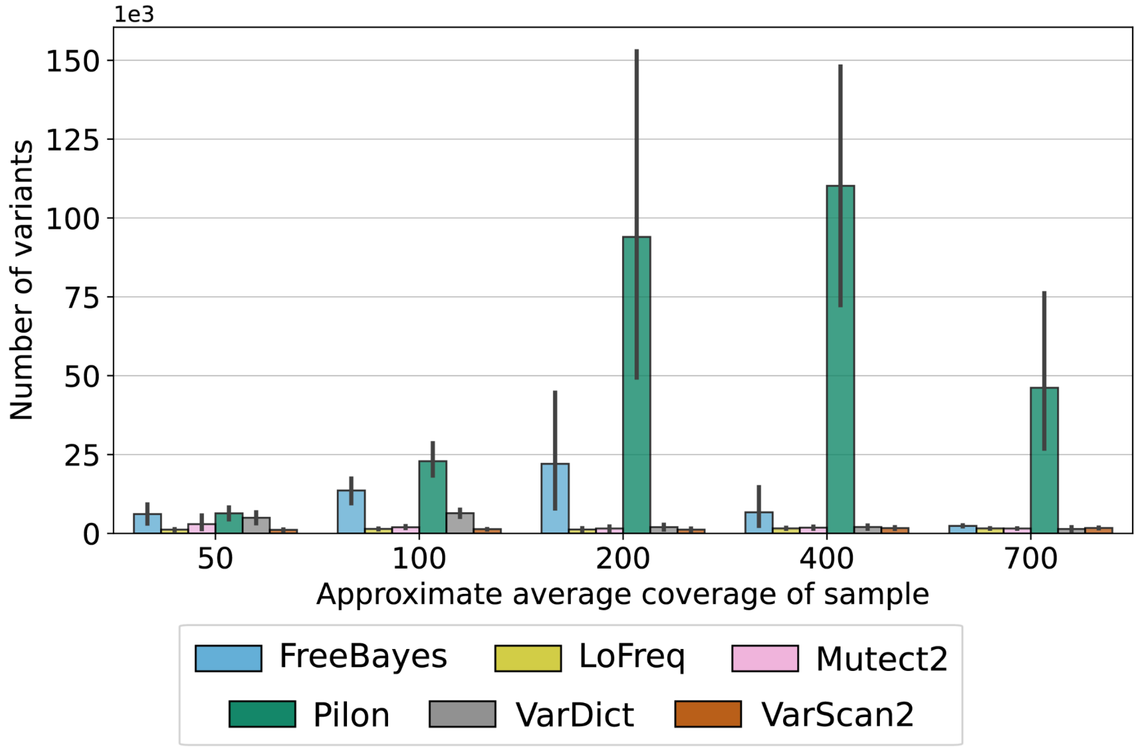


**Figure S8** Distribution of total number of variants found by each tool in clinical isolates (n=20) with different average sequencing coverages. Each bar displays the average number of variants found per isolate by each tool, and the error bar indicates the 95% confidence interval.

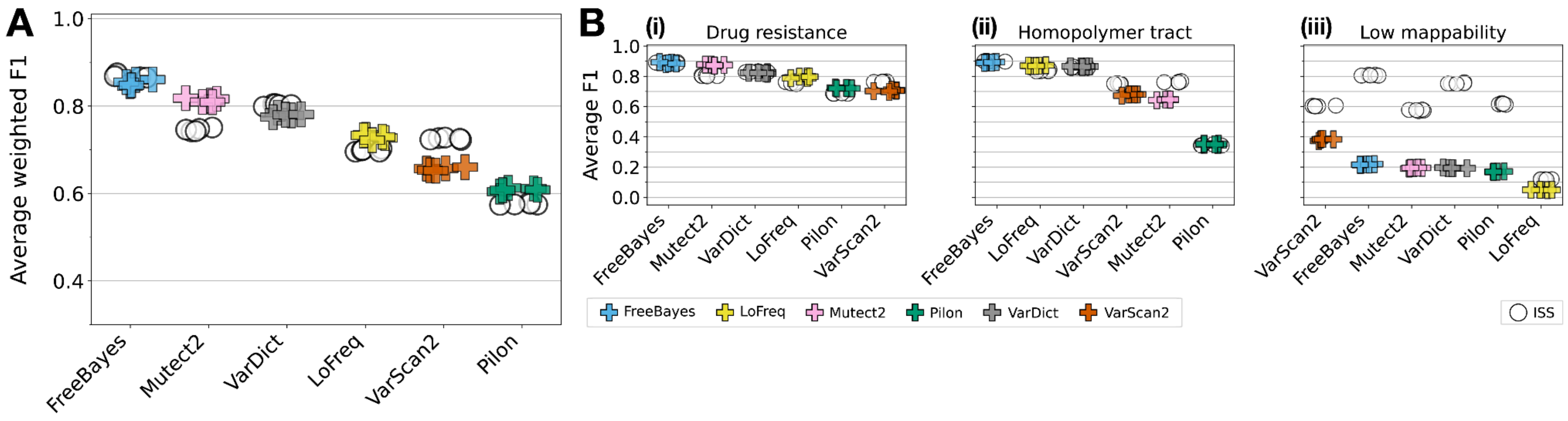


**Figure S9** Overall variant caller accuracy in the ART-simulated data. **a** Weighted F1 score achieved by each variant caller in the H37Rv strains, averaged over simulated variant AFs, depths and mutation regions. Each point represents the average weighted F1 score for one of the replicate simulations. ISS data points are white. **b** Average F1 score achieved by each variant caller in the H37Rv strains (i) drug resistance, (ii) homopolymer tract, and (iii) low mappability regions. ISS data points are in white.

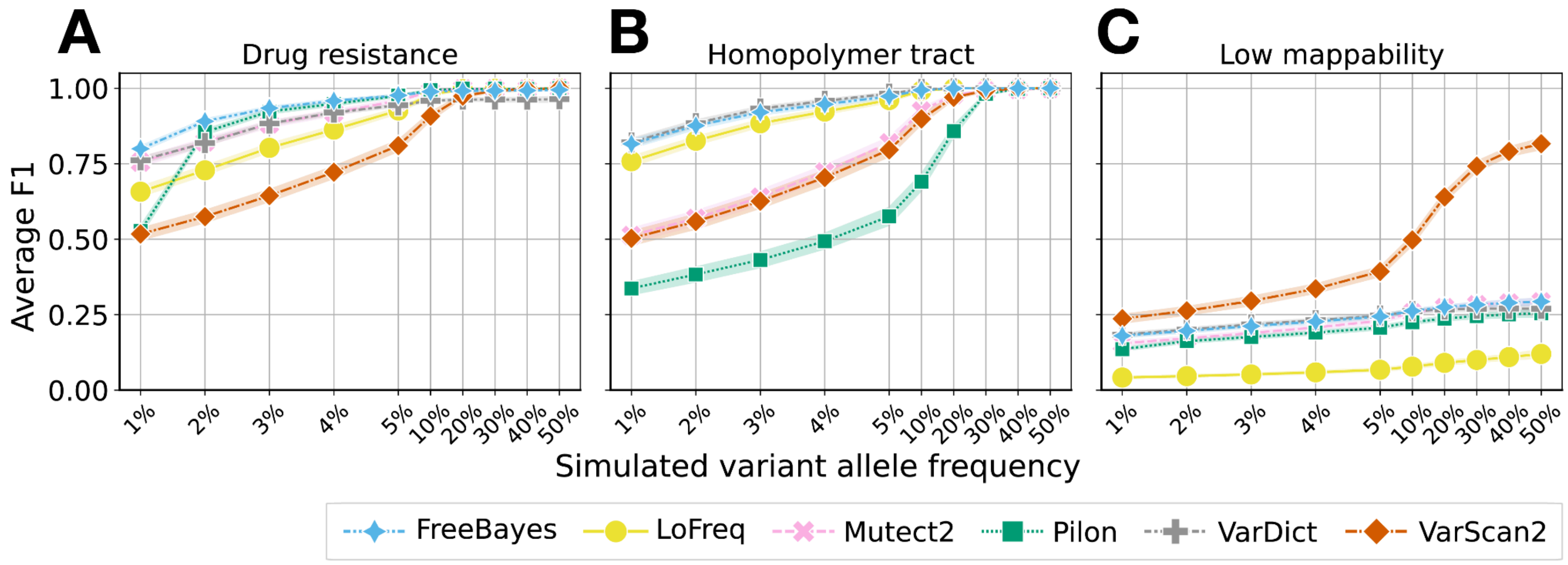


**Figure S10** Average cumulative F1 across variant AF pooled over depths of 50x, 100x and 200x in the ART-simulated strains. **a** Average cumulative F1 in drug resistance regions (H37Rv and L1-4 strains). **b** Average cumulative F1 in homopolymer tract regions (H37Rv strains only). **c** Average cumulative F1 in low mappability regions (H37Rv strains only). To compute the average cumulative F1, we computed cumulative precision and recall as a function of increasing minimum variant AF for each of the six tools, averaged over haplotype, depths 50-200x and replicate. The band around each line represents the 95% confidence interval. Note that the x-axis tick gaps are not proportional to the actual simulated variant AF, and are larger for AF < 10% as this is where the greatest tool-wise differences occur.

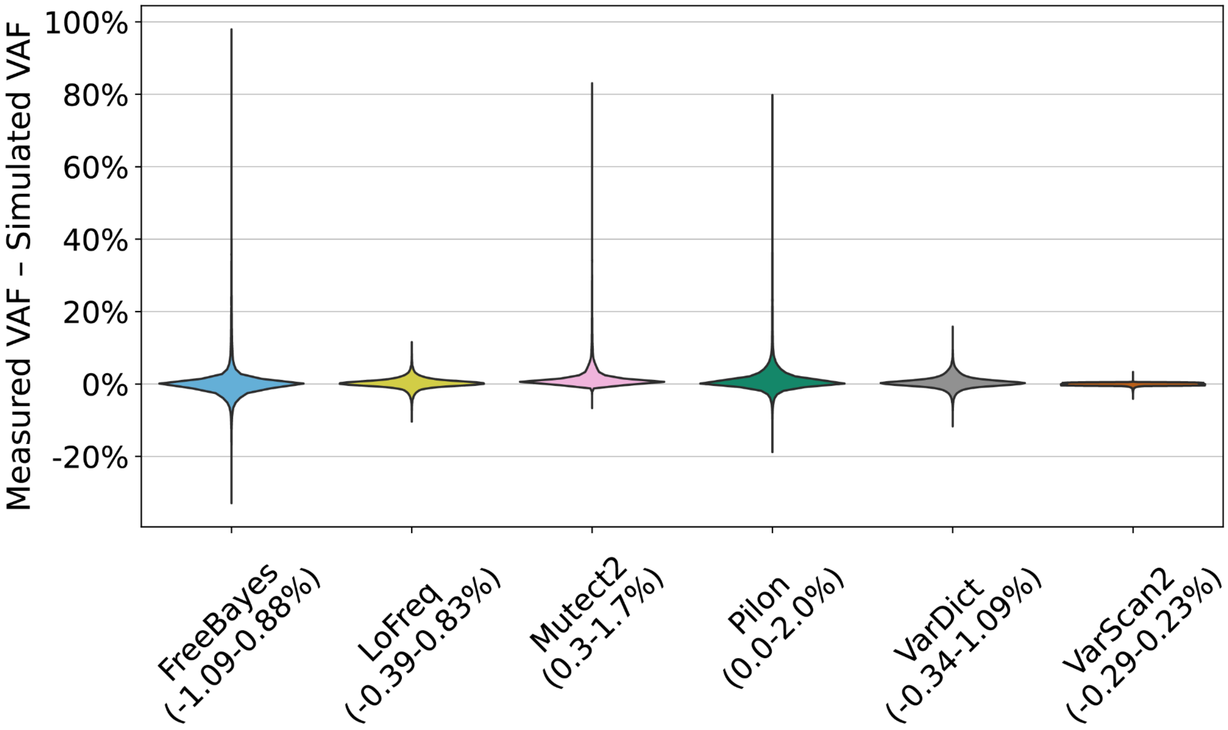


**Figure S11** Distribution of the difference between the measured allele frequency and the simulated allele frequency for variants across all variants, depths, haplotypes and replicates (ART-simulated data).

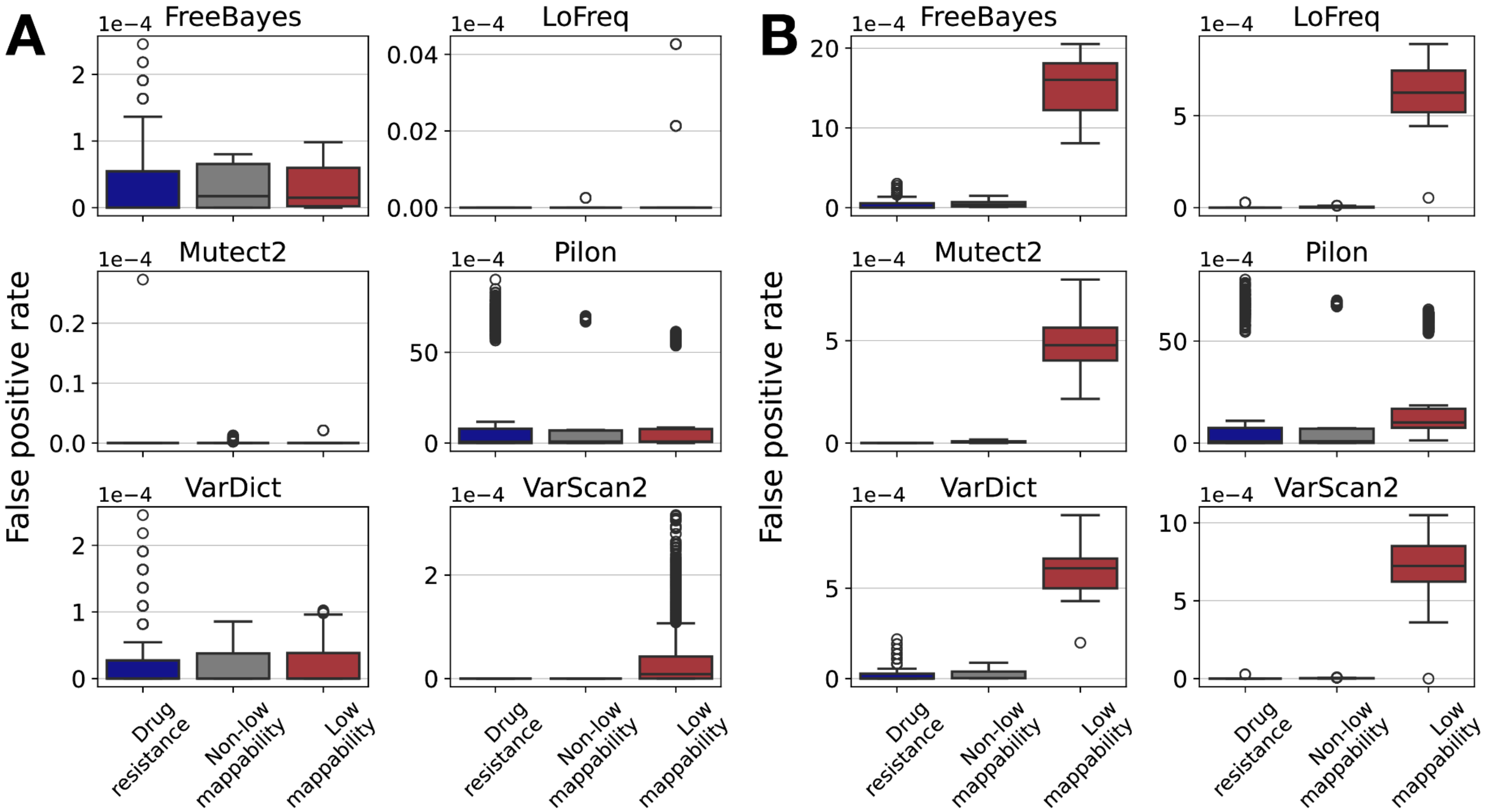


**Figure S12** False positive rates by tool in drug resistance, non-low mappability and low mappability regions in the ART-simulated strains. **a** The distribution of false positive rates per strain and region in H37Rv strains. **b** The distribution of false positive rates per strain and region in L1-4 strains. Each region is defined to be mutually exclusive for this comparison i.e. the non-low mappability regions do not include the drug resistance regions. Each box plot shows the distribution of FPRs in each region and for each tool. Note that the subplots do not share the same y-axis. The pairwise comparisons between each region for each tool are statistically significant in both groups of strains after Benjamini-Hochberg correction (Mann-Whitney U test with FDR = 0.05).

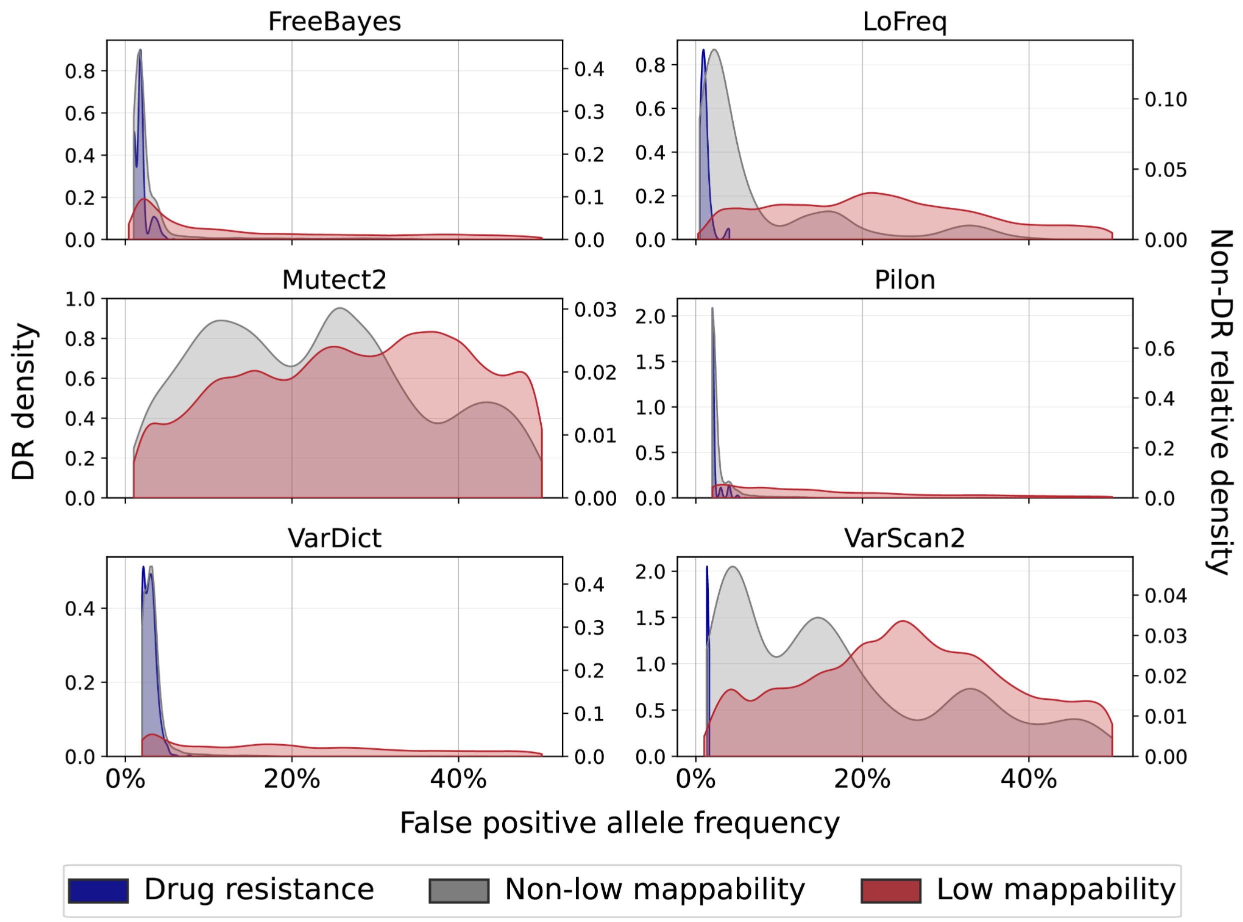


**Figure S13** False positive allele frequency distribution in the ART-simulated L1-4 strains by regions. Each region is defined to be mutually exclusive for this comparison i.e. the non-low mappability regions do not include the drug resistance regions. The left y-axis displays the densities of the DR AF distributions, and the right y-axis displays the relative densities of the low mappability (non-DR) and non-low mappability (non-DR) AF distributions (normalized independently). For Pilon we include only the FP with AF > 1% (a median of 29% of Pilon FPs across all strains have AF = 1%). All DR FP occur at AF < 11% and FP in the other two regions occur at AFs 1-50%. Between 1-82% of the FP in non-low mappability regions occur at AF > 10%, while more than 60% of the LM FPs occur at AF > 10% for all tools except FreeBayes and Pilon (48% and 59% of the LM FPs have AF > 10% for these tools respectively).

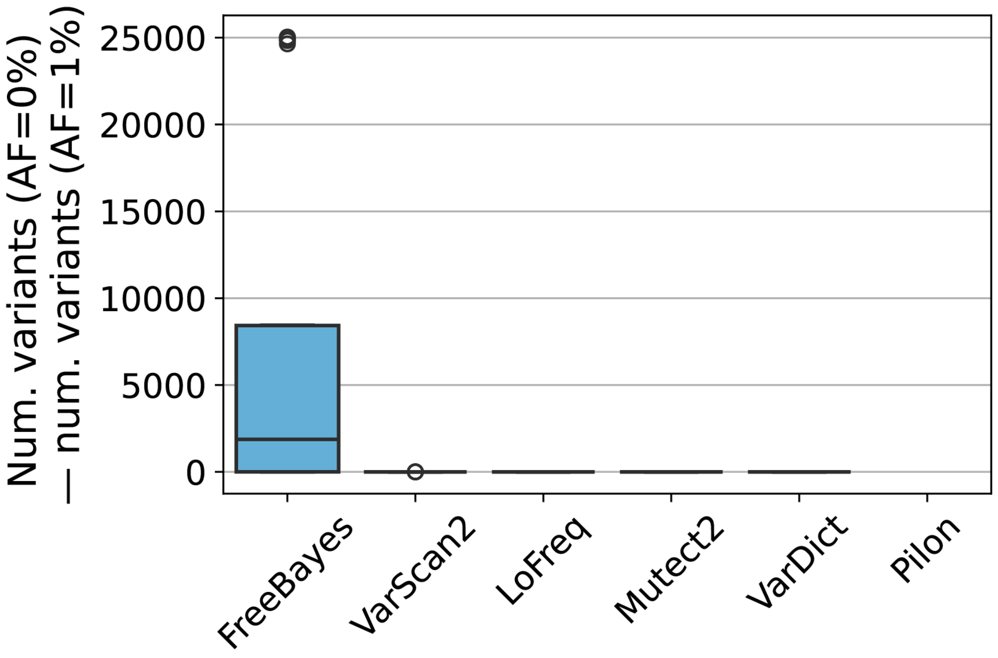


**Figure S14** Distribution of the difference in the number of variants called by each tool (except Pilon) when the minimum variant allele frequency is set to 0% versus 1%. FreeBayes called, on average, 7018 more variants. VarScan2 called at most 5 more variants per strain. No difference was observed in the number of variants called by LoFreq, Mutect2 and VarDict. Pilon was excluded because 1% is the minimum reported quality-weighted allele frequency for a call.

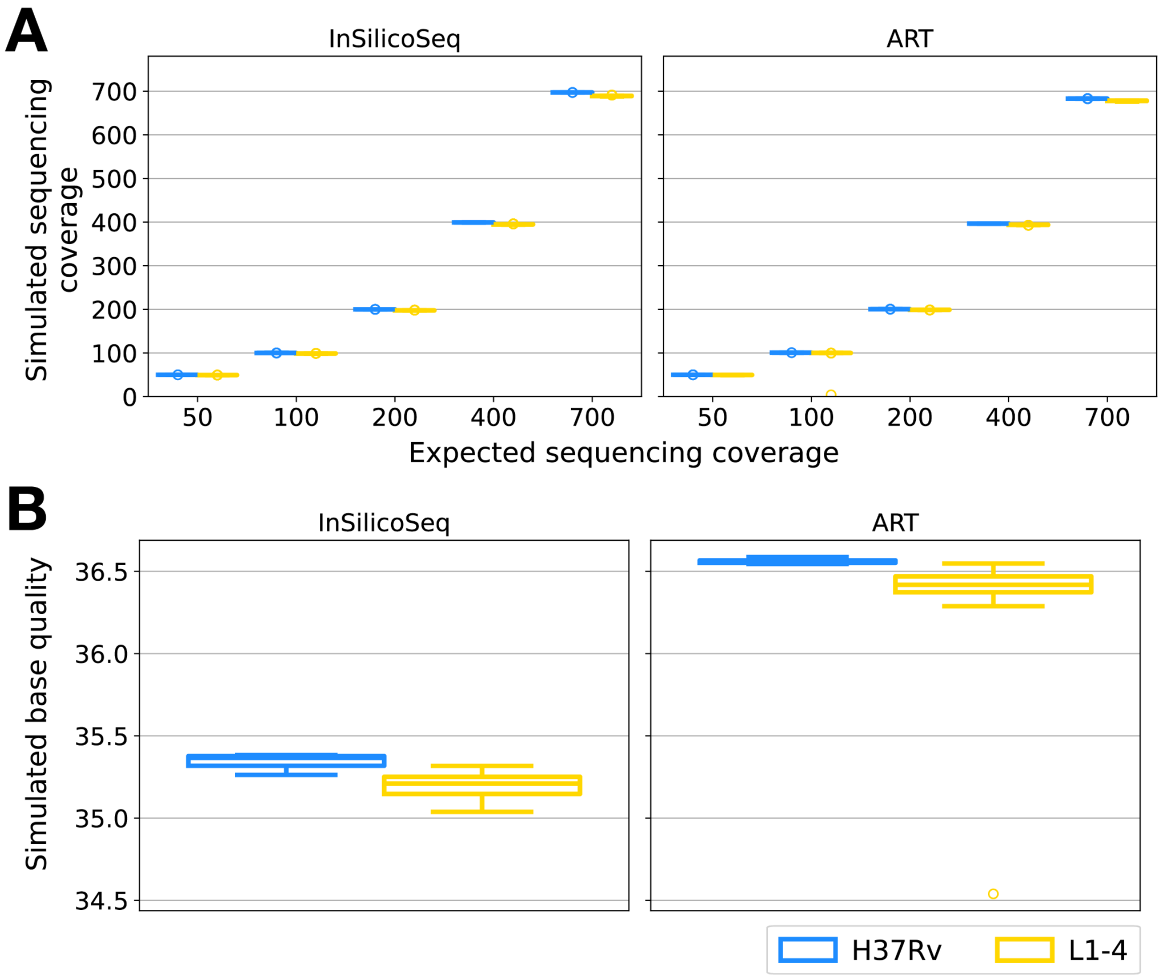


**Figure S15** Simulated sequencing coverage and base quality achieved by InSilicoSeq (ISS) and ART. The coverage and base quality are displayed for the strains simulated from H37Rv and those simulated from an L1-4 background genome separately. **a** Simulated sequencing coverage. The differences in the average sequencing coverage between the H37Rv and L1-4 strains are similar for the ISS and ART simulations and minimal for both (average H37Rv – L1-4 coverage for ISS and ART respectively is 0.56, 0.37 at 50x, and 7.83, 5.18 at 700x where this difference is highest across all expected sequencing depths. **b** Simulated base quality. All average strain base qualities were within 35.03-35.39 for the ISS simulation, and within 34.53-36.59 for the ART simulation. The outlier with an average depth ~ 0 in the ART L1-4 100x strain group (average simulated depth = 4.2x) and outlier with an average base quality under 35.0 in the ART L1-4 simulation group (average base quality = 34.5) are the same strain. Both the simulated depth and simulated base quality were determined by running Pilon (as described in Supplementary Results F) on each simulated strain to produce a VCF entry for each genomic position. The DP (depth) and BQ (base quality fields) were then extracted for each position and averaged for each strain. The distribution of these averages is displayed in the plots above.

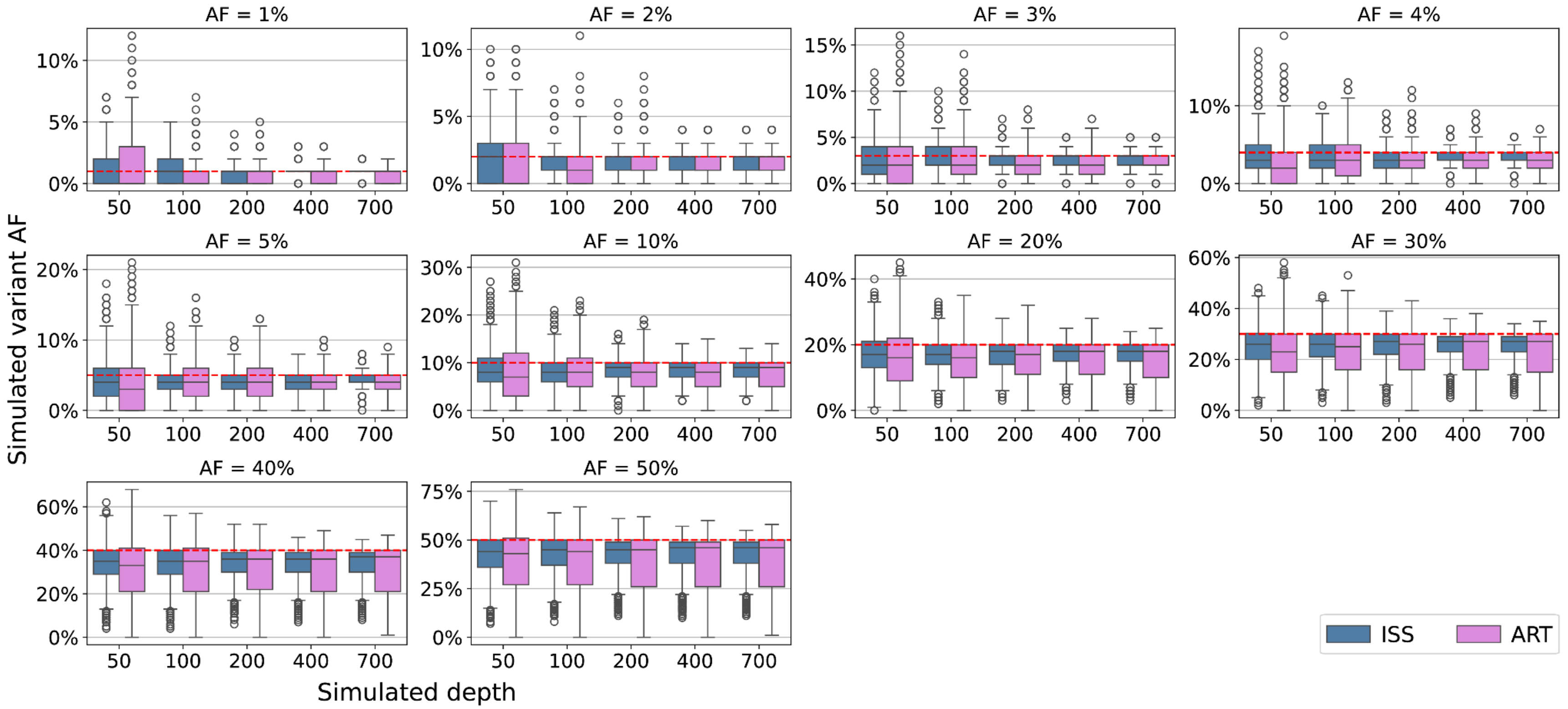


**Figure S16** Simulated variant allele frequency achieved by InSilicoSeq (ISS) and ART in each expected sequencing coverage group. The dotted red line indicates the expected allele frequency. The maximum average difference in the deviation of variant allele frequency between depth groups was low at 0.1% for ISS (200x vs 400x, expected AF = 1%) and 0.2% for ART (100x versus 200x, expected AF = 2%).

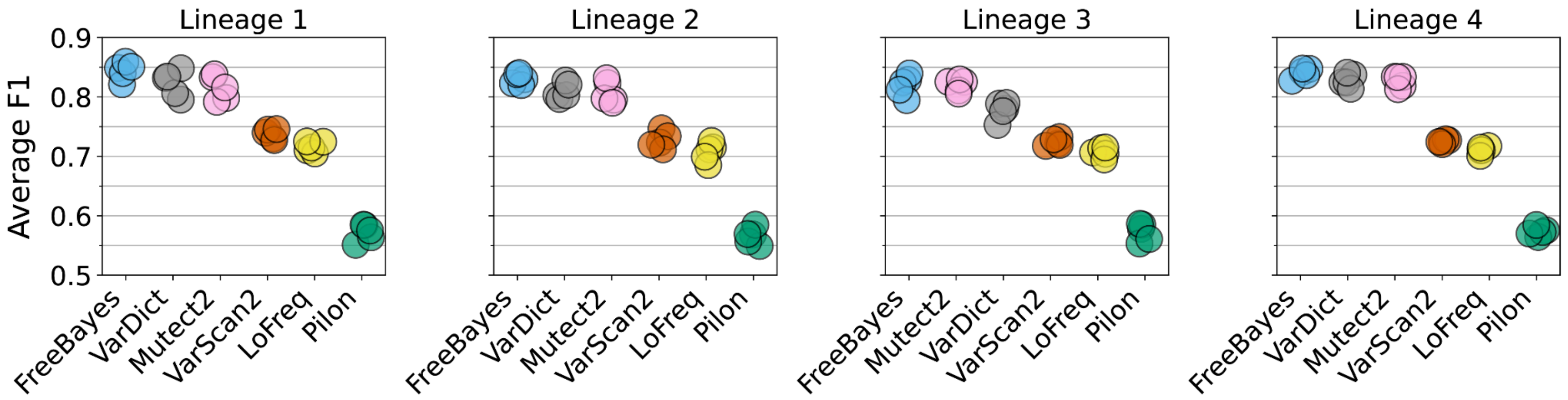


**Figure S17** F1 score achieved by each variant caller, averaged over simulated variant AFs and depths. Each point represents the average weighted F1 score for one of the replicate simulations.

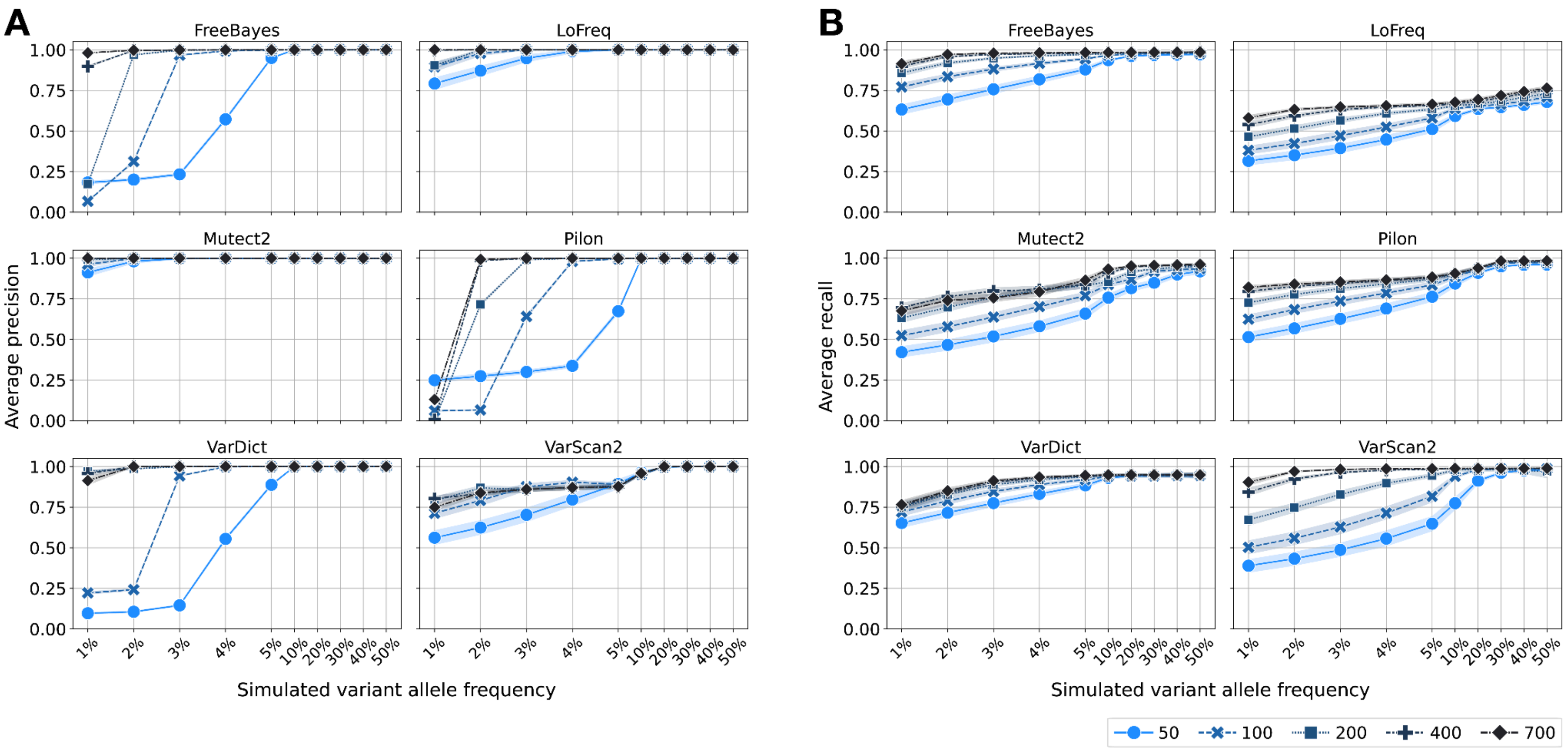


**Figure S18** Average precision and recall across variant allele frequency (1-50%) and sequencing depth (50-700x) in the H37Rv samples. **a** Average precision across variant allele frequency and sequencing depth. **b** Average recall across variant allele frequency and sequencing depth. For each tool, the average precision or recall (y-axis) is computed for samples simulated at each sequencing depth, considering variants at some minimum allele frequency (x-axis), averaged over haplotype, depth and replicate. The band around each line represents the 95% confidence interval. Note that the x-axis tick gaps are not proportional to the actual simulated variant AF, and are larger for AF < 10% as this is where the greatest tool-wise differences occur.

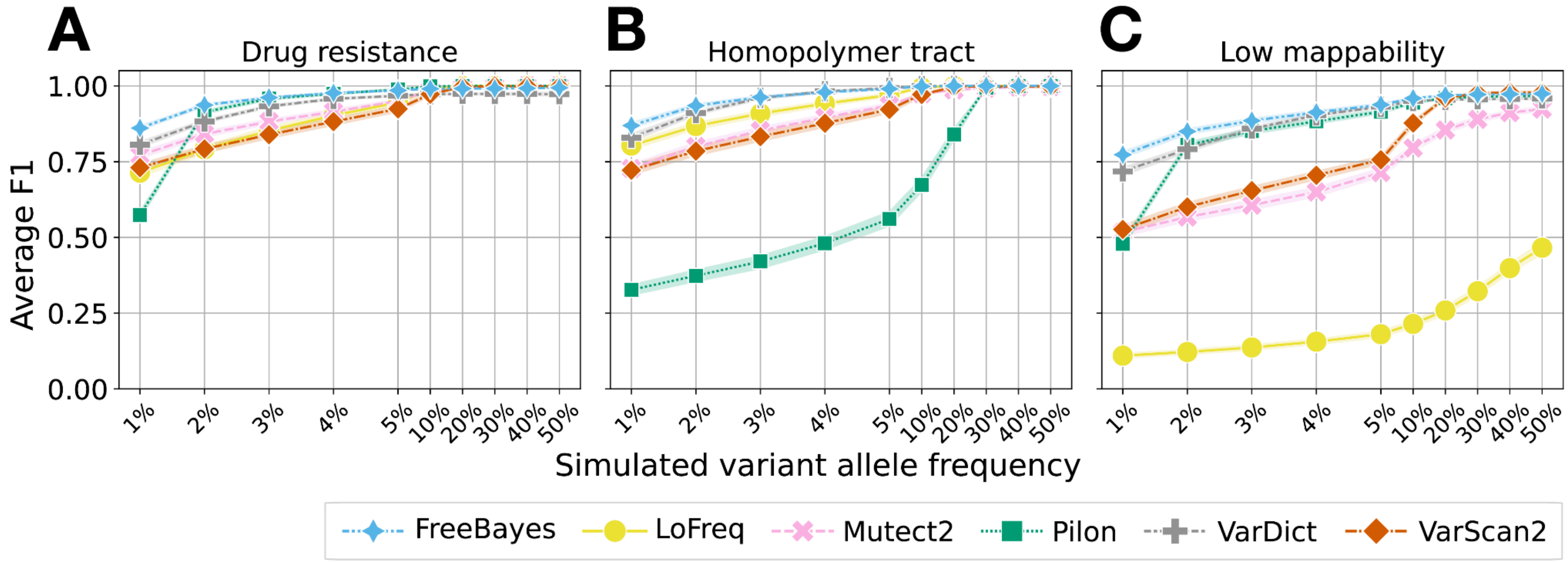


**Figure S19** Average cumulative F1 across variant AF pooled over all simulated depths. **a** Average cumulative F1 in drug resistance regions (H37Rv and L1-4 strains). **b** Average cumulative F1 in homopolymer tract regions (H37Rv strains only). **c** Average cumulative F1 in low mappability regions (H37Rv strains only). To compute the average cumulative F1, we computed cumulative precision and recall as a function of increasing minimum variant AF for each of the six tools, averaged over haplotype, depth and replicate. The band around each line represents the 95% confidence interval. Note that the x-axis tick gaps are not proportional to the actual simulated variant AF, and are larger for AF < 10% as this is where the greatest tool-wise differences occur.

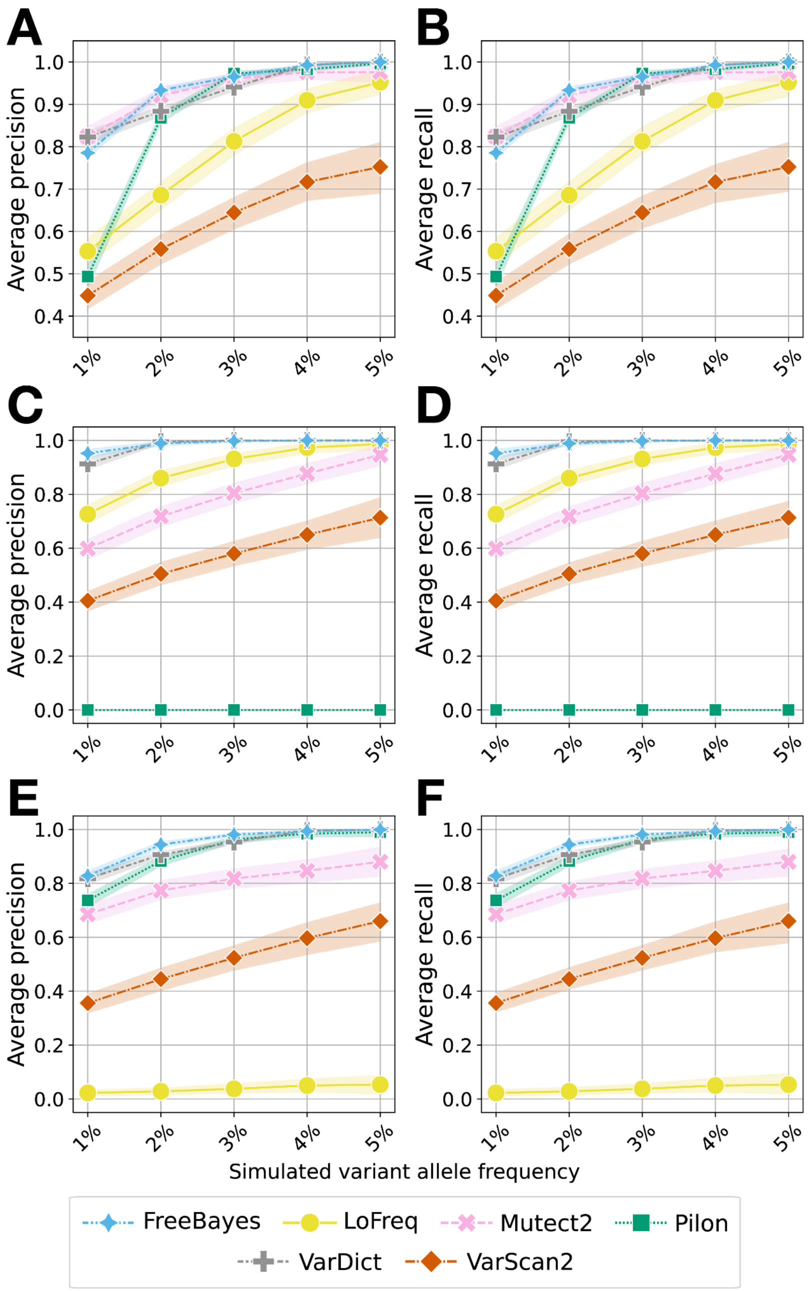


**Figure S20** Average cumulative precision and recall across variant AF ≤ 5% pooled over depths of 50x, 100x and 200x. **a, b** Precision and recall in drug resistance (H37Rv and L1-4 strains). **c, d** Precision and recall in homopolymer tract regions. **e, f** Precision and recall in low mappability regions. We computed cumulative precision and recall as a function of increasing minimum variant AF for each of the six tools, averaged over haplotype, depths 50-200x and replicate, considering only variants detected at AF ≤ 5%. The band around each line represents the 95% confidence interval. Note that the y-axis is from 0.4-1.0 for A and B, and 0-1.0 for C-F. The x-axis tick gaps are not proportional to the actual simulated variant AF, and are larger for AF < 10% as this is where the greatest tool-wise differences occur.

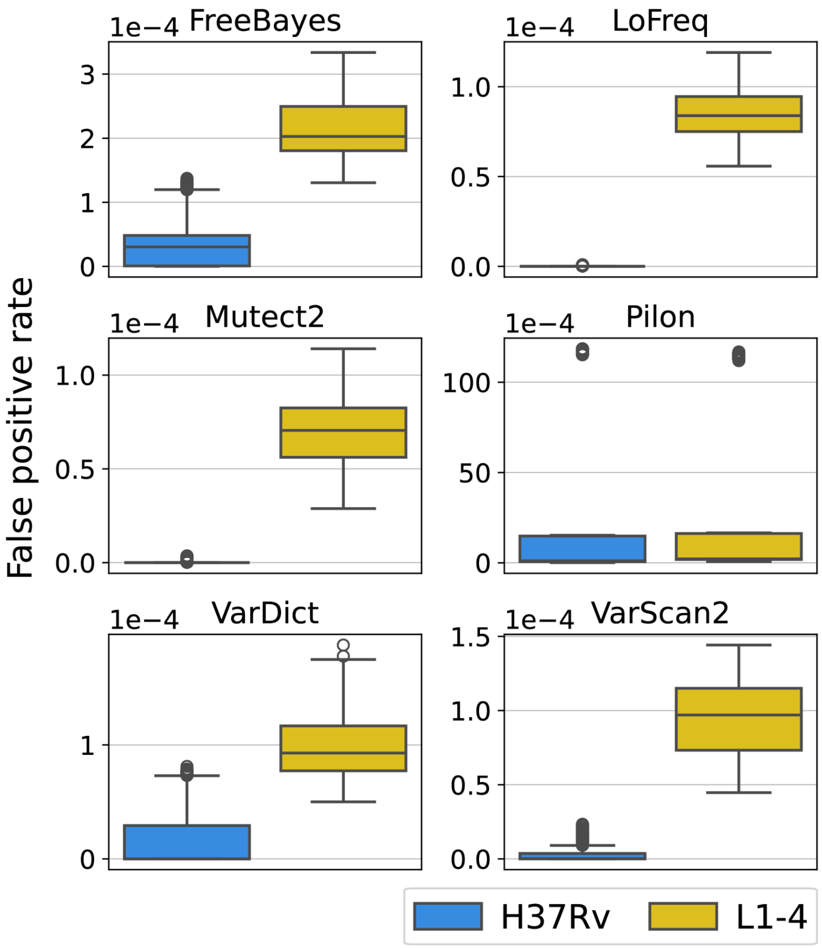


**Figure S21** Genome-wide false positive rate in H37Rv versus L1-4 strains for AF 5-50%. False positive rate is calculated per-base across the entire genome. The average genome-wide FPRs for the FreeBayes, LoFreq, Mutect2, Pilon, VarDict and VarScan2 are 3.96E-05, 4.81E-08, 3.63E-08, 2.67E-03, 1.91E-05 and 2.43E-06 respectively for the H37Rv strains, and 2.15E-04, 8.5E-05, 6.9E-05, 2.71E-03, 9.7E-05 and 9.5E-05 respectively for the L1-4 strains. The pairwise comparisons for each tool are statistically significant after Benjamini-Hochberg correction (Mann-Whitney U test with FDR = 0.05).

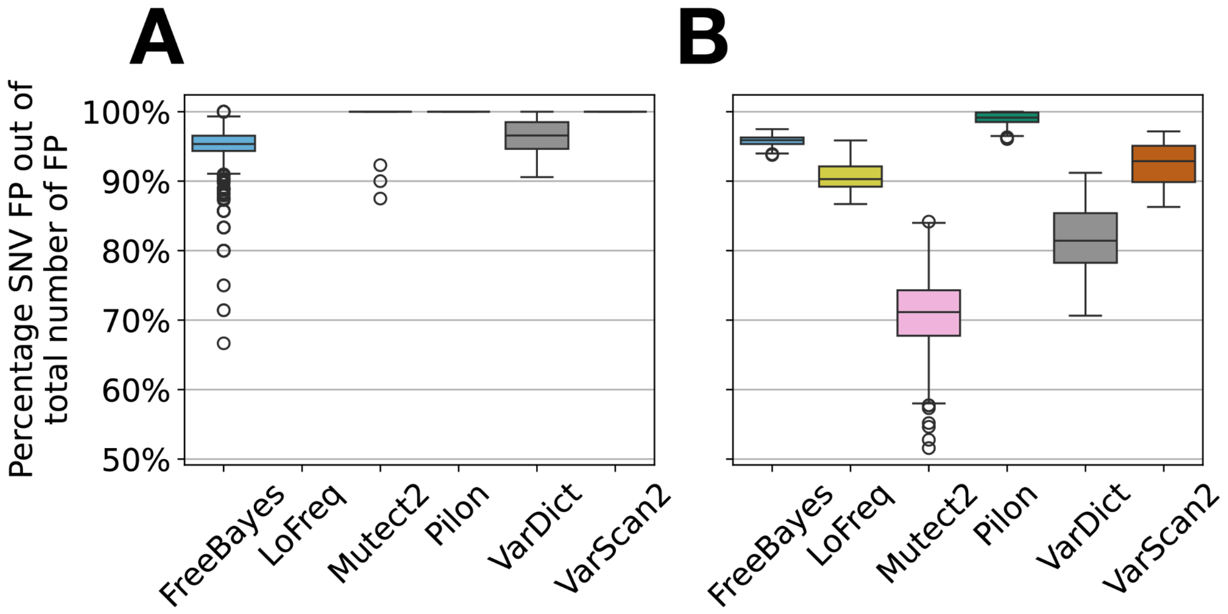


**Figure S22** Distribution of the percentage of FPs that are SNVs per tool and genome background. **a** The distribution of SNV FP percentages per strain in H37Rv strains. **b** The distribution of SNV FP percentages per strain in L1-4 strains. For each tool, the percentage of FPs detected in a single strain that are SNVs was calculated for all strains with at least 5 FPs detected by that tool.

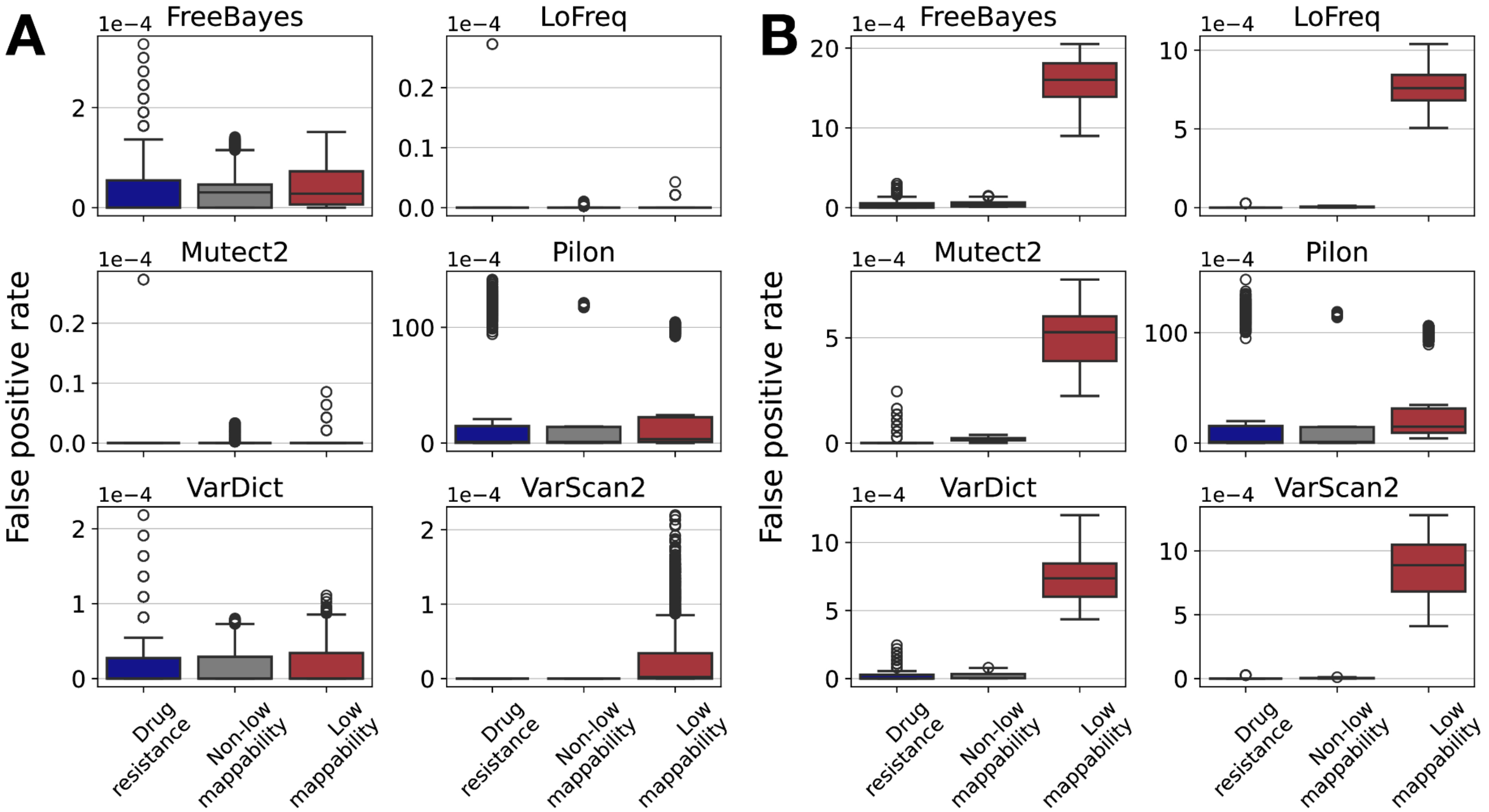


**Figure S23** False positive rates by tool in drug resistance, non-low mappability and low mappability regions. **a** The distribution of false positive rates per strain and region in H37Rv strains. **b** The distribution of false positive rates per strain and region in L1-4 strains. Each region is defined to be mutually exclusive for this comparison i.e. the non-low mappability regions do not include the drug resistance regions. Each box plot shows the distribution of FPRs in each region and for each tool. Note that the subplots do not share the same y-axis. The pairwise comparisons between each region for each tool are statistically significant in both groups of strains after Benjamini-Hochberg correction (Mann-Whitney U test with FDR = 0.05).

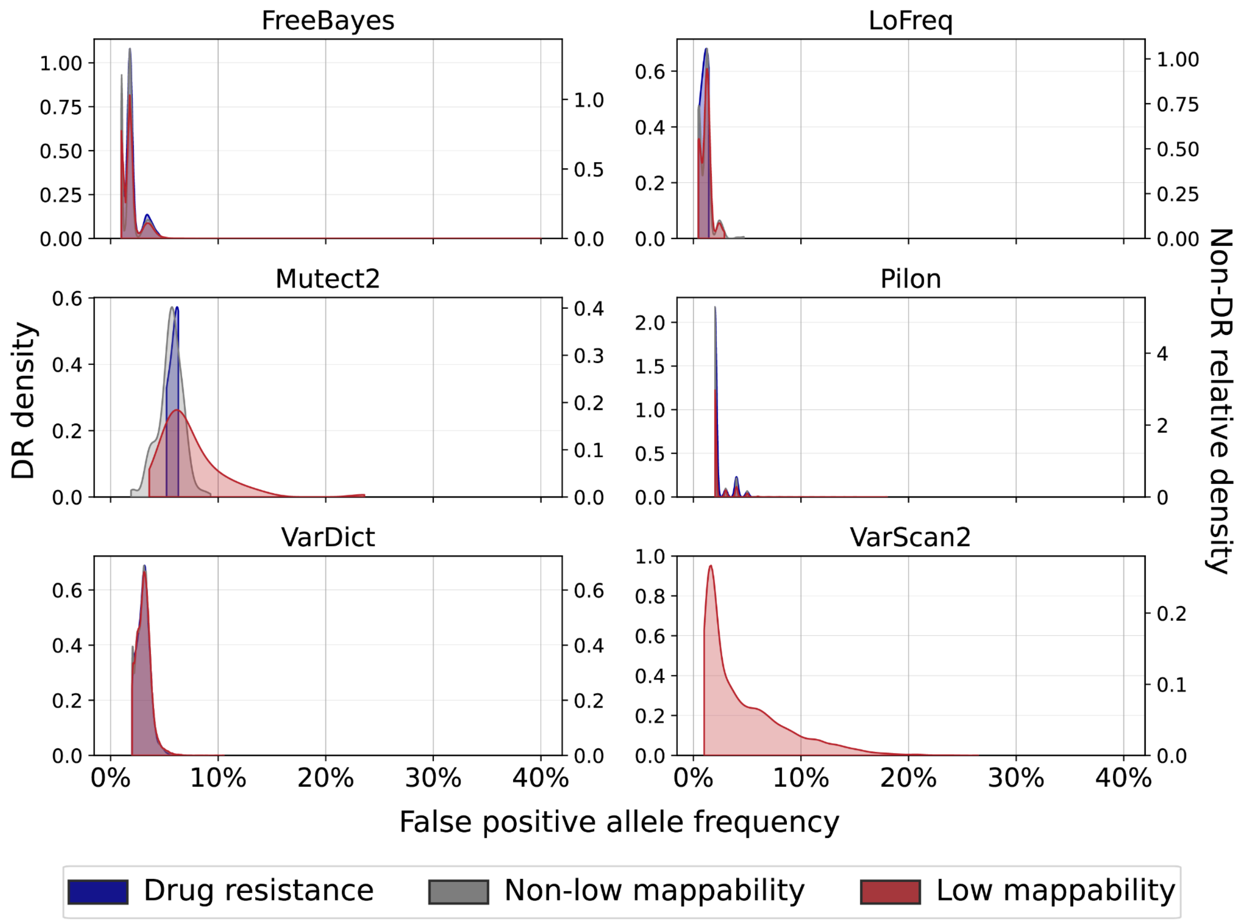


**Figure S24** False positive allele frequency distribution in the H37Rv strains by region. Each region is defined to be mutually exclusive for this comparison i.e. the non-low mappability regions do not include the drug resistance regions. The left y-axis displays the densities of the DR AF distributions, and the right y-axis displays the relative densities of the low mappability (non-DR) and non-low mappability (non-DR) AF distributions (normalized independently). For Pilon we include only the FP with AF > 1% (a median of 99.96% of Pilon FPs across all strains have AF = 1%). All DR FP and non-low mappability FP occur at AF < 7% and AF ≤ 10% respectively. FP in low mappability regions occur at AFs 1-40%.

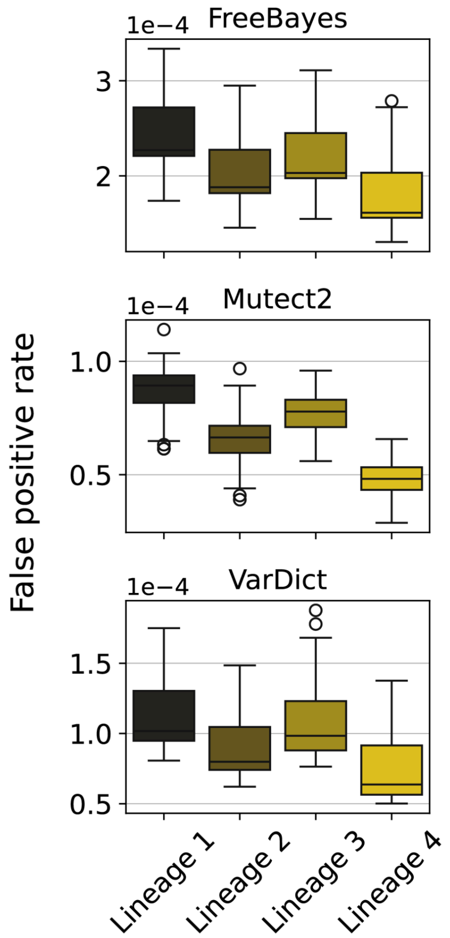


**Figure S25** Genome-wide false positive rate by lineage background (lineages 1-4). False positive rate is calculated per-base across the entire genome. The largest difference between any one pair of lineages for any tool occurs between the FPRs for lineages 1 and 4 in FreeBayes (average FPR is 2.5E-04 and 1.8E-04 respectively).

**Figures S26-30** Distributions of read mapping and quality characteristics at variant sites split by true and false positives in the L1-4 strains for each tool.

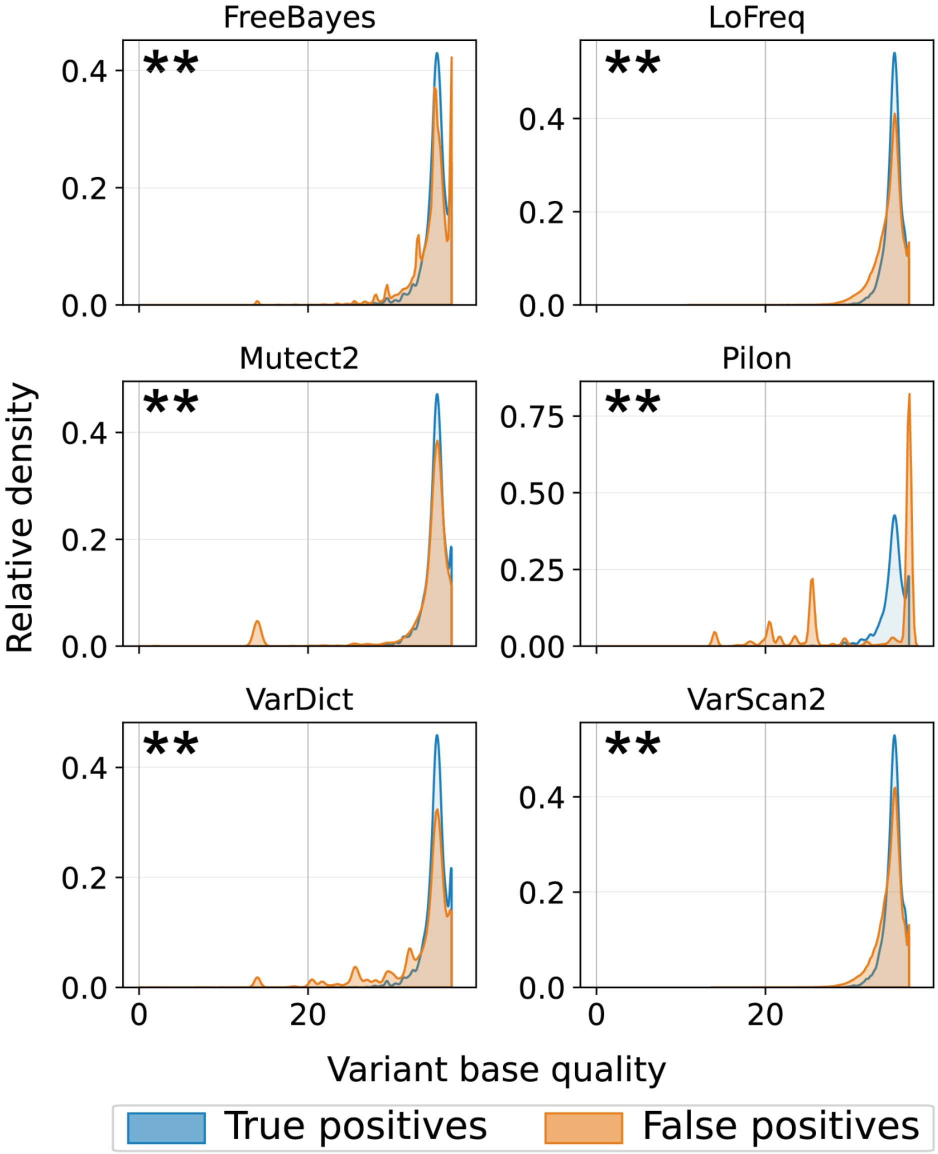


**Figure S26** Variant allele base quality distribution at true and false positive variant sites. The differences between the TP and FP distributions are statistically significant for all tools after Benjamini-Hochberg correction (indicated by asterisks; Mann-Whitney U test with FDR = 0.05).

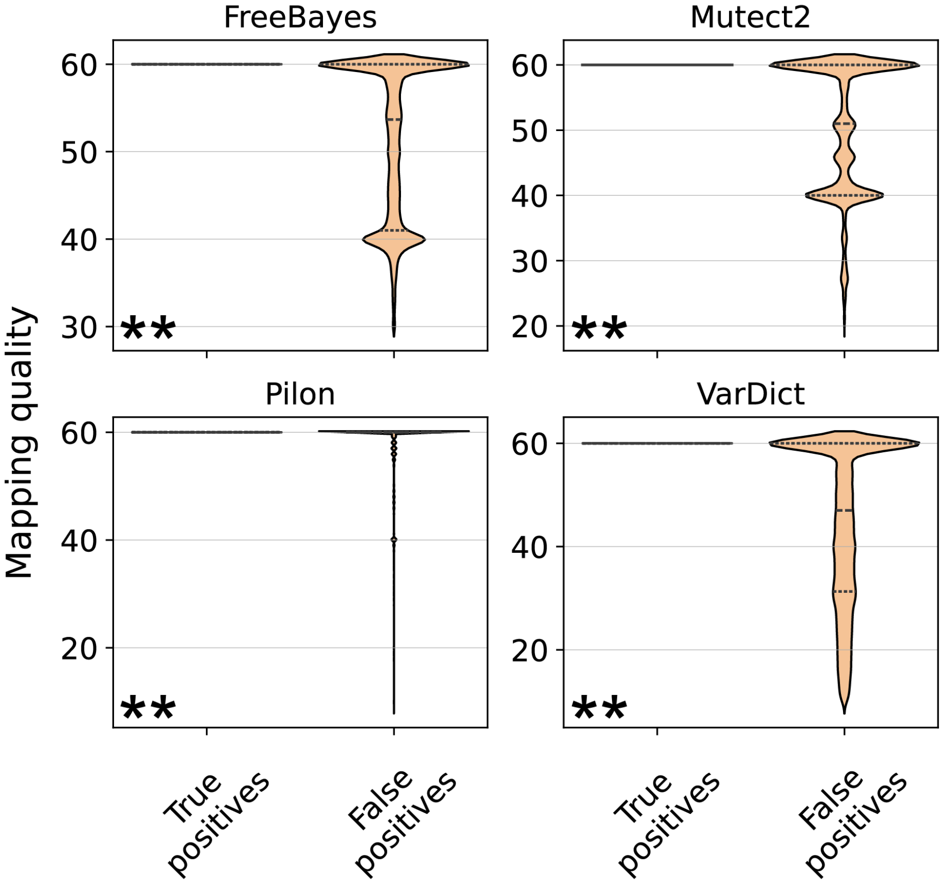


**Figure S27** Mapping quality distribution at true and false positive variant sites. Mapping quality was available for variant calls made by FreeBayes, Mutect2, Pilon and VarDict only (LoFreq and VarScan2 do not provide the mapping quality for variant alleles in their output files). The differences between the TP and FP distributions are statistically significant for all tools after Benjamini-Hochberg correction (indicated by asterisks; Mann-Whitney U test with FDR = 0.05).

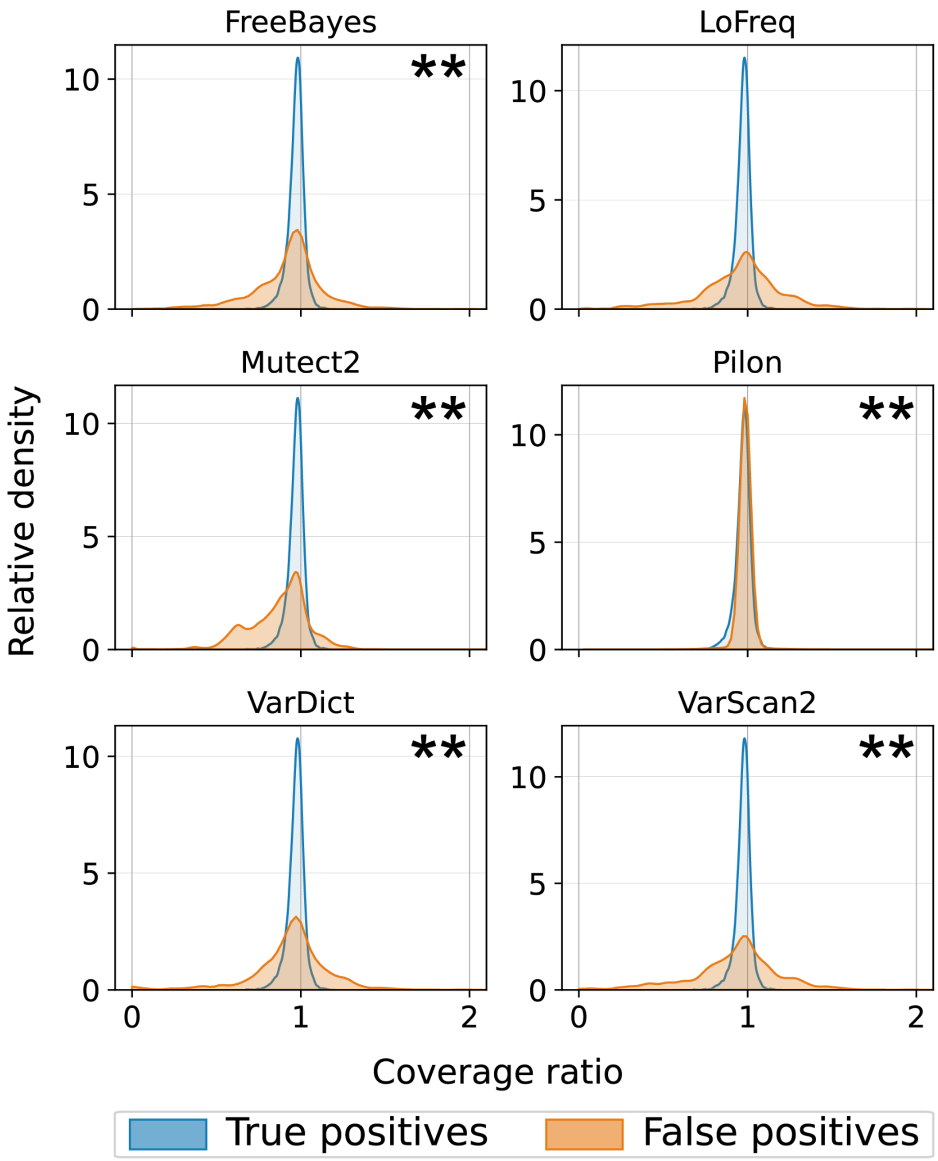


**Figure S28** Coverage ratio distribution at true and false positive variant sites. The coverage ratio is equal to the coverage at a variant site relative to the average regional coverage. The x-axis is cut-off at 2. Across all tools, 0% of TP sites and 0.01-0.22% of FP sites have a coverage ratio > 2. The differences between the TP and FP distributions are statistically significant for all tools except LoFreq after Benjamini-Hochberg correction (indicated by asterisks; Mann-Whitney U test with FDR = 0.05).

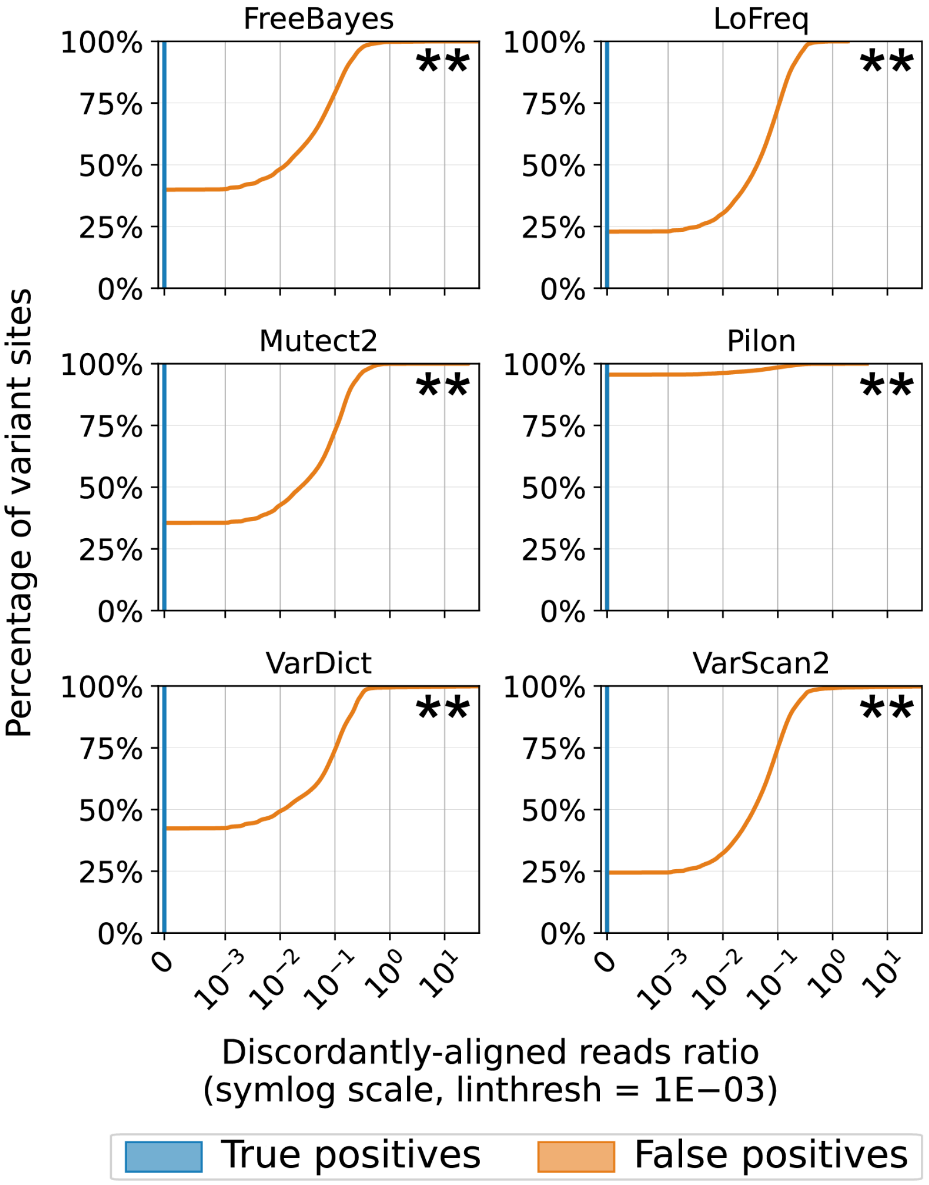


**Figure S29** Discordantly-aligned reads ratio distribution at true and false positive variant sites. The discordantly-aligned read ratio is equal to the number of discordantly-aligned reads at a variant site relative to site coverage. The empirical cumulative distribution function is shown on a symmetric logarithmic x-axis with a linear threshold of 1E-03. Across all tools, 100% of TP sites and 23-96% of FP sites have a discordantly-aligned reads ratio of 0. The ratio distributions for Pilon are the least practically different between TPs and FPs. The differences between the TP and FP distributions are statistically significant for all tools after Benjamini-Hochberg correction (indicated by asterisks; Mann-Whitney U test with FDR = 0.05).

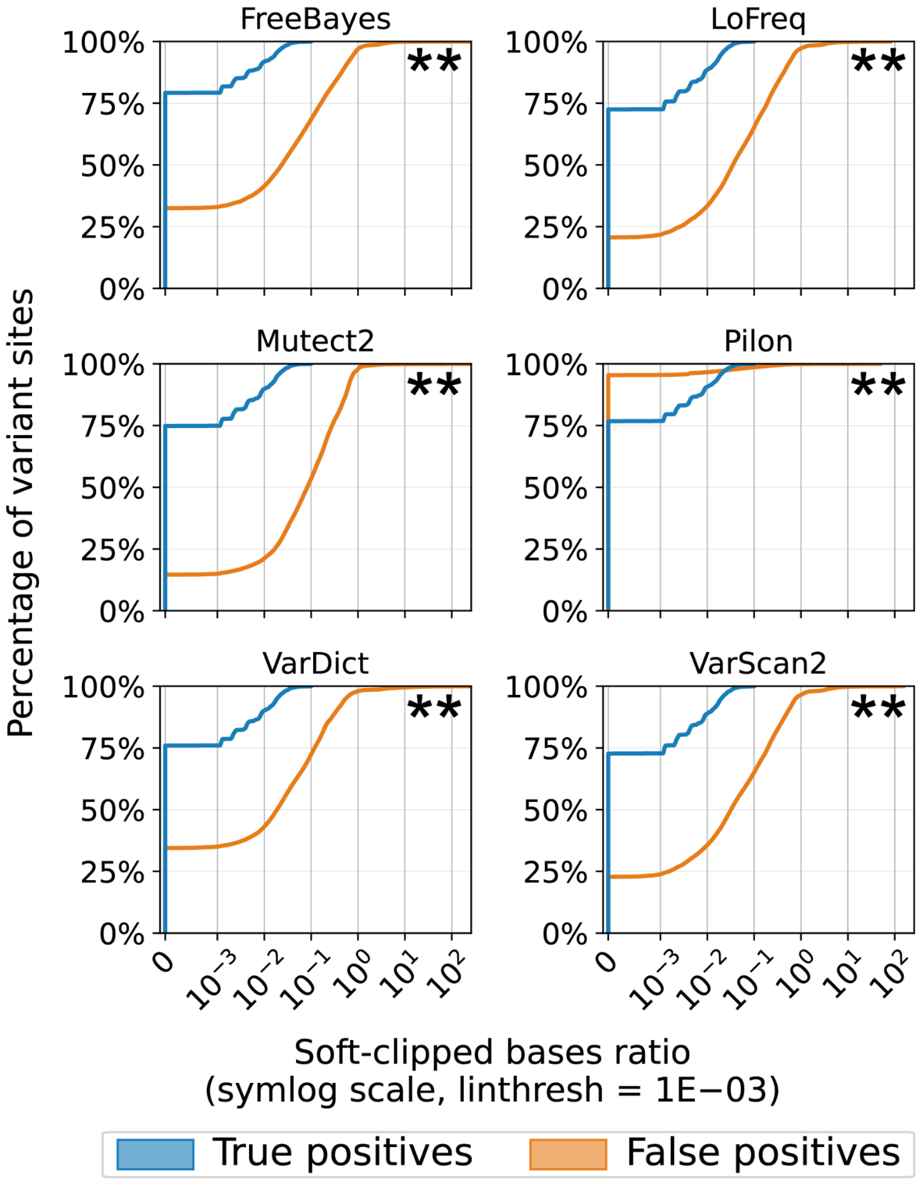


**Figure S30** Soft-clipped bases ratio distribution at true and false positive variant sites. The soft-clipped bases ratio is equal to the number of soft-clipped bases at a variant site relative to site coverage. The empirical cumulative distribution function is shown on a symmetric logarithmic x-axis with a linear threshold of 1e-3. Across all tools, 72-79% of TP sites and 15-95% of FP sites have a soft-clipped bases ratio of 0. Pilon is the only tool to exhibit a higher percentage of false positive calls at variant sites with a soft-clipped bases ratio of 0, which is likely due to the fact that a median of 42% of the FPs called by Pilon per L1-4 strain occur at AF = 1%. The differences between the TP and FP distributions are statistically significant for all tools after Benjamini-Hochberg correction (indicated by asterisks; Mann-Whitney U test with FDR = 0.05).

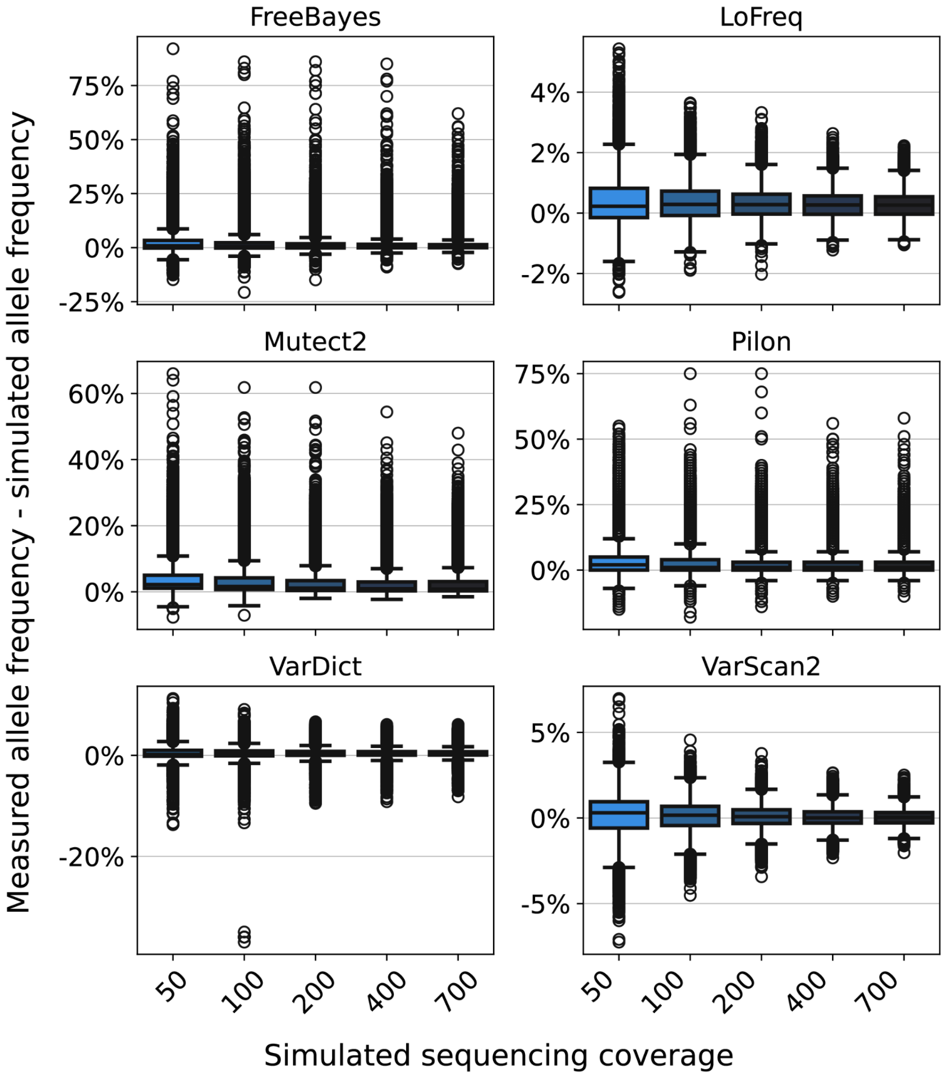


**Figure S31** Distribution of the difference between the measured allele frequency and the simulated allele frequency for variants simulated at each sequencing depth. While the differences between some pairwise depth comparisons are statistically significant after Benjamini-Hochberg correction (Mann-Whitney U test with FDR = 0.05), the maximum average difference of the measured AF and simulated AF for any pair of depths and any tool is -1.57%. Note that the subplots do not share the same y-axis.

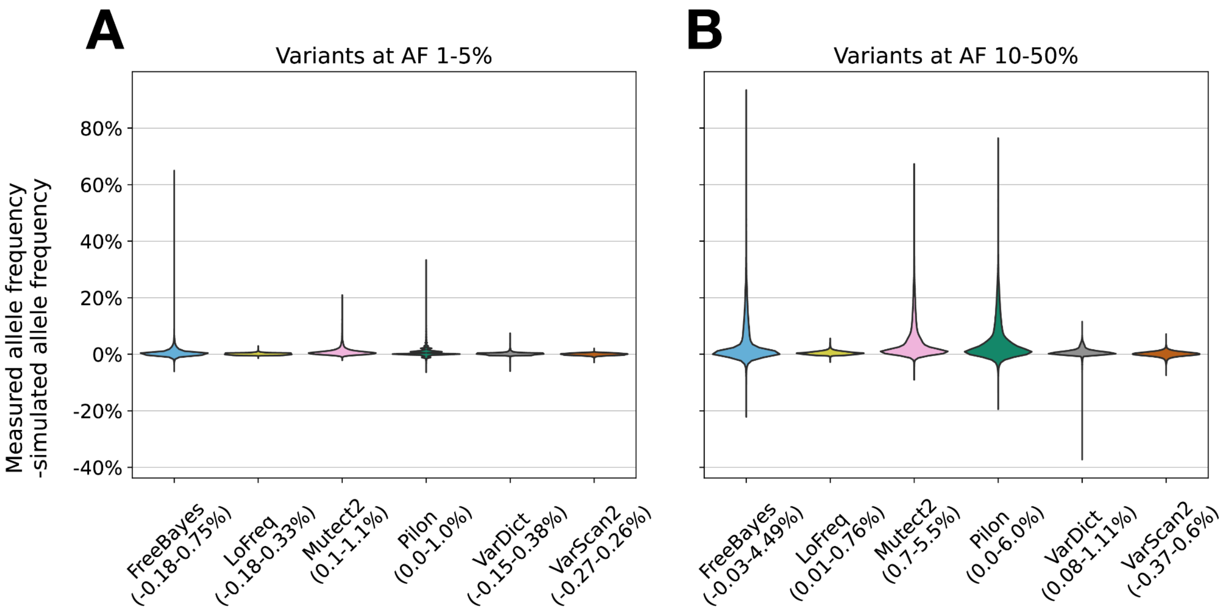


**Figure S32** Distribution of the difference between the measured and simulated allele frequency for variants across all variants, depths, haplotypes and replicates. **a** Distribution for variants simulated at 1% ≤ AF ≤ 5%. **b** Distribution for variants simulated at 10% ≤ AF ≤ 50%. The IQR for each distribution is displayed next to the tool name. Overall, as expected, the highest AF differences are observed for variants simulated at an AF in the higher range (10-50%).

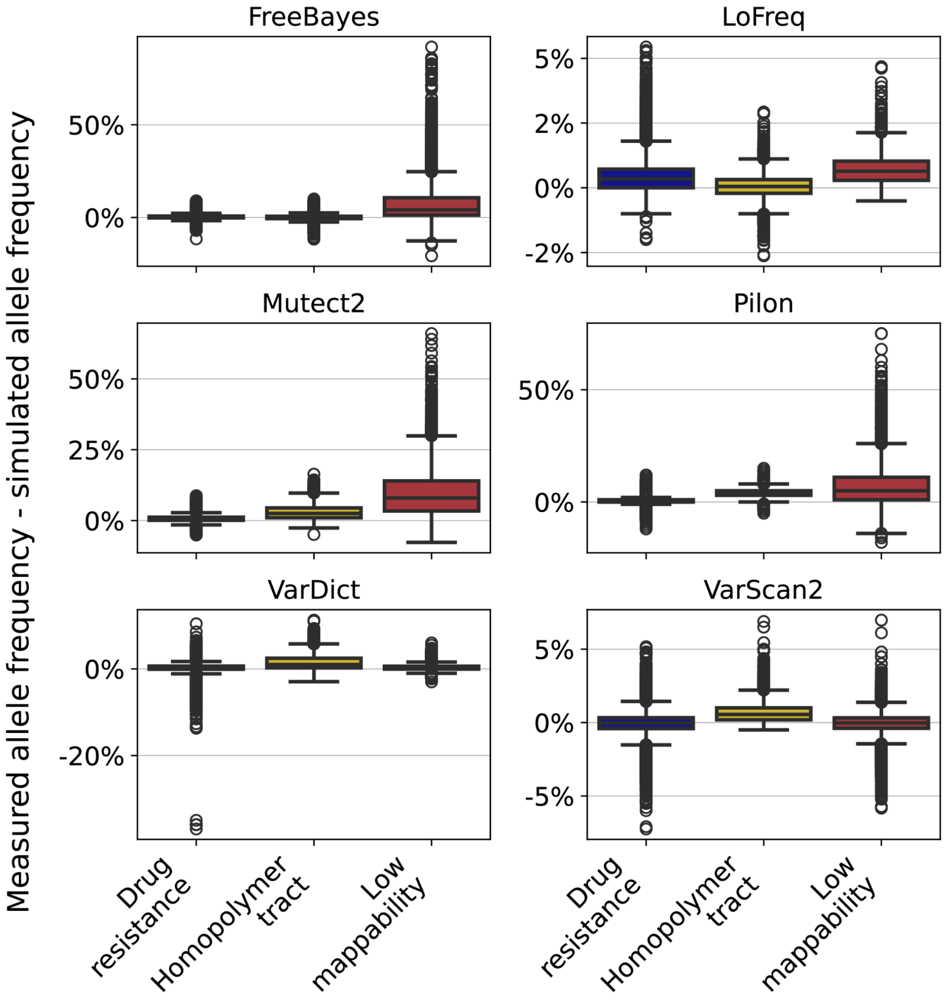


**Figure S33** Distribution of the difference between the measured allele frequency and the simulated allele frequency for variants simulated in each mutation region. All pairwise comparisons are statistically significant after Benjamini-Hochberg correction (Mann-Whitney U test with FDR = 0.05), except the DR-LM comparisons for VarDict and VarScan2. The average difference in measured and simulated allele frequency across all tools, however, is not substantially different between DR and HT regions for all tools except Mutect2 and Pilon (mean AF difference for Mutect2 and Pilon: DR = 0.58%, HT = 3.33%; mean AF difference for other tools: DR = 0.24%, HT = 0.54%). The AFs of LM variants measured by FreeBayes, Mutect2 and Pilon are practically the most different from those in any other region measured by these tools, or those measured in all other regions by any other tool (mean AF difference: FreeBayes, Mutect2 and Pilon LM = 7.91%; FreeBayes, Mutect2 and Pilon DR and HT = 1.36%, LoFreq, VarDict and VarScan2 all regions = 0.42%). Note that the subplots do not share the same y-axis.

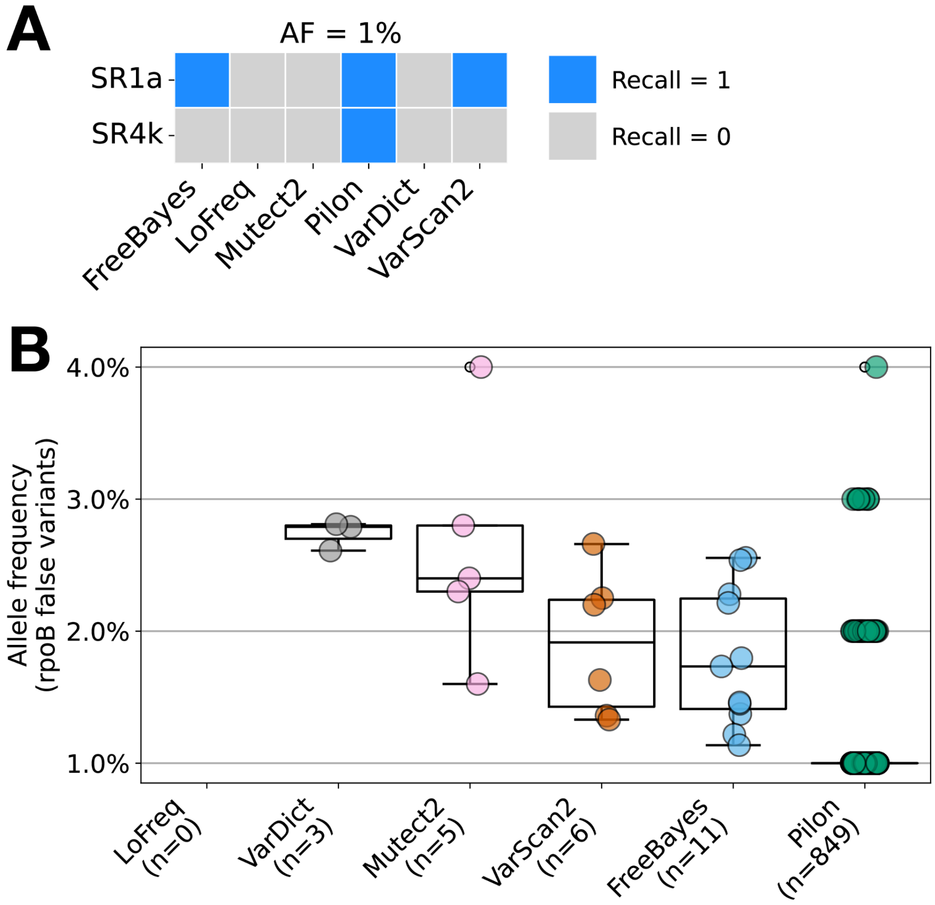


**Figure S34** Variant caller performance in in-vitro samples with introduced *rpoB* mutations across all allele frequencies (*rpoB* only). **a** Heatmap of recall for each tool on each in-vitro isolate with mutations introduced at AF = 1%. A Ser531Leu mutation was introduced in the SR1a isolates; a His526Pro mutation was introduced in the SR4k isolates. **b** Distribution of the false variants AFs detected in *rpoB* by each tool. The number of total false variants detected by a tool is shown in parentheses below each tool name.

#### Supplementary Tables

**Table S1** Average difference between the simulated AF and the expected AF in each region for ISS and ART.

| | Simulated AF – excepted AF | | | | | |
|---|---|---|---|---|---|---|
| | ISS | | | ART | | |
| | **DR** | **HT** | **LM** | **DR** | **HT** | **LM** |
| **AF < 10%** | -0.012% | -0.352% | -0.963% | 0.006% | -0.301% | -1.825% |
| **AF ≥ 10%** | -0.652% | -3.443% | -10.285% | -0.413% | -3.170% | -18.237% |

This table describes the average difference between the simulated AF and the expected AF in drug resistance (DR), homopolymer tract (HT) and low mappability (LM) regions. The average difference for DR regions includes both H37Rv and L1-4 strains. The largest average differences between the simulated and expected AFs occur in LM regions.

**Table S2** Median number of low-frequency variants found in DR, HT, LM and all other regions across 38 clinical samples.

| Genomic region | Median number of low-frequency variants |
|---|---|
| Drug resistance | 0 |
| Homopolymer tract | 1 |
| Low mappability | 4 |
| Other | 106 |

The number of low-frequency variants per strain did not differ by (1) lineage, or (2) average sequencing coverage after multiple Mann-Whitney U tests (Benjamini-Hochberg corrected, FDR = 0.05).

**Table S3** Average DR-F1 per tool pooled across strains from an H37Rv and L1-4 background.

| Tool | Average pooled F1 | Average L1-4 F1 – H37Rv F1 |
|---|---|---|
| FreeBayes | 0.86 | -0.038** |
| VarDict | 0.81 | 0.005 |
| Mutect2 | 0.77 | 0.065** |
| VarScan2 | 0.73 | -0.003 |
| LoFreq | 0.71 | -0.005 |
| Pilon | 0.57 | -0.003 |

The asterisks indicate the differences that were statistically significant after Benjamini-Hochberg correction (Mann-Whitney U test with FDR = 0.05).

The average F1 in drug resistance regions pooled across all genomic backgrounds is displayed in the first column. The average difference in average DR F1 score (L1-4 – H37Rv) is displayed in the second column.

**Table S4** Difference in average DR F1 score for pairwise comparisons between each lineage (L1-4) for each tool.

| | Average DR F1 difference | | | | | |
|---|---|---|---|---|---|---|
| **Tool** | **L2 – L1** | **L3 – L1** | **L4 – L1** | **L3 – L2** | **L4 – L2** | **L4 – L3** |
| FreeBayes | -0.014 | -0.024** | -0.004 | -0.010** | 0.010 | 0.020** |
| LoFreq | -0.008 | -0.009 | -0.005 | -0.001 | 0.003 | 0.004 |
| Mutect2 | -0.008 | 0.004 | 0.011 | 0.012 | 0.019 | 0.007 |
| Pilon | -0.006 | 0.001 | 0.002 | 0.007 | 0.008 | 0.001 |
| VarDict | -0.013 | -0.046** | 0.005 | -0.033** | 0.018 | 0.051** |
| VarScan2 | -0.011 | -0.014 | -0.013 | -0.003 | -0.002 | 0.001 |

The asterisks indicate the differences that were statistically significant after Benjamini-Hochberg correction (Mann-Whitney U test with FDR = 0.05).

The value in each cell is the average DR F1 score difference between each pair of lineages, e.g. L2 – L1 indicates the average DR F1 score difference between lineage 2 and lineage 1 strains. The only pairwise comparisons that are statistically significant are those involving L3 for FreeBayes and VarDict. We discuss the issue these tools had with consistently missed mutations in L3 in Supplementary Results C. The P-values for all other comparisons are > 0.05.

**Table S5** Average F1 score by tool and genomic region at depths 50-200x for increasing minimum variant allele frequency 1-50%.

| | F1 score | | | | | | | | | | | | | | | | | |
|---|---|---|---|---|---|---|---|---|---|---|---|---|---|---|---|---|---|---|
| | Drug resistance | | | | | | Homopolymer tract | | | | | | Low mappability | | | | | |
| Minimum variant AF | FB | LF | MT | PL | VD | VS | FB | LF | MT | PL | VD | VS | FB | LF | MT | PL | VD | VS |
| 1% | 0.80 | 0.61 | 0.72 | 0.51 | 0.78 | 0.59 | 0.83 | 0.71 | 0.63 | 0.33 | 0.82 | 0.58 | 0.70 | 0.09 | 0.46 | 0.63 | 0.70 | 0.44 |
| 2% | 0.90 | 0.68 | 0.79 | 0.86 | 0.85 | 0.66 | 0.90 | 0.79 | 0.70 | 0.38 | 0.89 | 0.65 | 0.78 | 0.10 | 0.51 | 0.71 | 0.77 | 0.49 |
| 3% | 0.94 | 0.76 | 0.85 | 0.93 | 0.91 | 0.73 | 0.94 | 0.85 | 0.77 | 0.43 | 0.94 | 0.72 | 0.83 | 0.11 | 0.56 | 0.78 | 0.83 | 0.54 |
| 4% | 0.97 | 0.84 | 0.90 | 0.96 | 0.95 | 0.80 | 0.97 | 0.90 | 0.84 | 0.49 | 0.97 | 0.79 | 0.87 | 0.13 | 0.61 | 0.83 | 0.88 | 0.61 |
| 5% | 0.98 | 0.91 | 0.94 | 0.98 | 0.96 | 0.87 | 0.98 | 0.95 | 0.90 | 0.57 | 0.99 | 0.87 | 0.91 | 0.15 | 0.67 | 0.88 | 0.92 | 0.69 |
| 10% | 0.99 | 0.98 | 0.98 | 1.00 | 0.97 | 0.95 | 1.00 | 0.99 | 0.96 | 0.69 | 1.00 | 0.95 | 0.95 | 0.17 | 0.73 | 0.93 | 0.95 | 0.84 |
| 20% | 0.99 | 1.00 | 0.99 | 1.00 | 0.97 | 0.99 | 1.00 | 1.00 | 0.98 | 0.86 | 1.00 | 0.99 | 0.96 | 0.21 | 0.80 | 0.95 | 0.95 | 0.94 |
| 30% | 0.99 | 1.00 | 1.00 | 1.00 | 0.97 | 1.00 | 1.00 | 1.00 | 0.99 | 0.99 | 1.00 | 1.00 | 0.97 | 0.26 | 0.86 | 0.96 | 0.95 | 0.97 |
| 40% | 0.99 | 1.00 | 1.00 | 1.00 | 0.97 | 1.00 | 1.00 | 1.00 | 0.99 | 1.00 | 1.00 | 1.00 | 0.97 | 0.33 | 0.89 | 0.96 | 0.96 | 0.97 |
| 50% | 0.99 | 1.00 | 1.00 | 1.00 | 0.97 | 1.00 | 1.00 | 1.00 | 1.00 | 1.00 | 1.00 | 1.00 | 0.97 | 0.41 | 0.91 | 0.96 | 0.96 | 0.97 |

FB = FreeBayes; LF = LoFreq; MT = Mutect2; PL = Pilon; VD = VarDict; VS = VarScan2

The value in each cell is the average F1 score for each tool in a specific genomic region, enforcing an increasing minimum variant allele frequency (AF) in each row. The F1 scores in drug resistance regions are averaged across strains from both an H37Rv and non-H37Rv background, while the F1 scores in the homopolymer tract and low mappability regions are for strains from an H37Rv background only.

**Table S6** Median FPR across H37Rv and L1-4 strains per tool and overall by region.

| | False positive rate | | |
|---|---|---|---|
| **Tool** | **Drug resistance** | **Non-low mappability** | **Low mappability** |
| FreeBayes | 0.00 | 3.23E-05 | 6.40E-05 |
| LoFreq | 0.00 | 0.00 | 0.00 |
| Mutect2 | 0.00 | 0.00 | 0.00 |
| Pilon | 1.09E-04 | 1.11E-04 | 9.40E-04 |
| VarDict | 0.00 | 2.56E-06 | 2.77E-05 |
| VarScan2 | 0.00 | 0.00 | 2.35E-05 |
| Overall | 0.00 | 1.54E-06 | 4.10E-06 |

This table describes the false positive rate (FPR) in drug resistance, non-low mappability and low mappability regions. Each region is defined to be mutually exclusive for this comparison i.e. the non-low mappability regions do not include the drug resistance regions. LoFreq and Mutect2 achieve the lowest median FPR across all strains. The median FPR is lowest in drug resistance resistance and highest in low mappability regions for all tools.

**Table S7** Percentage of FP detected in LM regions and at AF = 1% by each tool for the H37Rv simulations and L1-4 simulations separately.

| | Median number of FP | | Median % LM FP | | Median % FP AF = 1% | |
|---|---|---|---|---|---|---|
| **Tool** | **H37Rv** | **L1-4** | **H37Rv** | **L1-4** | **H37Rv** | **L1-4** |
| Pilon | 481 | 968 | 10.91 | 53.14 | 99.96 | 41.72 |
| FreeBayes | 141 | 894 | 13.54 | 79.64 | 0.00 | 0.00 |
| VarDict | 130 | 410 | 10.00 | 95.26 | 0.00 | 0.00 |
| VarScan2 | 13 | 428 | 100.00 | 97.40 | 0.00 | 0.00 |
| LoFreq | 1 | 370 | 0.00 | 95.81 | 0.00 | 0.00 |
| Mutect2 | 2 | 311 | 0.00 | 76.13 | 0.00 | 0.00 |

For each tool we list the median total number of genome-wide FP detected at all variant AFs in H37Rv and L1-4 separately (first two columns), the percentage of these total FP that occur in low mappability regions (second two columns), and the percentage of these FP total FP that occur at AF = 1%. The tools are sorted in descending order by the median total number of FP detected (H37Rv + L1-4).

**Table S8** SNV filtering (only hard filtering or region masking) of false and true variants called by FreeBayes (AF 5-50%) in L1-4 strains.

| | | | **Percentage filtered (%)** | | | |
|---|---|---|---|---|---|---|
| | **Total number pre-filtering** | | **Hard filtering** | | **Region masking** | |
| **Sequencing coverage** | **FP** | **TP** | **FP** | **TP** | **FP** | **TP**[a] |
| 50x | 55 492 | 2 012 | 46.1 | 20.4 | 94.9 | 16.1 |
| 100x | 55 366 | 2 017 | 36.0 | 3.8 | 95.7 | 15.7 |
| 200x | 54 980 | 2 063 | 29.2 | 0.1 | 95.8 | 15.0 |
| 400x | 54 813 | 2 045 | 26.4 | 0.0 | 96.0 | 15.9 |
| 700x | 53 920 | 2 029 | 25.7 | 0.0 | 96.3 | 14.7 |
| Overall | 274 571 | 10 166 | 32.7 | 4.9 | 95.7 | 15.5 |

[a]Three of the 20 variants simulated across all L1-4 genomic backgrounds are within the regions masked by the region masking filtering scheme.

The pre-filtering group of columns displays the total number of FP and TP SNVs called by FreeBayes across all strains simulated at a specific sequencing depth. A total of 120 strains are considered in each sequencing depth group (six variant AFs 5-50%, four background genomes, five replicates). Hard filtering excludes variants with forward and reverse strand allele counts < 2, depth < 5, mapping quality < 40, and region masking excludes variants in low mappability regions and rRNA genes. FP and TP statistics are broken down by simulated sequencing coverage group, and summarized overall.

**Table S9** SNV filtering of false and true variants called by FreeBayes (AF < 5%) in L1-4 strains.

| | Total number pre-filtering | | Percentage filtered (%) | | | | | | | | | |
|---|---|---|---|---|---|---|---|---|---|---|---|---|
| | | | F1: Error model filtering | | F2: Error model + hard filtering | | F3: Error model + region masking | | F4: Hard filtering + region masking | | F5: Error model + hard filtering + region masking | |
| Sequencing coverage | FP | TP | FP | TP | FP | TP | FP | TP[a] | FP | TP[a] | FP | TP[a] |
| 50x | 18 618 | 337 | 30.0 | 0.4 | 100.0 | 100.0 | 49.9 | 17.2 | 100.0 | 100.0 | 100.0 | 100.0 |
| 100x | 60 338 | 815 | 23.3 | 0.1 | 99.6 | 85.8 | 39.0 | 15.0 | 99.8 | 87.9 | 100.0 | 86.9 |
| 200x | 42 259 | 1 049 | 43.1 | 0.4 | 97.8 | 45.0 | 65.1 | 15.5 | 98.4 | 52.5 | 99.7 | 47.4 |
| 400x | 28 625 | 1 212 | 65.8 | 0.0 | 94.7 | 8.8 | 95.6 | 14.7 | 95.0 | 22.9 | 99.2 | 14.4 |
| 700x | 29 253 | 1 261 | 65.3 | 0.0 | 92.2 | 0.9 | 95.6 | 15.4 | 93.8 | 16.3 | 99.0 | 6.4 |
| Overall | 179 093 | 4 674 | 45.5 | 0.7 | 96.9 | 46.8 | 69.0 | 15.5 | 97.4 | 54.8 | 99.6 | 54.8 |

[a]Three of the 20 variants simulated across all L1-4 genomic backgrounds are within the regions masked by the region masking filtering scheme.

The pre-filtering group of columns displays the total number of FP and TP SNVs called by FreeBayes across all strains simulated at a specific sequencing depth. A total of 80 strains are considered in each sequencing depth group (four variant AFs 1-4%, four background genomes, five replicates). The F1-F5 groups of columns correspond to each of the three sequential SNV filtering schemes and display the average percentages of FPs and TPs filtered out per strain. Error model filtering excludes variants with a model predicted probability ≤ 0.46; hard filtering excludes variants with forward and reverse strand allele counts < 2, depth < 5, mapping quality < 40); region masking excludes variants in low mappability regions and rRNA genes. FP and TP statistics are broken down by simulated sequencing coverage group, and summarized overall.

**Table S10** INDEL filtering of false and true variants called by FreeBayes (AF 5-50%) in L1-4 and H37Rv strains respectively.

| Sequencing coverage | Total number pre-filtering | | Percentage filtered (%) F1: AF adjustment | | F2: AF adjustment + hard filtering | | F3: AF adjustment + hard filtering + region masking | |
|---|---|---|---|---|---|---|---|---|
| | FP | TP | FP | TP | FP | TP | FP | TP |
| 50x | 2 527 | 2 616 | 45.4 | 1.6 | 61.1 | 23.9 | 98.8 | 23.9 |
| 100x | 2 698 | 2 624 | 45.1 | 2.1 | 57.6 | 6.7 | 98.8 | 6.7 |
| 200x | 2 754 | 2 636 | 43.1 | 1.0 | 53.8 | 1.4 | 98.9 | 1.4 |
| 400x | 2 800 | 2 603 | 41.5 | 0.4 | 52.0 | 0.4 | 98.9 | 0.4 |
| 700x | 2 802 | 2 582 | 39.9 | 0.0 | 49.3 | 0.0 | 99.0 | 0.0 |
| Overall | 13 581 | 13 061 | 43.0 | 1.0 | 54.8 | 6.5 | 98.9 | 6.5 |

The pre-filtering group of columns displays the total number of FP and TP INDELs called by FreeBayes across all strains simulated at a specific sequencing depth. A total of 120 L1-4 strains are considered in each sequencing depth group for the FP statistics (six variant AFs 5-50%, four background genomes, five replicates), and a total of 300 H37Rv strains are considered in each sequencing depth group for the TP statistics (six variant AFs 5-50%, ten haplotypes, five replicates). The F1-F3 groups of columns correspond to each of the three sequential INDEL filtering schemes and display the average percentages of FPs and TPs filtered out per strain. F1 (Filter 1): filtering after AF adjustment, F2: filtering after AF adjustment and hard filtering, F3: filtering after AF adjustment, hard filtering and region masking (low mappability regions, rRNA genes and sites within 100bp of an insertion sequence or phage). FP and TP statistics are broken down by simulated sequencing coverage group, and summarized overall. Note for F3 that we simulated variants in the regions masked in this filtering scheme but these variants were not detected by FreeBayes in most strains in the first place.
